# QTL analysis of essential oil, seed shattering, thousand-grain weight and stalk attachment rate in *Carum carvi*

**DOI:** 10.64898/2026.10.01.755638

**Authors:** Daniel von Maydell, Jolien Zimmer, Akhil Pinnuri, Fang-Shiang Lim, Martin Junghanns, Frank Marthe

## Abstract

Caraway (*Carum carvi* L., 2n = 2x = 20) is a medicinal and aromatic plant of the Apiaceae family valued for its essential oil with the major terpenoids carvone and limonene. While the essential oil content represents an important quality trait, low seed shattering and high thousand-grain weight (TGW) are key determinants of seed yield. As molecular tools remain scarce within Apiaceae breeding beyond carrot, we conducted a QTL mapping for these seed-related traits using two previously established F populations (Effective N(P1) = 236; N(P2) = 266). In P1 with the non-shattering maternal parent ‘Rekord’, QTL mapping revealed at least one shattering locus on LG03 (PVE: 9.9%). In addition, loci associated with essential oil (LG08, 10.5% PVE) and TGW (LG09, 11.1% PVE) were detected. In P2, we identified six essential-oil loci (LG02–06, LG10; combined PVE: 49.2%), with five loci affecting both carvone and limonene, and one (LG03) exclusively limonene. Additionally, a major QTL for TGW mapped to LG02 (PVE: 15.6%). For stalk attachment rate one minor QTL was detected (LG07, 8.5% PVE), while no QTL co-localized with the major shattering locus. To pave the path towards fine-mapping, we designed and validated 26 diagnostic PACE markers covering five key QTLs.

## Introduction

Caraway (*Carum carvi* L., 2n = 2x = 20) is a major essential oil crop of the Apiaceae family valued for its essential oil rich in major terpenoids carvone and limonene (Németh, 1998; Pank, 2012). It is used for various pharmaceutical, culinary and industrial purposes (Mahboubi, 2019). Currently, its cultivation is endangered by changing climatic conditions, urging adaptation through breeding. However, due to the necessity to simultaneously select for various agronomic and quality traits, breeding can be a challenging process, particularly in crops with limited resources. Therefore, genomic tools such as marker-assisted selection (MAS) can greatly facilitate effective breeding. However, such tools are largely lacking in caraway breeding (Misginna et al., 2026).

So far, several studies have characterized cultivated and wild derived germplasm on a phenotypic and chemotypic level (Ferrie et al., 2011; Raal et al., 2012; von Maydell et al., 2024; Singh et al., 2026; von Maydell et al., 2026a; Hazrati et al., 2026) or analyzed population structure and genetic diversity using RAPD (Laribi et al., 2011; Seidler-Lozykowska et al., 2014) or SNP markers (von Maydell et al., 2021a; de Haro Reyes et al., 2025), providing basic resources for breeding. In contrast, QTL or association studies linking genetic data to phenotypes are scarce. Bocianowski & Seidler-Lozykowska (2012) found some associations between RAPD markers and several traits based on regression analysis of data from 22 caraway genotypes. However, to our knowledge these markers were not utilized in breeding. The low reproducibility of RAPD markers (Grover & Sharma, 2016), in combination with a low number of genotypes and low marker density could be possible reasons. More recently, a draft genome assembly for caraway with a structural and functional gene annotation was established as basic genomic framework (von Maydell et al., 2026c). In parallel, potential QTLs for flowering time and flowering color were detected by utilizing SNPs from genotyping by sequencing (GBS). Despite these advances, genetic mapping studies for the most economically critical seed traits remain lacking.

This represents a major bottleneck for caraway breeding, as the total essential oil content and the proportion of carvone represent important quality traits (Pank, 2012), while low seed shattering (SHT) and high thousand-grain weight (TGW) are key determinants of seed yield including yield stability (van Roon & Bleijenberg, 1964; von Maydell et al., 2020; von Maydell et al., 2026a). Furthermore, high stalk attachment rates (STA) impair product quality for whole-seed culinary applications.

To overcome this gap, we utilized two extended F_2_ mapping populations (N >280 each) derived from crosses between two biennial accessions (P1: cultivar ‘Rekord’; and P2: wild-derived ‘CARUM 63’) and one annual inbred line (‘L32’), which have been recently established to map flowering time loci (Preprint:von Maydell et al., 2026b). Because a large proportion of F_2_ plants showed annual flowering (P1: 97.9%; P2: 99.7%) and produced viable seeds in the first year of growing, these populations offer high potential for dissecting seed-related traits without strong confounding effects caused by differing flowering types (e.g., in contrast to a GWAS panel comprising both annual and biennial genotypes). These mapping populations are particularly well-suited for QTL detection for three main reasons: First, annual and biennial genotypes belong to the most contrasting gene pools in caraway (von Maydell et al., 2021a). This increases the likelihood that the parents harbor different QTLs for quantitative seed traits, which can be detected in a segregating F_2_ population even if the parents do not display extreme phenotypic contrasts. Second, the maternal biennial parents are genetically distantly related (von Maydell et al., 2021a), increasing the potential to detect diverse QTLs across populations. Third, for seed shattering, which is presumably a monogenically inherited trait (Toxopeus, 1998), the parents of P1 ‘Rekord’ (low/non shattering) and ‘L32’ represent known contrasting phenotypes (von Maydell et al., 2024).

Therefore, by combining novel phenotypic data for seed traits with the SNP-based linkage maps from (<Preprint>) we addressed the following major objectives in this study:

### QTL detection for key agronomic and quality traits

To identify QTLs for (i) major essential oil components, (ii) seed shattering (SHT), (iii) thousand-grain weight (TGW), and (iv) stalk-attachment rate (STA). To efficiently process the large sample size, a multi-year, multi-genotype NIRS-based prediction model was used to estimate essential oil components (von Maydell et al., 2024).

### Analysis of potential pleiotropic effects with following focuses

To investigate whether QTLs associated with essential oils simultaneously affect limonene and carvone contents, and to determine whether previously detected correlations among SHT, TGW, and STA might stem from shared genetic factors. Additionally, potential confounding effects of flowering time were carefully assessed due to its high variability in the F_2_ populations. The goal was not to find evidence for true pleiotropic effects, but rather to screen for potential genetic dependence between traits as crucial information for breeding.

### Design and evaluation of diagnostic markers

To design PACE markers covering peak positions and flanking intervals of key QTLs to facilitate fine-mapping and future marker-assisted selection (MAS). For selected PACE markers, genotype calls were subsequently validated against GBS data.

## 1 Material and methods

### 1.1 Biparental mapping populations, Phenotyping & Data modelling

Crosses and F_2_ population development are described in (von Maydell et al., 2026b). Briefly, two biennial genotypes (P1: cultivar ‘Rekord’; and P2: wild-derived accession ‘CARUM 63’) were crossed with the annual inbred line ‘L32’. The attributes for seed traits of the parents are described in Table 1, predominantly based on the germplasm characterization by von Maydell et al. (2024). Notably, quantitative differences could be confounded by differing flowering types that flowered and ripened in different years. The characterization of the cultivar ‘Rekord’ is in accordance with the official description of the Federal Plant Variety Office (Bundessortenamt, 2002).

**Table 1:** Attributes of the parents of the F_2_ populations. Quantitative data from von Maydell et al. (2024).

| Trait | ‘Rekord’ | ‘CARUM 63’ | ‘L32’ |
| --- | --- | --- | --- |
| **Classification** | Cultivar | Wild-derived | Breeding line (5<sup>th</sup> inbred generation) |
| **Flowering type** | Biennial | Biennial | Annual |
| **Total essential oil content (OIL)** | High<br>(5.9 g/100 g) | Medium<br>(4.9 g/ 100 g) | Very high<br>(8.5 g/ 100 g) |
| **Proportion of carvone (CRVP)** | Medium-high<br>(41.9%) | Medium<br>(38.7%) | Medium-high<br>(42.9%) |
| **Shattering (SHT)** | Non-/low-shattering<br>(29.6%) | Free-/high-shattering<br>(98.5%) | Free-/high-shattering<br>(86%) |
| **TGW** | High<br>(3.1 g) | Low<br>(1.6 g) | Medium<br>(2.1 g) |
| **Stalk attachment rate (STA)** | High<br>(28.8%) | Low<br>(4.1%) | Medium<br>(14.4%) |

The field trial design is described in detail in (von Maydell et al., 2026b). Briefly, 288 plants per F_2_ population were pre-grown in the greenhouse in March (2025) and transplanted to the field in May. Trait measurements largely followed the methods of von Maydell et al. (2024) with some deviations in the number of technical replicates: SHT was measured twice, once before (SHT1) and once after harvest and drying (SHT2) by two different observers to obtain two independent estimates. At maturity, a few seeds were pulled ten times from different umbels. The number of events where seeds were detached from the umbel was counted, resulting in a score ranging from 0 to 10. Deviating from von Maydell et al. (2024), we treated the data as an ordinal score rather than converting it into a continuous rate. This choice accounts for the semi-quantitative nature of the measurement, which resulted from evaluating a single technical replicate due to the limited number of available umbels per single plant. Afterwards, seeds were harvested and dried (Figure 1 A) for approximately two days at 30° C. SHT2 was measured on dry material using the same method as for SHT1. Afterwards seeds were threshed and cleaned. TGW was estimated by weighing 100 randomly selected seeds per sample. Stalk attachment rate (STA) was estimated by counting the number of seeds among those 100 seeds with stalks longer than ∼1 cm (Figure 1 B-C); intact double achenes were regarded as two seeds with one stalk.

**Figure 1:**
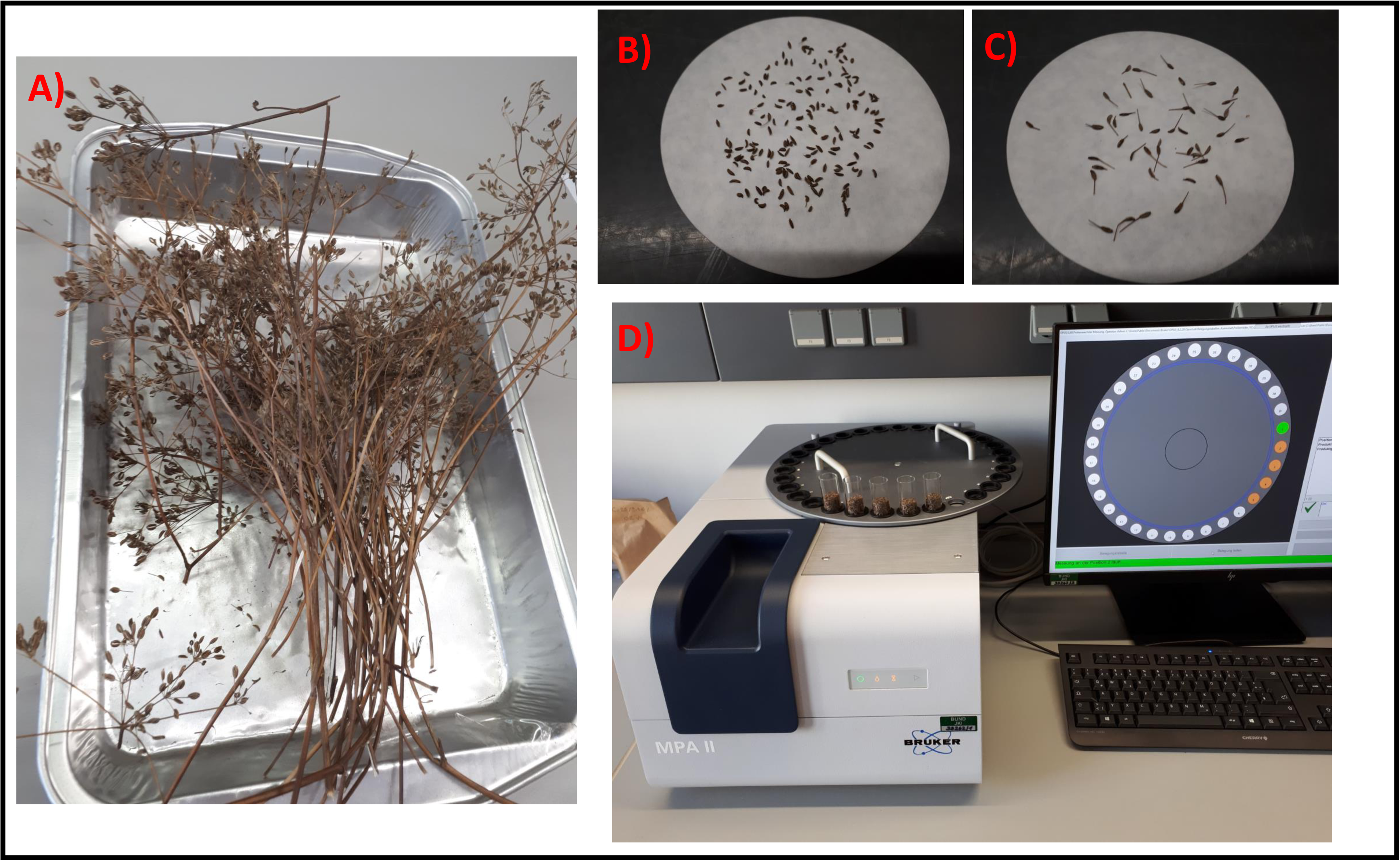
Seed samples. A) Typical single-plant sample before threshing; B) Typical cleaned seed sample with low stalk-attachment rate; C) Sample with high stalk-attachment rate; D) NIRS measurements with MPA2.

Limonene (LIM) and carvone (CRV) contents were estimated using the validated multi-environment, multi-genotype NIRS-based prediction model by von Maydell et al. (2024) with two technical replicates. NIRS measurements were conducted using the FT-NIR Multi Purpose Analyzer II (Bruker, Figure 1D). Technical replication demonstrated high precision and low technical noise for both essential oil components. CRV predictions showed a coefficient of variation for technical replicates (CV_tech) of 5.8% (P1) and 6.8% (P2), and a technical repeatability (R_tech) of 0.82 (P1) and 0.83 (P2). Similarly, LIM predictions showed a CV_tech of 5.6% (P1) and 5.7% (P2), and a slightly higher R_tech of 0.88 (P1 and P2). Based on the mean CRV and LIM estimates, the total essential oil content (OIL) was calculated as the sum of both components, and the proportion of carvone (CRVP) was determined.

Prior to modeling, samples with a seed weight <0.5 g were discarded, resulting in a final sample size of 236 and 266 for P1 and P2, respectively. Seed weight showed very low correlations with all other traits, including beginning of flowering (BOF, P1: r = -0.08), indicating that this filtering step introduced no significant systematic bias. To account for the high variability in flowering time, adjusted trait values were calculated by adjusting CRVP, CRV, LIM, OIL, and TGW for beginning of flowering data from von Maydell et al. (2026b) using simple linear regression (y = trait ∼ BOF) in base R (R Core Team, 2025). The resulting residuals were extracted (Trait_adj). To evaluate the impact of flowering time on genetic architecture, both unadjusted traits and adjusted residuals were subsequently analyzed and compared in genetic mapping. Since SHT1, SHT2, and STA data strongly deviated from normal distribution (including skewness or kurtosis >|1|) we did not perform a parametric adjustment for BOF. Instead, detected QTLs were carefully compared with potentially co-localized QTLs for flowering time.

Figures were created using ggplot2 (Wickham, 2016).

### 1.2 QTL mapping

In this study, we used the linkage maps and the consensus physical map of von Maydell et al. (2026b) built from GBS data. For QTL mapping, the genetic maps thinned to one marker per 1 cM were used (P1: 392 SNPs; P2: 337 SNPs).

QTL mapping was conducted using MapQTL 5 (van Ooijen, 2004). Genome-wide significance thresholds (α = 0.05) were established for each trait through 1,000 permutation tests. We first performed interval mapping (IM) to identify candidate regions, followed by multiple QTL mapping (MQM); the final cofactor selections per trait are detailed in Table S1. Peak positions and phenotypic variance explained (PVE) by individual QTLs were derived from the final MQM models, whereas 1- and 2-LOD support intervals alongside flanking markers were determined via restricted MQM (rMQM). For traits showing pronounced deviations from normality (SHT1, SHT2, and STA), MQM results were cross-validated against non-parametric Kruskal-Wallis test results.

### 1.3 Diagnostic markers for selected flanking and peak markers

PACE markers (3cr Bioscience), each consisting of two allele specific and one common primer, were developed and validated for the peak positions and flanking regions of several key QTLs, as presented in Table 3. To balance both mapping precision and positional reliability, flanking markers were chosen at the boundaries of the 1.5-LOD support interval to balance mapping precision and positional certainty (Dupuis & Siegmund, 1999). Primers were designed by 3cr Bioscience and synthesized by Life Technologies (Thermo Fisher Scientific). Sequence details flanking the target SNP (50 bases upstream and downstream) alongside primer sequences are provided in Table S3.

**Table 2:**
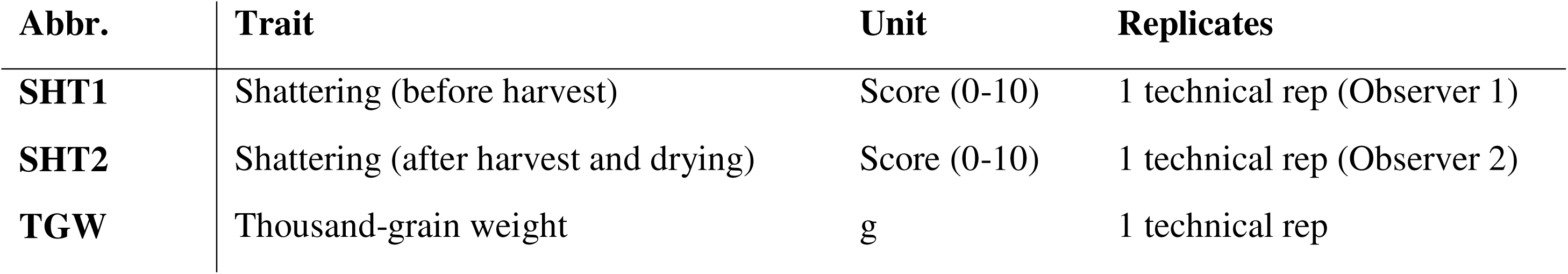
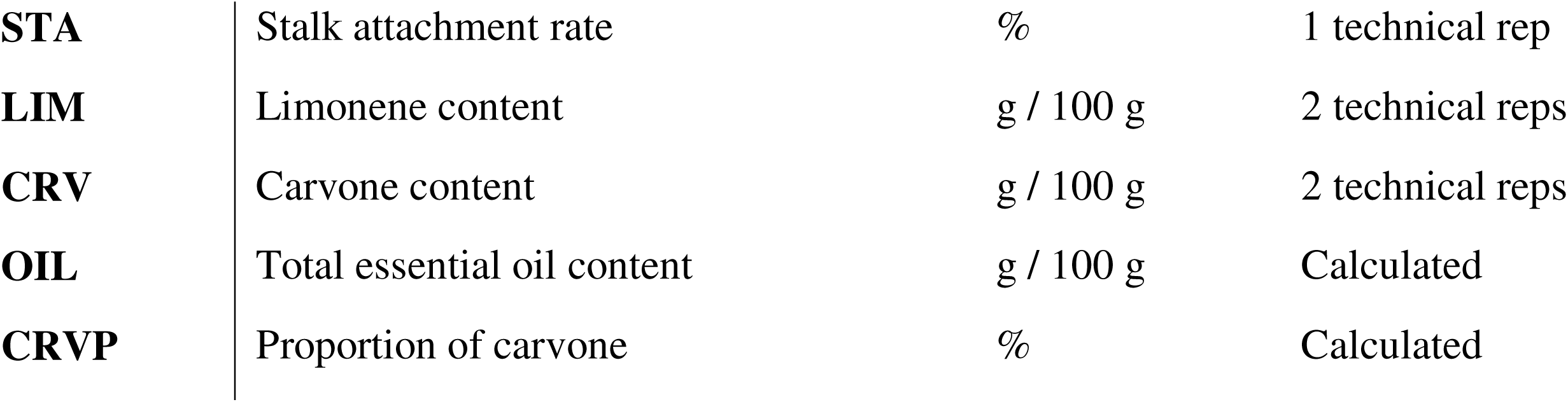
Summary of evaluated traits, units, and number of technical replicates.

**Table 3:** List of 15 representative PACE markers developed in this study and their position to the peak of the QTL. Extended informations including primer sequences and additional markers in Table S3. Peak = in proximity to the peak. Population = F_2_ population, in which QTL was detected.

| **QTL** | **Population** | **Left flank** | **Peak** | **Right flank** |
| --- | --- | --- | --- | --- |
| Cc-SHT03.1 | P1 | M205 | M211 | M215 |
| Cc-OIL08.1 | P1 | M218 | M219 | M221 |
| Cc-OIL05.1 | P2 | M223 | M225 | M227 |
| Cc-OIL10.1 | P2 | M230 | M232 | M233 |
| Cc-TGW02.1 | P2 | M235 | M238 | M240 |

PACE assays were performed using CFX96 real-time system and CFX Maestro software (Bio-Rad), adhering to the protocol described by von Maydell et al. (2020). The standard PCR profile comprised 40 cycles (excluding touchdown) at an annealing temperature of 57°C. To benchmark the assays against GBS results, 88 F_2_ individuals were tested for six selected markers, with parental DNA and extraction buffer serving as controls. To mirror high-throughput screening conditions, the identical DNA samples from the GBS analysis were used without normalizing their concentrations.

## 2 Results

### 2.1 Phenotypic data and trait-correlations

Raw phenotypic data for P1 and P2 are provided in Table S1A and B, respectively. The data distributions and key summary statistics for each trait across both populations are presented in Table 4 and Figure 2. Notably, P2 exhibited a lower mean TGW, which aligns with the lower TGW observed in the biennial wild-derived parent. In contrast, P2 displayed slightly higher average values for OIL, CRV, and LIM, alongside a broader phenotypic range for these traits.

**Figure 2:**
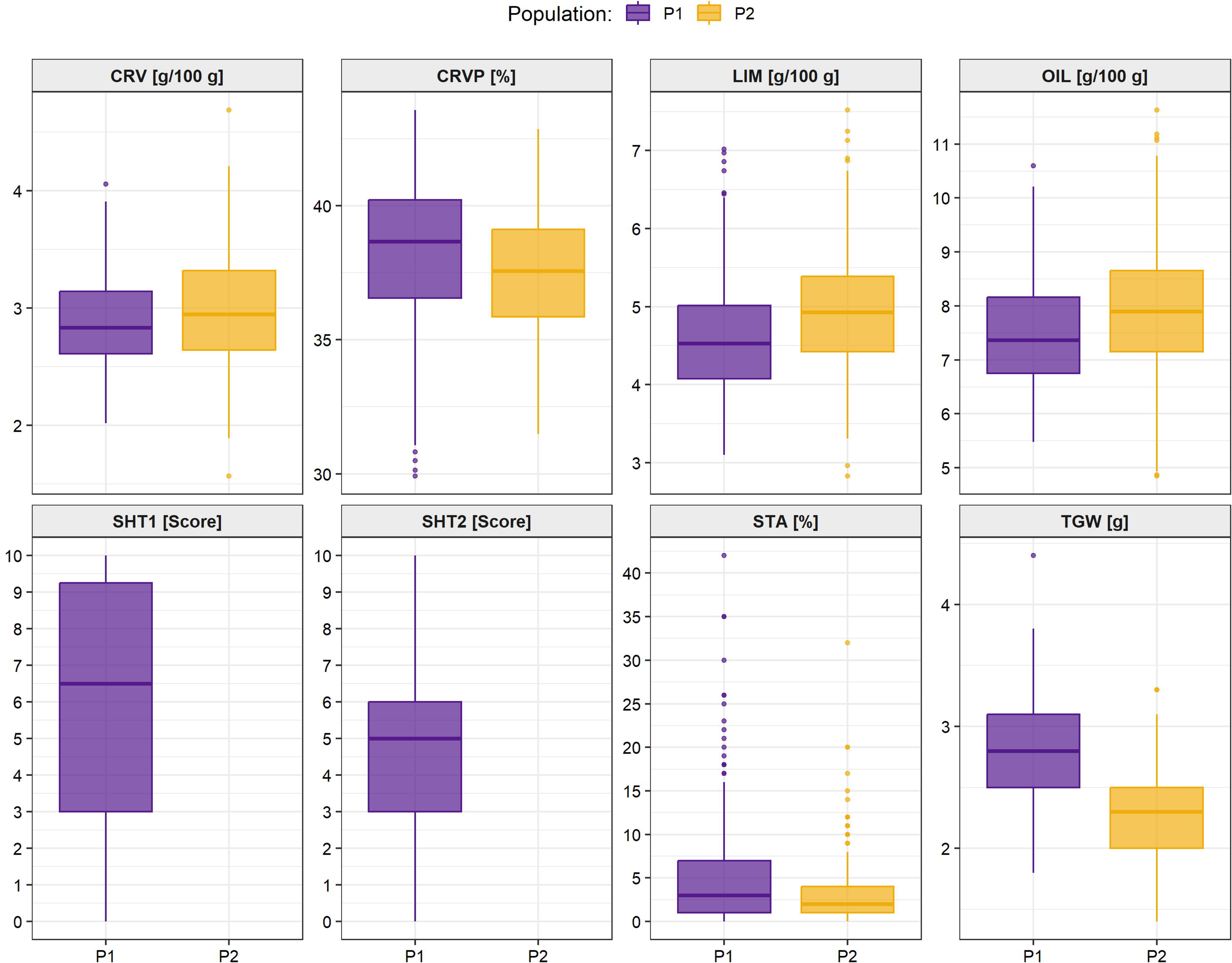
Boxplots illustrate the distribution of phenotypic values for agronomic, chemical, and morphological traits in mapping populations P1 (purple) and P2 (yellow). Horizontal lines represent the median, boxes indicate the interquartile range (IQR), and dots depict statistical outliers. Trait units are specified in brackets.

**Table 4:** Descriptive statistics and phenotypic variation for evaluated traits across F_2_ populations P1 and P2. Trait values are presented as mean, standard deviation (SD), coefficient of variation (CV in %), and range (minimum–maximum).

| Trait | Unit | P1 |  |  |  |  | P2 |  |  |  |
| --- | --- | --- | --- | --- | --- | --- | --- | --- | --- | --- |
|  |  | Mean | SD | CV | Range |  | Mean | SD | CV | Range |
| CRV | g/100 g | 2.88 | 0.37 | 13.0 | 2.02 - 4.06 |  | 2.98 | 0.47 | 15.7 | 1.57 - 4.69 |
| CRVP | % | 38.29 | 2.79 | 7.3 | 29.92 - 43.57 |  | 37.52 | 2.26 | 6.0 | 31.48 - 42.86 |
| LIM | g/100 g | 4.66 | 0.73 | 15.8 | 3.10 - 7.02 |  | 4.97 | 0.79 | 15.8 | 2.83 - 7.52 |
| OIL | g/100 g | 7.53 | 1.02 | 13.5 | 5.48 - 10.60 |  | 7.95 | 1.20 | 15.1 | 4.85 - 11.63 |
| SHT1 | Score | 5.97 | 3.52 | 58.9 | 0.00 - 10.00 |  |  |  |  |  |
| SHT2 | Score | 4.77 | 2.38 | 49.8 | 0.00 - 10.00 |  |  |  |  |  |
| STA | % | 5.11 | 6.60 | 129.0 | 0.00 - 42.00 |  | 3.23 | 3.89 | 120.6 | 0.00 - 32.00 |
| TGW | g | 2.82 | 0.44 | 15.6 | 1.80 - 4.40 |  | 2.25 | 0.35 | 15.5 | 1.40 - 3.30 |

Trait correlations are presented in Figure 3, contrasting Pearson (r) and Spearman (rho) coefficients to accommodate ordinal and non-parametric traits (SHT1, SHT2, STA). Overall, both correlation methods yielded highly consistent results. Essential oil components (OIL, LIM, CRV) displayed strong positive inter-correlations (r = 0.65–0.97), yet remained largely independent of agronomic and morphological seed traits. Deviating from this pattern, CRVP was negatively correlated with flowering time (BOF; r_P1_ = -0.67, r_P2_ = -0.50) in both populations, and showed a moderate positive association with TGW in P2 (r_P2_ = 0.48; vs. r_P1_ = 0.24). Moderate correlations were also observed between BOF and several morphological traits (e.g., r_P2_ = -0.47 for TGW). This suggests potential developmental dependencies, highlighting the importance of evaluating detected QTLs for raw data using adjusted values.

**Figure 3:**
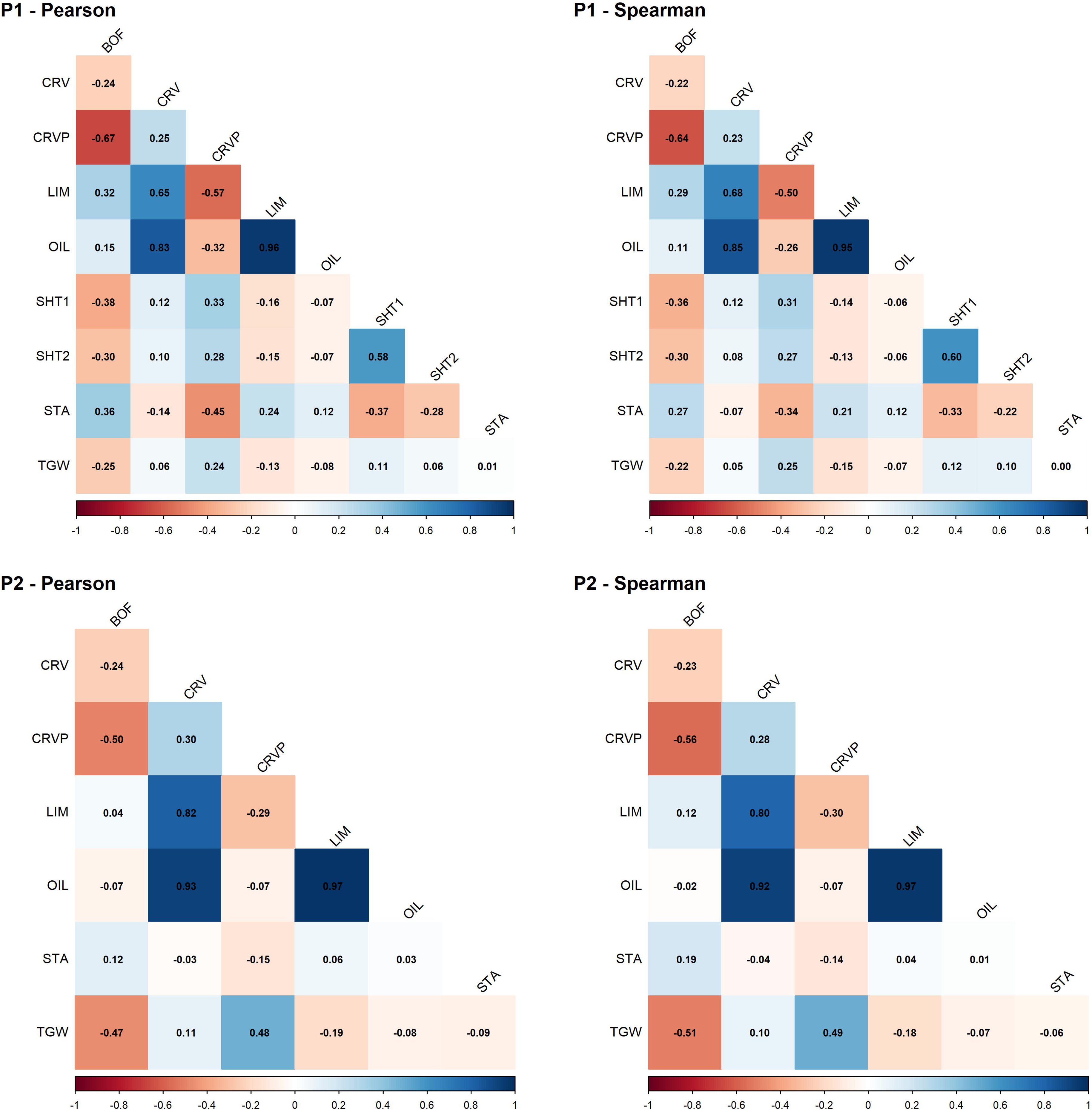
Heatmaps displaying parametric Pearson (r) and non-parametric Spearman (rho) correlation coefficients among in populations P1 (top) and P2 (bottom). Color intensity and numeric values indicate the direction and strength of phenotypic relationships. Trait definitions and units in Table 2.

### 2.2 QTL mapping

#### 2.2.1 Overview

Raw IM, rMQM, MQM, and Kruskal–Wallis results are provided in Table S1C–F. A summary of selected QTLs is provided in Table 5, while an extended version including 2-LOD intervals and flanking markers is provided in Table S1G–H. LOD profiles per trait, including adjusted values, are shown in Figures S1–S2 (IM), Figures 4–5 (rMQM), and Figure S3 (Kruskal–Wallis). In general, IM and Kruskal–Wallis results supported the QTLs detected by MQM. As a minor difference, some QTLs detected by MQM showed sub-threshold maxima in the corresponding IM profiles. Therefore, the following sections focus on the MQM results. In Figure 6, 1-LOD intervals of the detected QTLs are projected onto the linkage maps and the consensus map from von Maydell et al. (2026b). The consensus map allows to gain approximate physical information on QTL confidence intervals and to compare QTLs across studies and populations. Notably, some QTL intervals are affected by the conflicts between P1 and P2 linkage maps, visible by peak markers of P2 residing out of 1-LOD intervals (Figure 6). Figures S4–S5 present effects for allele combinations of all detected QTLs, while Figures 7–9 highlight selected QTLs and QTL combinations.

**Figure 4:**
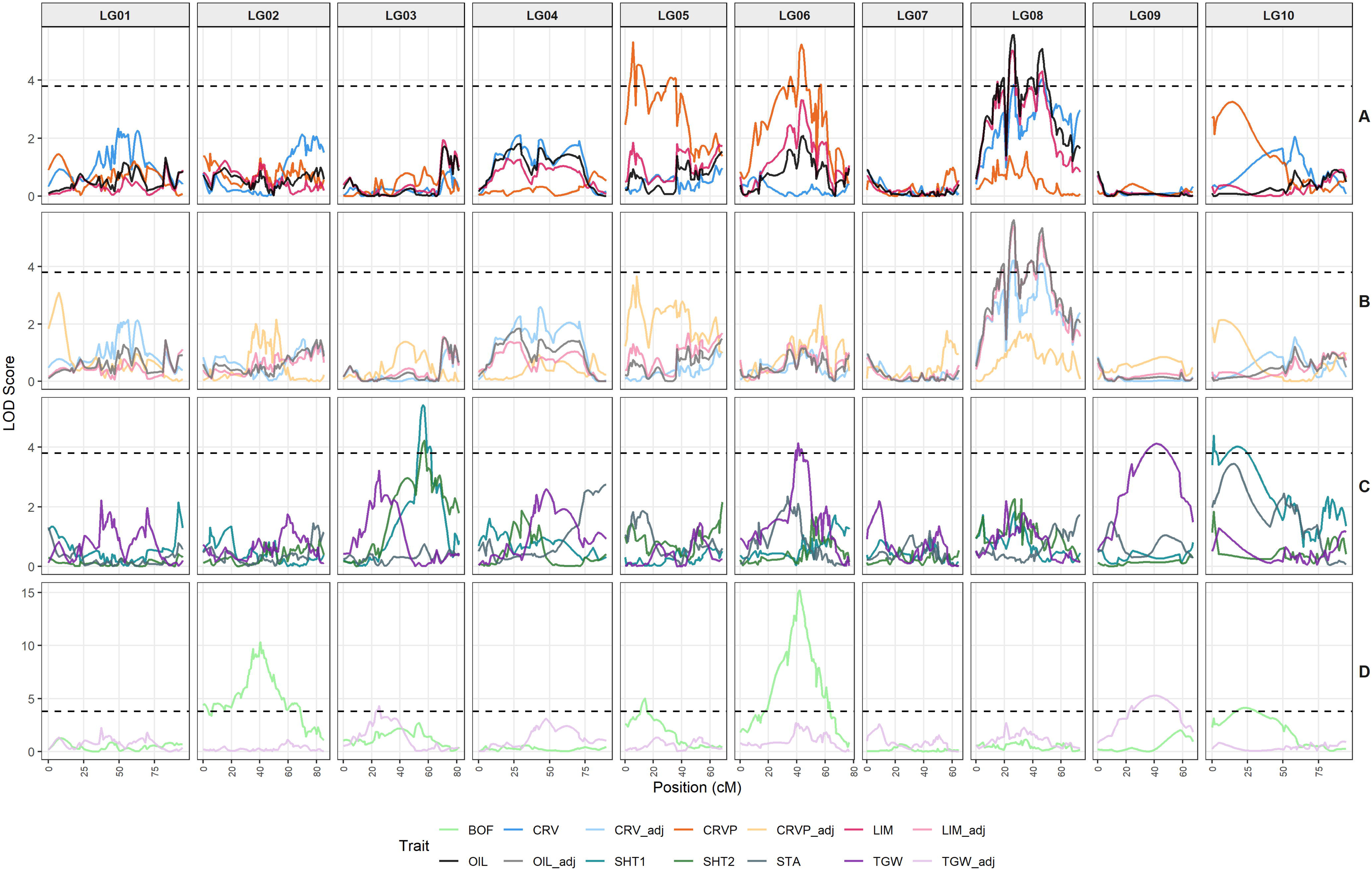
LOD profiles of P1 based on rMQM results. Trait definitions and units in Table 2. Dashed line indicates genome-wide significance-threshold at 3.8, other thresholds (3.6, 3.7 and for STA only 4.0) not shown. Trait definitions and units in Table 2.

**Figure 5:**
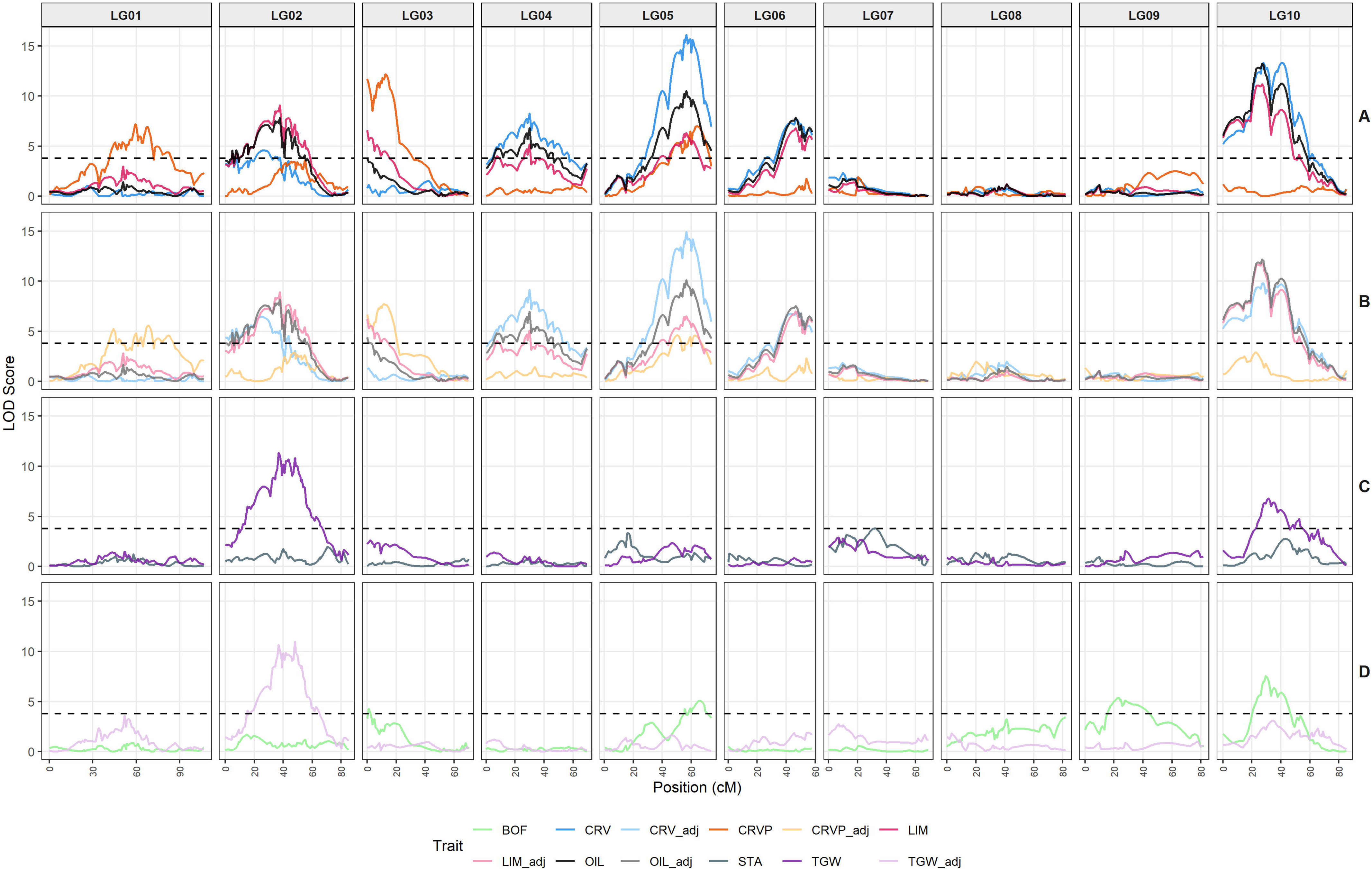
LOD profiles of P2 based on rMQM results. Trait definitions and units in Table 2. Dashed line indicates genome-wide significance-threshold at 3.8, other thresholds (3.5, 3.6, 3.7) not shown. Trait definitions and units in Table 2.

**Figure 6:**
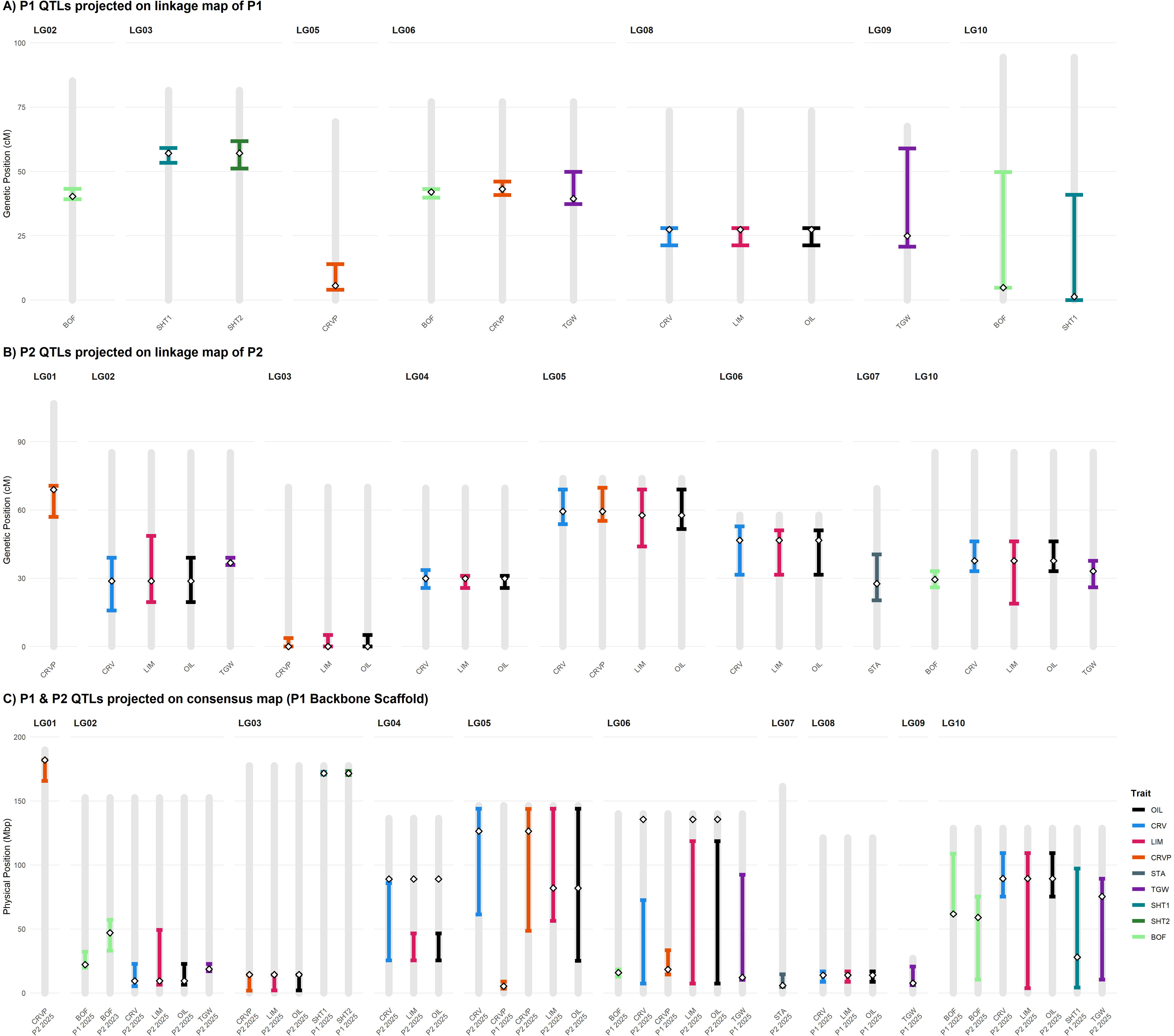
1-LOD intervals of detected QTLs in P1 and P2 (including major flowering time QTLs from von Maydell et al. (2026b)) projected on the respective linkage maps (A and B) and on the consensus map (C) from von Maydell et al. (2026b) using P1 as backbone scaffold. Trait definitions and units in Table 2.

**Figure 7:**
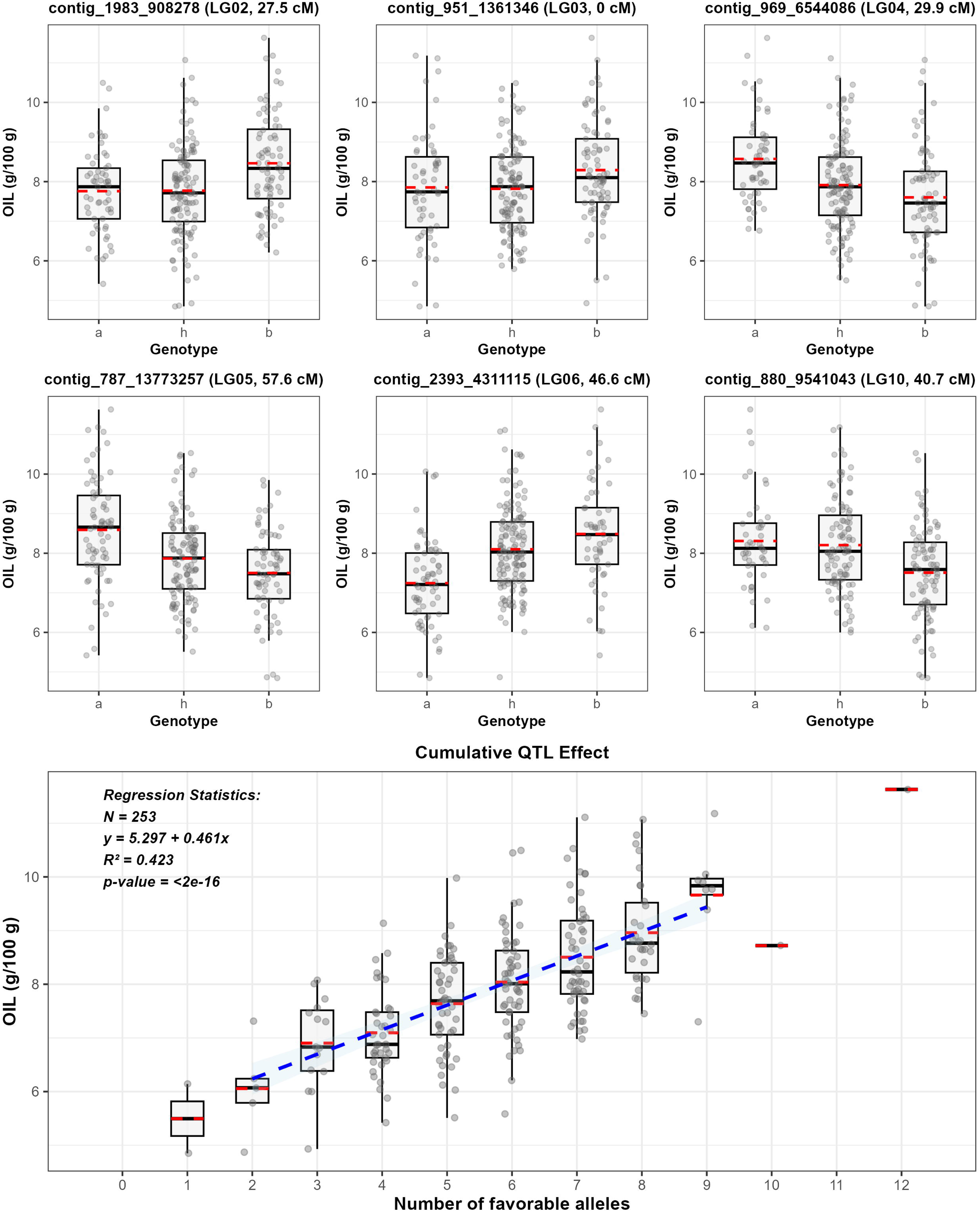
Phenotypic effects of six individual QTLs detected for OIL and their cumulative additive effect. Red dashed lines denote mean values per group, solid lines indicate medians, and gray points depict individual sample observations. Blue dashed line indicates the regression line.

**Figure 8:**
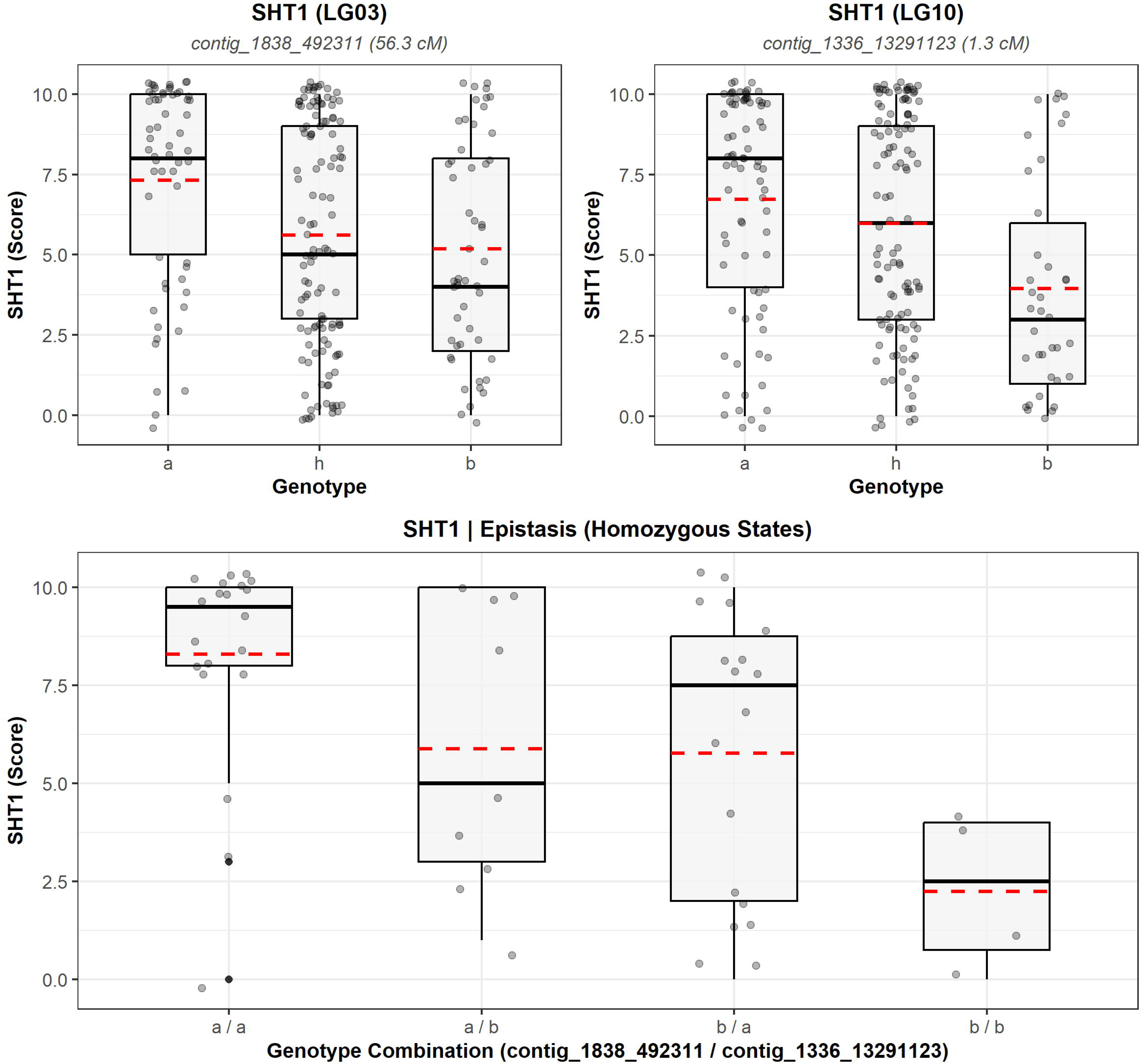
Phenotypic effects of individual QTLs and their epistatic interaction on SHT1. (A, B) Distribution of SHT1 scores across genotypes for loci on LG03 and LG10. (C) Epistatic interaction between both loci considering homozygous genotype combinations. Red dashed lines denote mean values per group, solid lines indicate medians, and gray points depict individual sample observations.

**Figure 9:**
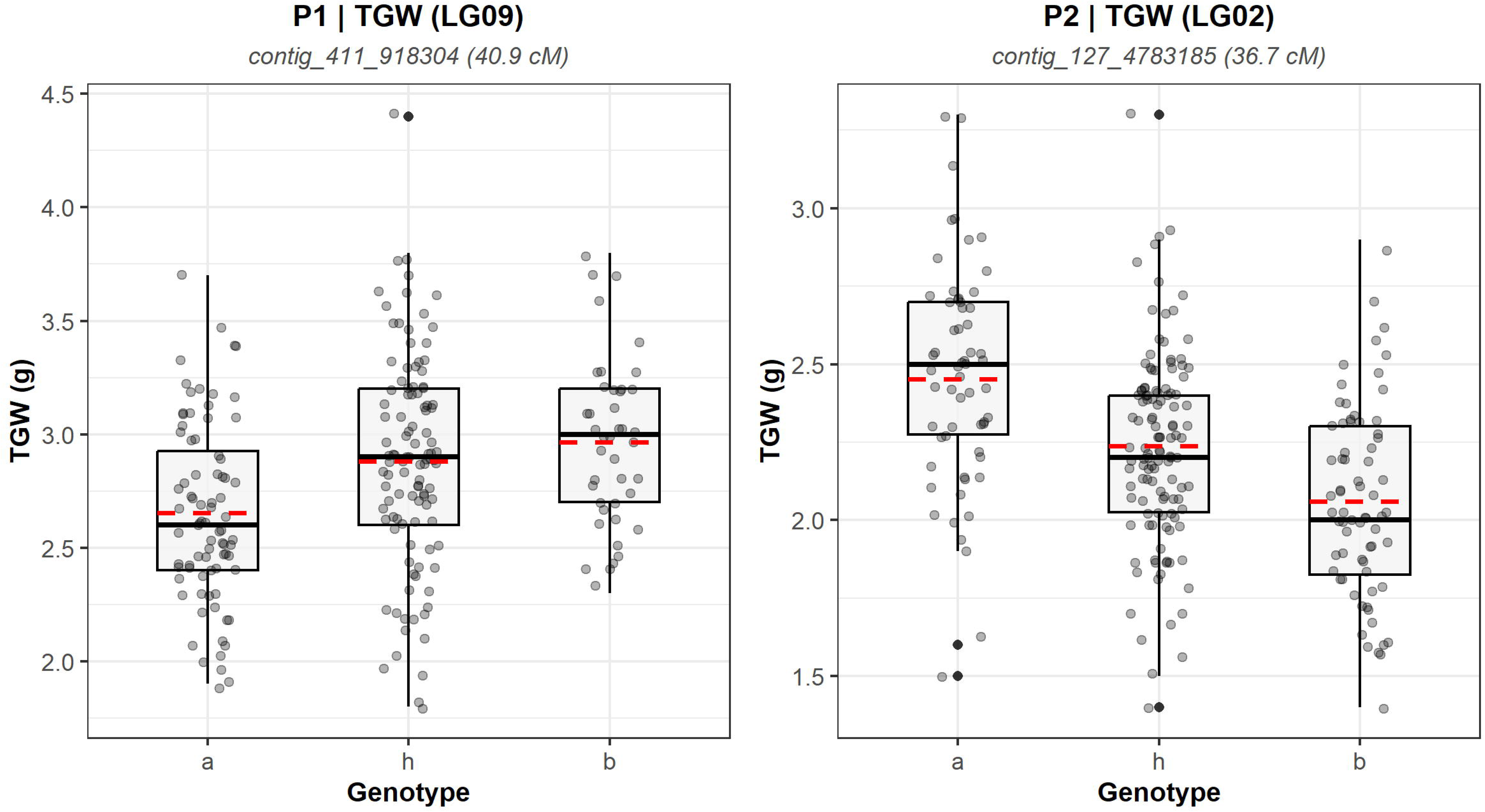
Phenotypic effects of individual major QTLs of P1 and P2 on TGW. Red dashed lines denote mean values per group, solid lines indicate medians, and gray points depict individual sample observations.

**Table 5:** MQM mapping statistics of selected significant QTLs associated with seed traits. The 1-LOD confidence intervals were obtained from the restricted MQM model. Expl. Var. denotes the proportion of phenotypic variance explained (%), while Add. and Dom. represent the additive and dominance effects of the annual allele, respectively. Trait definitions and units in Table 2.

| QTL_ID | Population | Trait | LG | Peak cM | 1-LOD<br>cM | LOD | PVE<br>% | Add | Dom | Peak Locus |
| --- | --- | --- | --- | --- | --- | --- | --- | --- | --- | --- |
| Cc-OIL08.1 | P1 | OIL | LG08 | 26.56 | 23.23 - 27.95 | 5.56 | 10.5 | 0.21 | 0.59 | contig_2320_2322072 |
| Cc-SHT03.1 | P1 | SHT1 | LG03 | 56.32 | 53.32 - 59.06 | 5.41 | 9.9 | 1.13 | -1.43 | contig_1838_492311 |
| Cc-SHT10.1 | P1 | SHT1 | LG10 | 1.28 | 0.00 - 30.79 | 4.39 | 7.6 | 1.46 | 0.49 | contig_1336_13291123 |
| Cc-TGW09.1 | P1 | TGW | LG09 | 40.93 | 21.76 - 58.66 | 4.11 | 11.1 | -0.13 | 0.20 | contig_411_918304 |
| Cc-OIL01.1 | P2 | CRVP | LG01 | 68.03 | 56.97 - 70.63 | 6.92 | 8.9 | 0.93 | 0.17 | contig_5215_20225 |
| Cc-OIL02.1 | P2 | OIL | LG02 | 27.47 | 21.47 - 38.70 | 7.09 | 7.2 | -0.39 | -0.31 | contig_1983_908278 |
| Cc-OIL03.1 | P2 | OIL | LG03 | 0 | 0.00 - 4.68 | 3.88 | 3.8 | -0.32 | -0.12 | contig_951_1361346 |
| Cc-OIL04.1 | P2 | OIL | LG04 | 29.87 | 25.77 - 30.87 | 6.8 | 6.8 | 0.45 | -0.18 | contig_969_6544086 |
| Cc-OIL05.1 | P2 | OIL | LG05 | 57.64 | 51.65 - 63.08 | 9.94 | 10.2 | 0.54 | -0.04 | contig_787_13773257 |
| Cc-OIL06.1 | P2 | OIL | LG06 | 46.65 | 40.53 - 51.08 | 7.88 | 7.9 | -0.50 | 0.15 | contig_2393_4311115 |
| Cc-OIL10.1 | P2 | OIL | LG10 | 40.65 | 35.09 - 44.65 | 11.26 | 13.3 | 0.48 | 0.45 | contig_880_9541043 |
| Cc-STA07.1 | P2 | STA | LG07 | 31.60 | 24.32 - 37.60 | 3.81 | 8.5 | 1.24 | -1.73 | contig_91_5831757 |
| Cc-TGW02.1 | P2 | TGW | LG02 | 36.74 | 35.81 - 38.70 | 11.34 | 15.6 | 0.19 | -0.01 | contig_127_4783185 |

For future reference, QTL IDs (Table 5 and Table S1) were defined according to the following criteria: All QTLs related to essential oil traits were denoted with ‘OIL’, and QTLs for CRV, CRVP, LIM, and OIL showing strong co-localization of their peaks were assigned identical IDs. For QTLs that lost significance after adjustment for flowering time and co-localized with previously identified flowering-time QTLs, no separate trait-specific QTL ID was assigned, as their effects could not be distinguished from the effects associated with flowering time. For example, no separate TGW ID was assigned to the signal co-localizing with Cc-FT06.1 in P1. However, as it cannot be excluded that such co-localized signals represent distinct but closely linked genetic loci, alternative unambiguous IDs are provided in Table S1.

#### 2.2.2 Essential oil content (OIL, LIM, CRV, CRVP)

For total essential oil content (OIL), a pronounced difference was observed between the two populations. In P1, only one QTL, Cc-OIL08.1 on LG08, was detected (LOD = 5.56; PVE = 10.5%, Table 5, Figure 4A). The annual-parent allele showed a positive additive effect (Add = 0.21), while a comparatively strong dominance effect was observed (Dom = 0.59). Potentially, there could be a second QTL at about 46 cM because in MQM the variance is not completely absorbed by the co-factor at 27 cM and in rMQM there is a strong decay of the LOD value in between of both loci (Figure 4A).

In contrast, six QTLs for OIL were identified in P2 on LG02, LG03, LG04, LG05, LG06, and LG10. Their LOD scores ranged from 3.88 to 11.26 and explained 3.8–13.3% of the phenotypic variance individually (in total 49.2%, Table 5, Figure 5A). The annual-parent allele showed positive additive effects at Cc-OIL04.1, Cc-OIL05.1, and Cc-OIL10.1 (Add = 0.45, 0.54, and 0.48, respectively), whereas negative effects were observed at Cc-OIL02.1, Cc-OIL03.1, and Cc-OIL06.1 (Add = -0.39, -0.32, and -0.50, respectively). These QTLs showed a substantial cumulative effect on OIL in a simplified regression analysis ignoring effect size and dominance effects (Figure 7). Each additional favorable allele was associated with an average increase of approximately 0.46 g/100 g. The regression model explained 42.3% of the phenotypic variance (R² = 0.423).

QTL effects on essential oil composition were partly distinct from those affecting total oil content. Notably, Cc-OIL03.1 on LG03 affected LIM but not CRV (Figure 5A, Table S1H). As no corresponding effect on CRV was detected, this was accompanied by an increase in the relative proportion of carvone in relation to the annual-parent allele (CRVP; Add = 1.23). Cc-OIL05.1 on LG05 also affected CRVP, with a positive additive effect of the annual-parent allele (Add = 0.87), while no corresponding CRVP effect was detected for the other OIL QTLs (Table 5, Figure 5A). Cc-OIL01.1, detected in P2 on LG01 (LOD = 6.92; PVE = 8.9%), was particularly distinct, as it affected CRVP but not CRV, LIM, or OIL (Table 5, Figure 5A). The annual-parent allele showed positive effect on CRVP (Add = 0.93). Thus, Cc-OIL01.1 represents a specifically detected locus affecting oil composition rather than total essential oil content.

In P1, two additional significant CRVP QTLs were detected before adjustment for flowering time but disappeared or fell below the significance threshold after correction (Figure 4A-B), indicating that their effects on CRVP were not independent of flowering time.

#### 2.2.3 Shattering (SHT1, SHT2) & Stalk attachment rate (STA)

For shattering, two QTLs were identified in P1 for SHT1. The main locus, Cc-SHT03.1, was mapped to LG03 (LOD = 5.41; PVE = 9.9%, Table 5, Figure 4C), where the annual-parent allele increased shattering (Add = 1.13) alongside a substantial negative dominance effect (Dom = −1.43). Notably, this locus was also detected for SHT2 (LOD = 3.98; PVE = 7.8%; Add = 0.84; Dom = −0.58, Table S1G, Figure 4C), consistent with the shattering-promoting effect of the annual allele. The second SHT1 locus, Cc-SHT10.1, was located on LG10 (LOD = 4.39; PVE = 7.6%, Table 5) and similarly showed a positive additive effect (Add = 1.46), with the annual-parent allele enhancing shattering. We observed a considerable difference between the homozygous a/a and b/b combinations of both SHT1 loci, but the b/b combination had a very low sample size (Figure 8). Interestingly, this QTL was not detected for SHT2, measuring SHT after harvest and drying.

For STA, only a single minor QTL was detected in P2 on LG07 (Cc-STA07.1, LOD = 3.81; PVE = 8.5%), showing a positive additive effect (Add = 1.24), indicating an increasing effect of the annual-parent allele (Figure 5C, Table 5). Interestingly, a subthreshold maximum was observed on LG10 co-localizing with QTLs for SHT1 and BOF and (Figure 5C).

#### 2.2.4 Thousand-grain weight (TGW)

For TGW, two QTLs were identified. Cc-TGW09.1 was detected in P1 on LG09 (LOD = 4.11; PVE = 11.1%) and showed a negative additive effect (Add = −0.13), indicating a decreasing effect of the allele from the annual parent (Figure 4C, Table 5, Figure 9). In P2, Cc-TGW02.1 was detected on LG02 with a substantially higher LOD score (11.34) and explained 15.6% of the phenotypic variance. Its additive effect was positive (Add = 0.19), indicating an increasing effect of the annual-parent allele. Interestingly, in both populations, an additional QTL co-localized with the major flowering-time locus (P1: LG06; P2: LG10, Figures 4-6). In both cases, the annual-parent allele was associated with an increasing effect on TGW (Table S1G–H, Figure S4–5).

### 2.3 Diagnostic markers for selected flanking and peak markers

In total, we designed 26 PACE markers representing the flanks and peaks (1–2 markers per position) of five QTLs for OIL, SHT, and TGW, which clearly discriminated the parental genotypes. For six markers, we additionally performed PACE assays on 88 F plants and compared the PACE calls with the GBS calls. The conversion rate ranged from 92.41% to 100.00% (Table 6). In most cases (72.22%), mismatches resulted from a homozygous call in the PACE assay compared to a heterozygous call in the GBS SNP data.

**Table 6:** Comparison of GBS and PACE results for selected flanking and peak markers.

| Marker | Sample Size | PACE Failed | GBS Failed | Valid Comparisons | Matches | Conversion Rate |
| --- | --- | --- | --- | --- | --- | --- |
| M205 | 88 | 5 | 5 | 80 | 79 | 98.75 |
| M211 | 88 | 0 | 5 | 83 | 77 | 92.77 |
| M215 | 88 | 3 | 5 | 80 | 77 | 96.25 |
| M218 | 88 | 3 | 6 | 79 | 73 | 92.41 |
| M219 | 88 | 2 | 5 | 82 | 82 | 100.00 |
| M221 | 88 | 2 | 5 | 81 | 79 | 97.53 |

## 3 Discussion

### 3.1 General remarks on population design

Annual and biennial genotypes represent the most distinct gene pools in caraway (von Maydell et al., 2021a). Therefore, biparental mapping populations from crosses between these gene pools may be the most promising resources for detecting a high number of QTLs for various traits. This assumption is supported by the QTL detection in this study showing a broad spectrum of detected QTLs. We consider that using an F_2_ population directly is the most effective way for QTL detection, considering the short time required for its development and the relatively low bias. For example, a GWAS panel of annuals and biennials would have a much stronger bias due to population structure and different harvesting years. The limitations to disclose potential QTLs have been shown by von Maydell et al. (2022). Due to the high proportion of annual-flowering plants in our F_2_ populations under early sowing conditions (von Maydell et al., 2026b), only a small number of biennial F_2_ plants (P1: N = 7; P2: N = 1) were excluded from the analysis. As in our study, linear adjustments or comparison of co-localized QTLs can be used to account for effects of flowering time on a quantitative scale.

By choosing two extremely distantly related biennial maternal parents, we focused on and succeeded in revealing diverse QTLs for seed traits in caraway. As a disadvantage, both populations provide little cross-validation of the detected QTLs, which was already observed for flowering time in (von Maydell et al., 2026b). Lacking comparative studies in caraway, we also cannot compare our QTLs with previous findings. Hence, each QTL requires further validation, e.g., in F_3_ generations or in independent populations. In the following sections, however, we will discuss the detected QTLs under the assumption that they represent reproducible genetic effects.

### 3.2 Essential oil content (OIL, LIM, CRV, CRVP)

It is an interesting question why our dataset shows such a strong difference in QTL detection for essential oil traits between P1 (one QTL) and P2 (six QTLs, all distinct from the P1 QTL). First, P1 plants were much more severely affected by diseases, including powdery mildew, which was likely a major reason for lower seed setting resulting in the lower sample size (236) compared to P2 (266). (Besides, the visually healthier status of P2 plants, also in comparison to the annual parent, may be of interest regarding the wild-derived ‘Carum 63’ accession as a potential source of resistance.) Second, in the 1990s, high OIL biennial cultivars were introgressed into annual breeding material to increase its OIL. Afterwards, this material was intensely selected for high OIL and an annual flowering habit. Therefore, due to historical intercrossing and selection pressure, the cultivated biennial parent ‘Rekord’ and the annual breeding line ‘L32’ might share several identical QTL alleles. This is consistent with the lower difference in OIL between these parents measured in earlier field trials (von Maydell et al., 2024) and the lower range of OIL in F_2_ in this study.

Nevertheless, the single QTL detected in P1, Cc-OIL08.1, represents an interesting target for breeding. The favorable allele originates from the high-OIL annual parent. Since inbred lines per se show inbreeding depression in caraway, current breeding efforts focus on synthetic outcrossed populations to achieve sufficient yield (von Maydell et al., 2021b). In this regard, the strong dominance effect of this QTL could be highly advantageous. For establishing MAS for this QTL, it will be crucial to verify whether a second putative QTL is located approximately 20 cM away from Cc-OIL08.1.

Naturally, the six QTLs detected in P2 warrant particular attention, especially considering that in three cases, the wild-derived parent ‘Carum 63’ carries the favorable alleles, which might not yet be present in current breeding material. The linear regression of the cumulative number of favorable alleles against OIL represents a simplified model that omits effect sizes and dominance effects.

However, the high R^2^ value of 42% (comparable to the 49% PVE in MQM, which accounts for these effects) highlights the strong potential for pyramiding these alleles to increase OIL through breeding using their additive effects. Nevertheless, the QTL on LG03 should be utilized with caution, as it increases OIL predominantly by increasing limonene levels. Because the European Pharmacopoeia requires a minimum CRVP of >50% (Ph. Eur., 2020), this effect could be detrimental for commercialization. Notably, the low CRVP values observed in this study (all <50%), but high OIL values, are likely attributable to the analytical method used, which yields lower CRVP values, but higher OIL values, than steam distillation (the reference method of the Ph. Eur.).

Regarding pleiotropic effects, we observed that QTLs usually affected both LIM and CRV, indicating that the underlying causative genes act prior to LIM or CRV synthesis. Major exceptions are QTL Cc-OIL03.1, which exclusively affects LIM, and QTL Cc-OIL01.1, which affects CRVP without altering total terpenoid content. We found low phenotypic correlations of LIM, CRV, and OIL with other traits, which is consistent with observations from previous field trials (von Maydell et al., 2021b; von Maydell et al., 2024), that also demonstrated environment-dependent variations in the direction of correlation. This lack of strong phenotypic correlation aligns with the genetic data, as no OIL QTLs clearly co-localize with QTLs for other traits. Overall, these weak phenotypic and genetic correlations are advantageous for breeding, as desirable essential oil attributes can be selected largely independently of other agronomic traits. CRVP partially deviates from this pattern in P1, where a moderate correlation between CRVP and BOF as well as co-localization of corresponding loci on LG06 (a major flowering time locus) were observed. However, in two-year field data from a diversity set, only low correlations between BOF and CRVP were detected for biennials (r = -0.24), and no correlation was found for annuals (von Maydell et al., 2024). Thus, the association between CRVP and flowering time observed in P1 may represent a population-specific or single-year effect.

### 3.3 Shattering (SHT1, SHT2) & Stalk attachment rate (STA)

Toxopeus (1998) reported monogenic control of SHT, with non/low-shattering being dominant; however, it should be noted that we lack access to the methodology, raw data, or statistical analyses that led to this assumption. The QTL Cc-SHT03.1 on LG03, which was detected for both SHT1 and SHT2, aligns well with this hypothetical monogenic locus, as it also exhibits a substantial dominance effect originating from the non/low-shattering parent (‘Rekord’). Admittedly, the PVE and LOD scores for this locus appear relatively low for a trait under monogenic inheritance. However, the MQM results must be regarded as an approximation due to the non-parametric nature of this trait. Additionally, the difficult and partially subjective evaluation of SHT likely introduces a considerable degree of random noise. As discussed by von Maydell et al. (2026a), non/low-shattering reduces the likelihood of seed loss caused by heavy rain or strong winds, though some seed dislodgement inevitably occurs. This also affected our scoring, making a ‘zero SHT’ rating extremely rare in our dataset. Specific observational challenges included variations in ripening behavior, such as obvious variable water retention in ripening shoots towards maturity, and asynchronous ripening across different shoots. For a more precise evaluation of SHT, future studies might consider macro- to microscopic analyses of the abscission layer between seed and stalk similar to Toxopeus (1998), however, such a method were not feasible for the high-throughput phenotyping required in this study. Notably, the QTL Cc-SHT03.1 does not co-localize with any STA QTL. Therefore, although it seems reasonable that a missing abscission layer between seed and stalk inadvertently increases STA (Toxopeus, 1998), we find no evidence to support this assumption.

Interestingly, we detected a second QTL for SHT1 on LG10, which, however, was no longer detected when estimating SHT after drying (SHT2). Co-localizing with this locus, we observed a significant QTL for BOF. This suggests a potential association between plant development and SHT. As one possible explanation, the biennial parent allele at the LG10 locus might be associated with an unusual ripening phenotype. The resulting phenotyping artifact may have been mitigated by the subsequent drying process so that the QTL vanished in SHT2. Consequently, the locus on LG10 warrants further investigation before confirming it as a genuine second QTL for SHT. Interestingly also a sub-threshold maximum for STA co-localized with the QTL for SHT1 on LG10. This locus might account for the low to medium correlation observed between STA and SHT, an effect that could potentially be eliminated by selecting the allele originating from the annual parent.

### 3.4 Thousand-grain weight (TGW)

Notably, in both populations, we observed co-localization between major flowering time QTLs and TGW loci, which aligns with the moderate to strong correlations detected between these traits both in this study and in the literature (von Maydell et al., 2021b; von Maydell et al., 2024). Given the remarkably similar LOD profiles, these findings may point to pleiotropic effects controlled by the same causative genes. Although we cannot conclusively prove pleiotropy here, such genetic associations are well-documented across various plant species (Zanke et al., 2015; Li et al., 2026). Because the favorable (increasing) TGW allele is coupled with the early-flowering allele from the annual parent in both cases, this association is highly advantageous for breeding annual cultivars. Of particular interest is the major QTL Cc-TGW02.1 detected in P2, which acts independently of flowering time. While the annual parent allele again represents the favorable, TGW-increasing allele, the annual allele of Cc-OIL02.1, located just 10 cM away from Cc-TGW02.1, exhibits a decreasing effect on OIL. Consequently, marker-assisted selection (MAS) at both loci could be exceptionally valuable for differential selection in breeding programs. Overall, this genomic region on LG02 (contigs 127 and 1125) represents a highly promising target for breeding, as it harbors QTLs for BOF, CRV, LIM, OIL, and TGW across populations.

### 3.5 Candidate genes

Table S2 lists transcripts annotated within the 1-LOD intervals, extended to the complete contig at the interval boundaries to account for uncertainties in contig orientation and discrepancies in marker order between the linkage maps and draft assembly. Furthermore, it is possible that the causal gene resides outside of the 1-LOD interval. This pragmatic approach may overestimate transcript numbers and limit their comparability between QTLs. Nevertheless, all QTL intervals contain several hundred genes, precluding a meaningful candidate gene assignment without further fine mapping. We therefore highlight only selected plausible candidates for a few QTLs in proximity to the peak markers. Notably, terpene synthase encoding genes are rare within the OIL QTL intervals and do not co-localize with the peak markers. Instead, several genes encoding transcription factors or transporters are located in close proximity to these peak markers, representing plausible candidate genes for OIL as well (Gani et al., 2021; Huang et al., 2023). In close proximity (<0.02 Mb) to the peak marker of Cc-SHT-03.1, a gene encoding a MYB transcription factor is located. While transcription factors in general have been found to regulate seed shattering across plants (Dong & Wang, 2015), MYB transcription factors specifically have so far been identified in rice (Ning et al., 2023). For the QTL Cc-TGW-02.1, a gene encoding a glucose-6-phosphate dehydrogenase co-localizes with the peak marker, which could be linked to seed development (Ruan et al., 2022).

### 3.6 Diagnostic markers for fine mapping

We successfully validated 26 PCR-based PACE markers covering five major QTLs. The conversion rate based on GBS data was generally high, and the observed mismatches may rather be attributable to incorrect heterozygous SNP calls in the GBS data, although this cannot be conclusively determined. Notably, we do not claim that these markers can be directly used as diagnostic markers for a trait or across different genetic backgrounds. They may serve for marker-assisted selection (MAS) of a QTL when the selection of large genomic segments from one parent is acceptable, but primarily the PACE markers may be used for recombinant selection as a first step towards future fine mapping. Subsequently, additional PACE markers could be developed based on the dense genetic maps to saturate the genomic region of interest.

### 3.7 Limitations & future prospects of genetic analyses in caraway

#### 3.7.1 Genetic frameworks

The recent establishment of genetic maps, an annotated draft assembly, and the detection of QTLs for flowering time (von Maydell et al., 2026c; von Maydell et al., 2026b), represent important milestones in genetic and breeding research of caraway, which are complemented by the QTL detection for seed traits in this study. However, these current genetic frameworks may still harbor limitations that could affect their successful application in MAS. For example, discrepancies between the linkage maps of P1 and P2, resulting in peak markers of QTLs detected in P2 residing outside their confidence intervals (Figure 6C), may indicate mapping errors or structural chromosomal variation between parental genotypes, which may not be adequately represented by a single reference genome (Yuan et al., 2021). A chromosome-level assembly and in-depth sequencing of parental genotypes could help to resolve such discrepancies.

#### 3.7.2 QTL diversity

While the number of QTLs detected in crosses between biennial and annual caraway is already high, a sole focus on these QTLs, which currently need to be considered as population-specific, might be misleading for breeding programs. GWAS panels or additional biparental to multiparental mapping populations might be powerful tools to detect novel QTLs or to re-detect known QTLs in different genetic backgrounds (Würschum, 2012). In general, it would be valuable for breeding decisions to determine which alleles of major QTLs are present in which breeding lines or cultivars. In particular, it will be interesting to assess whether favorable alleles of the wild-derived parent detected in P2 are also present in more applied material, because introgression of wild material may require selection over several generations to reduce linkage drag (Kashyap et al., 2022).

#### 3.7.3 QTL validation & fine mapping

Beyond improvement of genetic frameworks and further QTL detection, validation of individual QTLs and fine mapping remain laborious challenges for future research. Currently, confidence intervals are likely too large for efficient MAS, unless selection of large chromosomal regions is acceptable. While a single F_1_ individual can produce a sufficiently large F_2_ population (>2,000 plants) for screening recombinants, genotyping and phenotyping of segregating F_3_ families require extensive resources and can therefore realistically only be applied to selected QTLs (Collard & Mackill, 2008). To pave the path towards fine mapping, this study provides starting PACE markers for five key QTLs. Validation of QTL effects can be performed in parallel with fine mapping using F_3_ families with defined allelic status. In particular, QTL effects should be assessed under targeted environmental conditions (El-Soda et al., 2014), because the effects measured in this study are based on somewhat artificial growing conditions. While functional validation of candidate genes is often the final goal in major crops, we consider that sufficiently reducing QTL confidence intervals to enable applied MAS and validating QTL effects may, in most cases, be more realistic objectives in an orphan crop like caraway.

#### 3.7.4 Final remarks

Recent breeding research in seed crops of the Apiaceae family, such as coriander and fennel, has provided valuable insights into genetic regulation through genome assemblies, transcriptomics, and in silico analyses of relevant gene families (Palumbo et al., 2018a; Palumbo et al., 2018b; Song et al., 2019; Song et al., 2020). However, population-based approaches such as GWAS and QTL mapping remain scarce in seed Apiaceae crops, further including anise, dill and cumin (Archangi et al., 2019; Palumbo et al., 2021; Magon et al., 2025), whereas they are frequently conducted in carrot as a root Apiaceae crop for various agronomic and quality traits (Cavagnaro et al., 2014; Ellison et al., 2018; Brainard et al., 2022; Shibaya et al., 2022; Dunemann et al., 2022; Rolling et al., 2025). As demonstrated in carrot, forward-genetic approaches are essential for linking genetic variation to breeding-relevant traits. Therefore, recent QTL mapping efforts in caraway, complemented by the findings on seed-related traits presented in this study, provide crucial insights to accelerate applied breeding.

## Glossary

CRV: Carvone content
OIL: Essential oil content
LIM: Limonene content
P1: F_2_ Population P1 originating from the cross of the biennial cultivar ‘Rekord’ x annual inbred line ‘L32’
P2: F_2_ Population P2 originating from the cross of the biennial wild-derived accession ‘CARUM 63’ x annual inbred line ‘L32’
SHT: Shattering
STA: Stalk attachment rate
TGW: Thousand-grain weight

## Declaration of generative AI and AI-assisted technologies in the manuscript preparation process

During the preparation of this work, the authors used Gemini to improve structure and general readability of the manuscript. The authors reviewed and edited the output as needed and take full responsibility for the content of the published article

## Acknowledgments

We thank the entire team at JKI, which participated in the field works. Particular thanks to Beate Kamm, Thomas Heck, Saba Gharaati, Kathrin Wolf, Bettina Albrecht, Kay Fischbach and Johanna Sophia Schmidt.

## Supplementary

**Table S1:** Raw data and summaries from biparental mapping. Detailed explanations/definition within tables. A/B) Phenotypic data; C/D) Mapping Results; E/F) Kruskal-Wallis, G/H) Full QTL Tables, I) List of genes in 1-LOD intervals of selected QTLs.

**Table S2:** Annotated transcripts in selected QTL intervals and summary.

**Table S3:** Detailed information of PACE markers including primer sequences.

**Figure S1:** LOD score profiles of P1 based on IM results. Trait definitions and units in Table 2. Dashed line indicates genome-wide significance-threshold at 3.8, other thresholds (3.6, 3.7 and for STA only 4.0) not shown. Trait definitions and units in Table 2.

**Figure S2:** LOD score profiles of P2 based on rMQM results. Trait definitions and units in Table 2. Dashed line indicates genome-wide significance-threshold at 3.8, other thresholds (3.5, 3.6, 3.7) not shown. Trait definitions and units in Table 2.

**Figure S3:** Kruskal-Wallis results (K statistics) for non-parametric traits (STA and SHT). Trait definitions and units in Table 2.

**Figure S4:** Effect of detected QTLs for P1 on traits per allele combinations.

**Figure S5:** Effect of detected QTLs for P2 on traits per allele combinations.

## Author contributions: CRediT

DvM: Writing - original draft, Conceptualization, Methodology, Investigation, Data curation, Formal analysis, Visualization, Funding acquisition, Project administration

JZ: Writing - review and editing, Investigation, Data curation, Formal analysis

AP: Writing - review and editing, Investigation, Data curation

FSL: Writing - review and editing, Software, Data curation

MJ: Writing - review and editing, Funding acquisition, Project administration

FM: Writing - review and editing, Funding acquisition, Project administration, Supervision

All authors read and approved the final version of the manuscript

## Data availability statement

All relevant raw data are provided in supplementary material.

