## Supplementary material for "QTL analysis of essential oil, seed shattering, thousand-grain weight and stalk attachment rate in *Carum carvi*": Figure S4

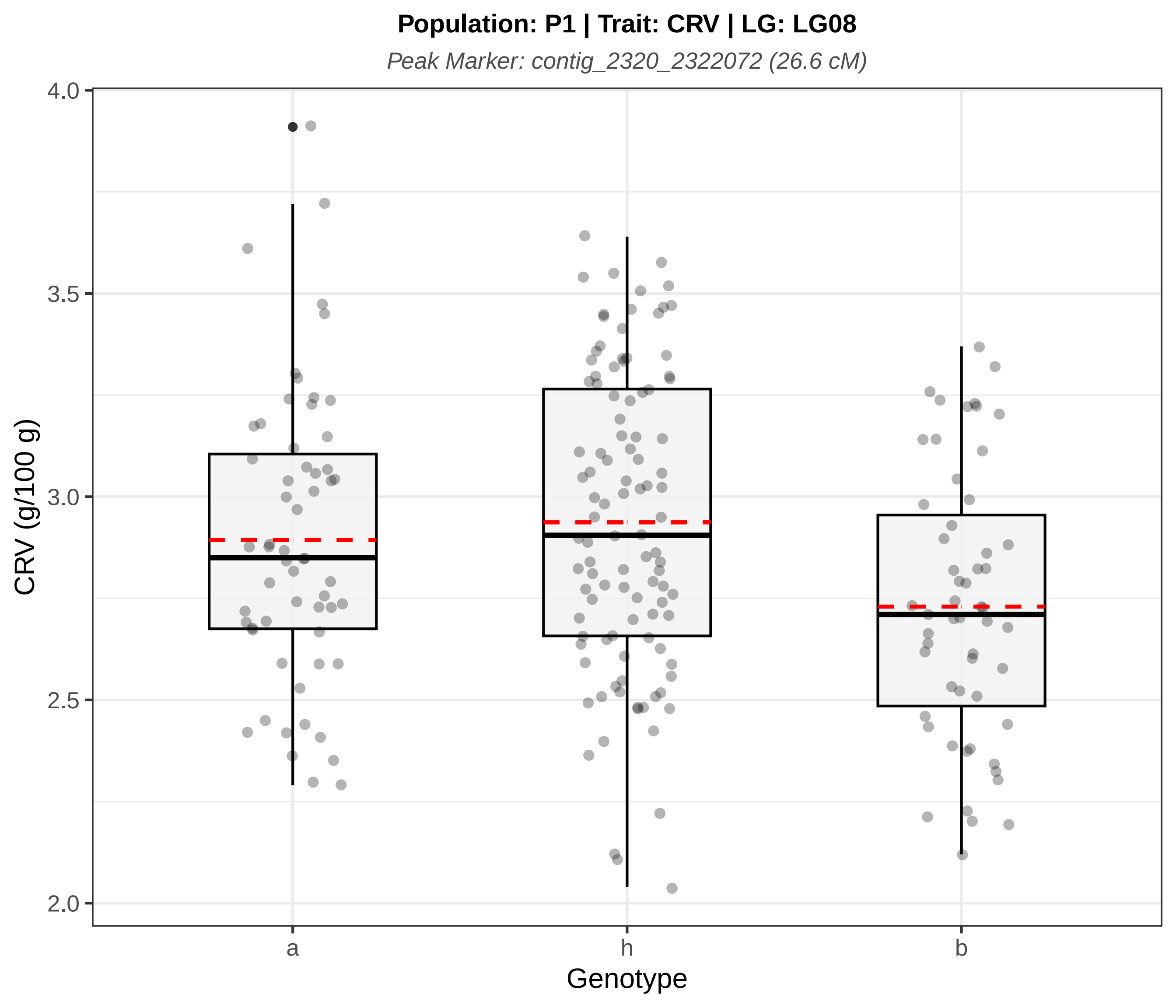
Population: P1 | Trait: CRV | LG: LG08
Peak Marker: contig_2320_2322072 (26.6 cM)
4.0
3.5
3.0
2.5
2.0
CRV (g/100 g)
a
h
b
Genotype

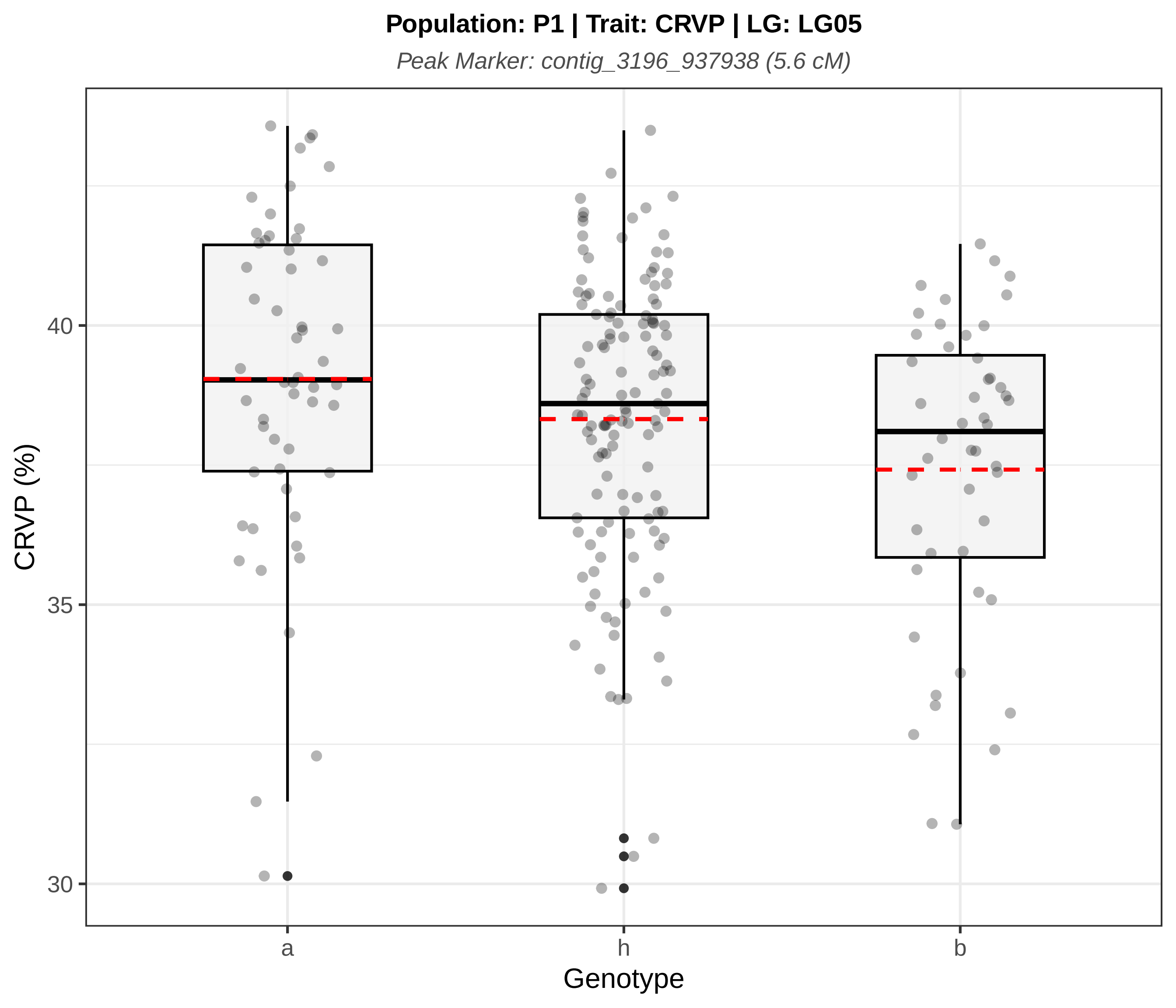

Population: P1 | Trait: CRVP | LG: LG05
Peak Marker: contig_3196_937938 (5.6 cM)
CRVP (%)
40
35
30
a
h
b
Genotype

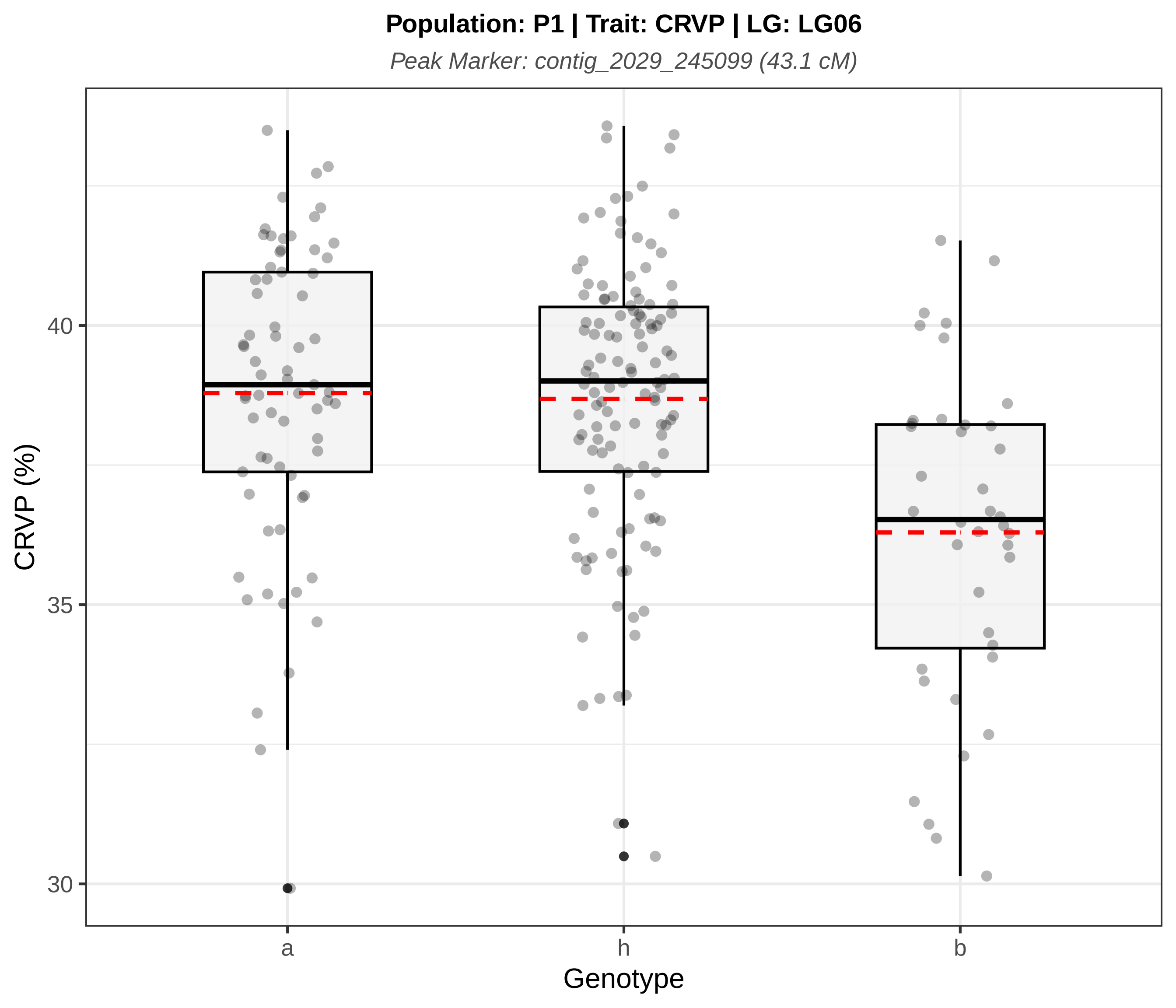

Population: P1 | Trait: CRVP | LG: LG06
Peak Marker: contig_2029_245099 (43.1 cM)
CRVP (%)
40
35
30
a
h
b
Genotype

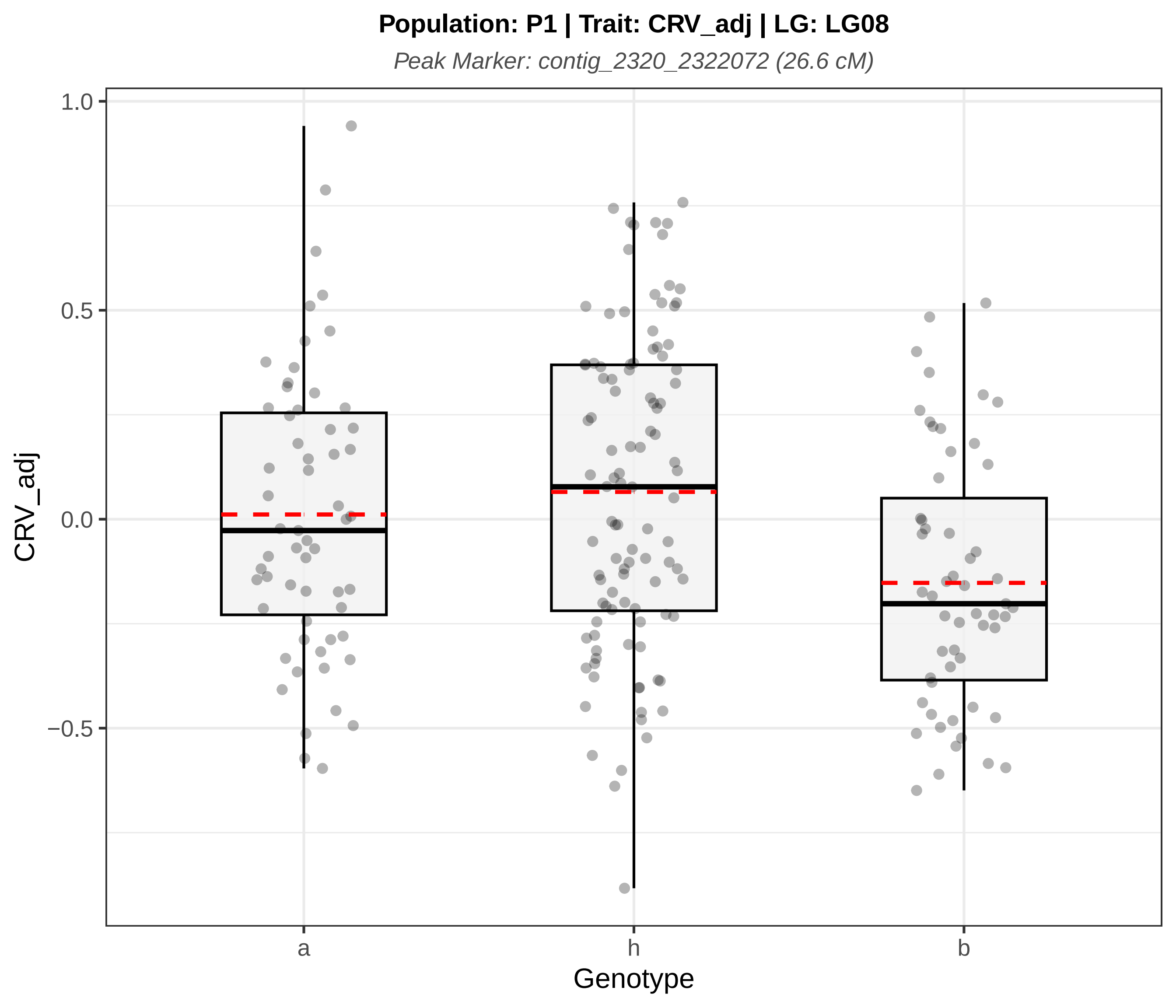

Population: P1 | Trait: CRV_adj | LG: LG08
Peak Marker: contig_2320_2322072 (26.6 cM)
1.0
0.5
0.0
−0.5
CRV_adj
a
h
b
Genotype

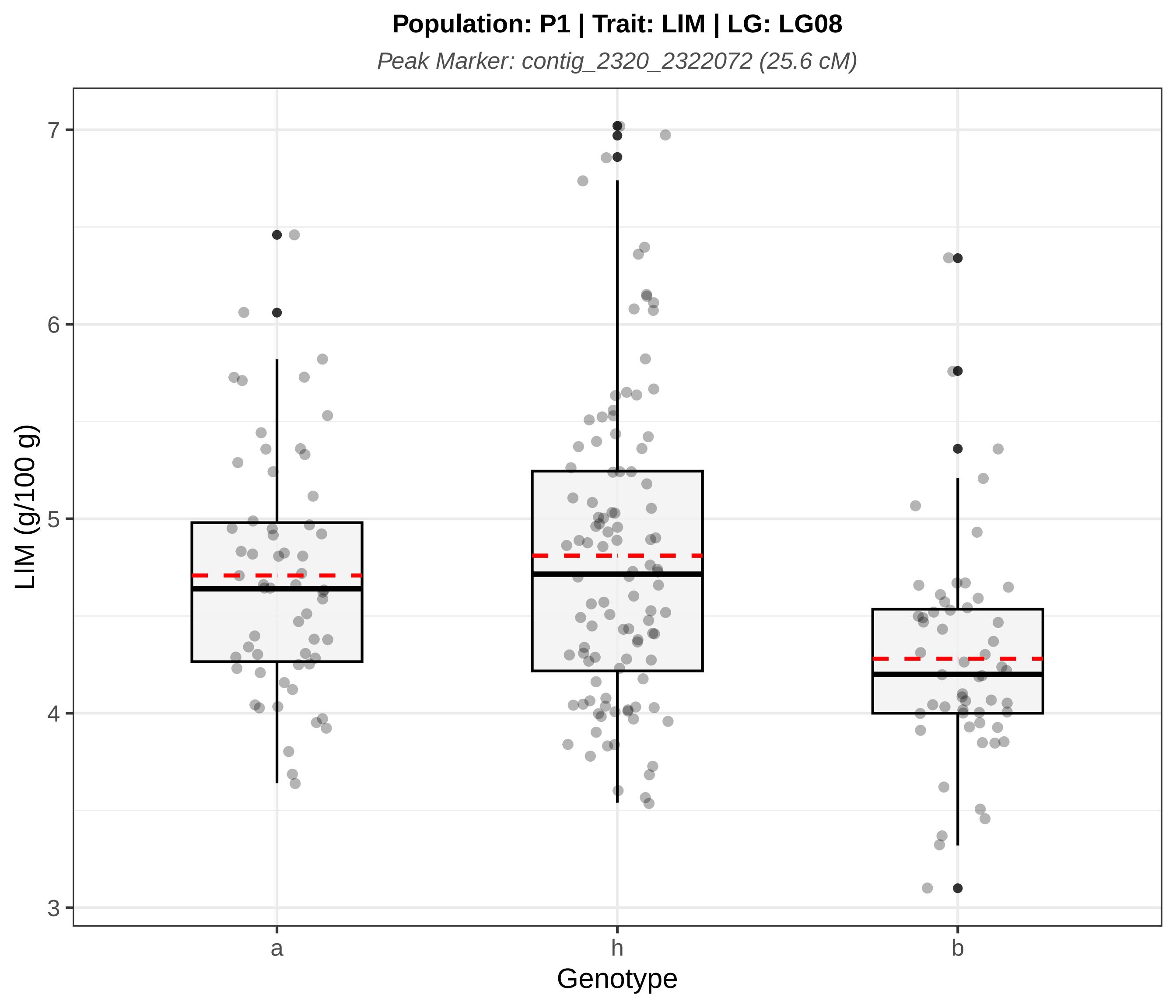

Population: P1 | Trait: LIM | LG: LG08
Peak Marker: contig_2320_2322072 (25.6 cM)
LIM (g/100 g)
7
6
5
4
3
a
h
b
Genotype

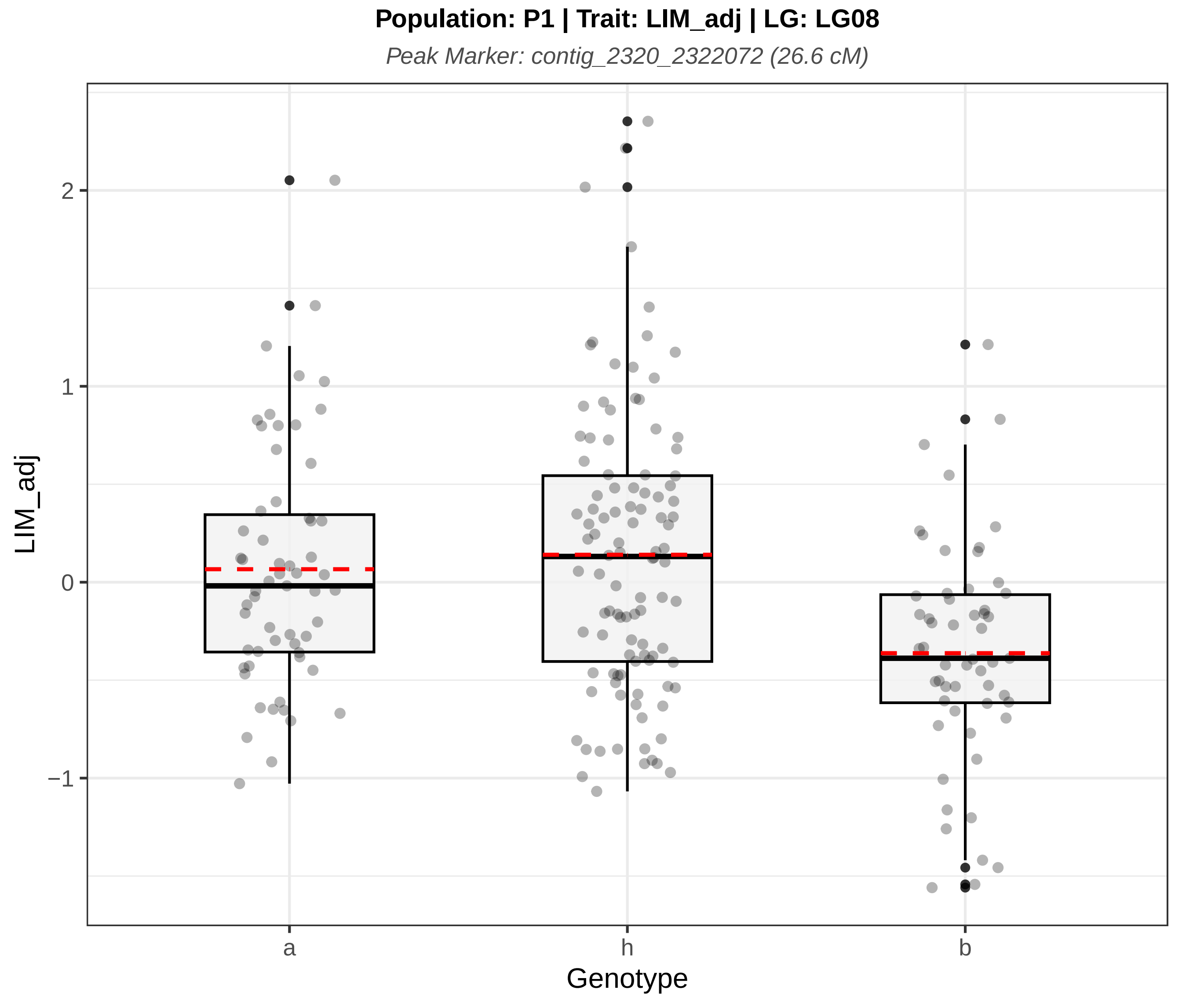

Population: P1 | Trait: LIM_adj | LG: LG08
Peak Marker: contig_2320_2322072 (26.6 cM)
LIM_adj
2
1
0
−1
a
h
b
Genotype

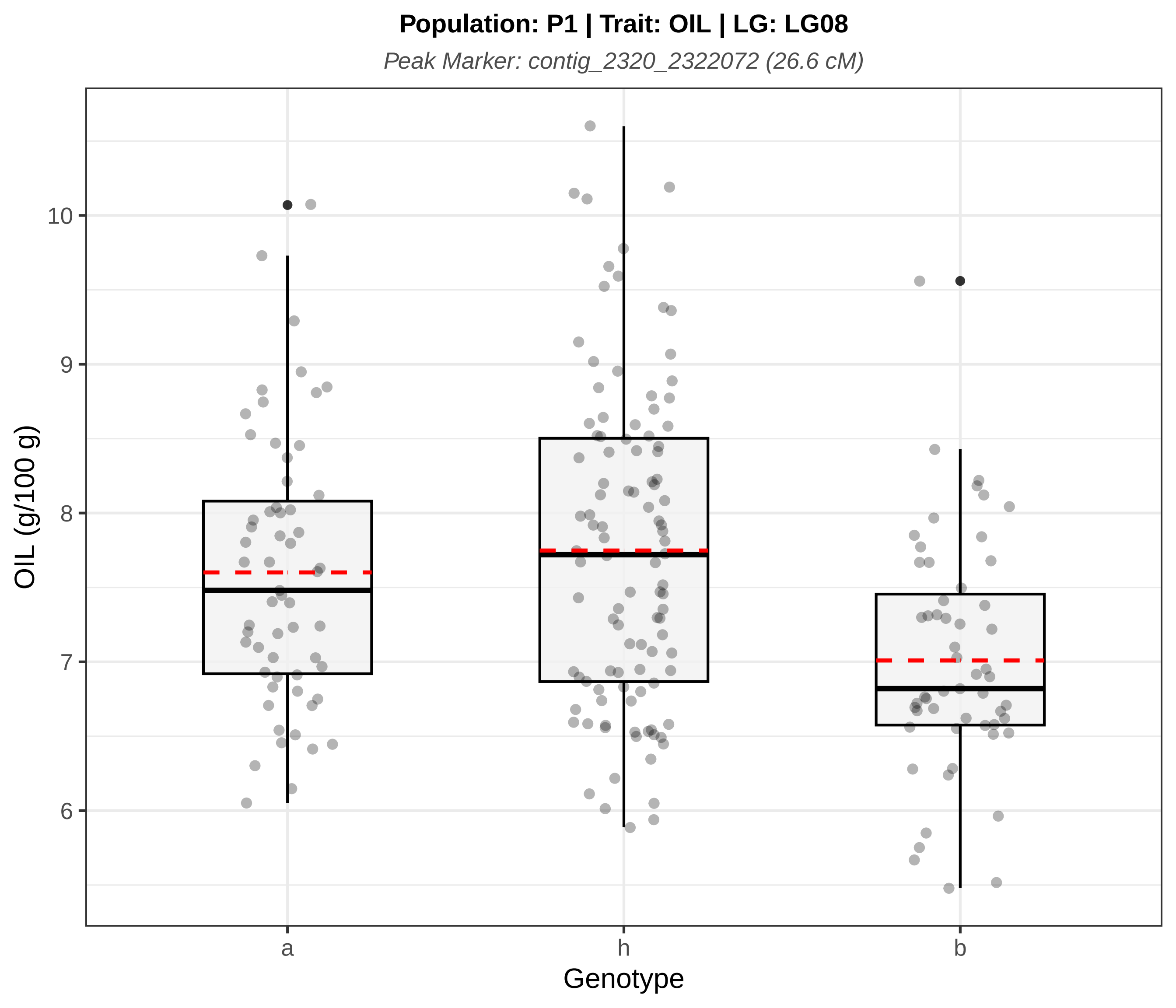
Population: P1 | Trait: OIL | LG: LG08
Peak Marker: contig_2320_2322072 (26.6 cM)
OIL (g/100 g)
10
9
8
7
6
a
h
b
Genotype

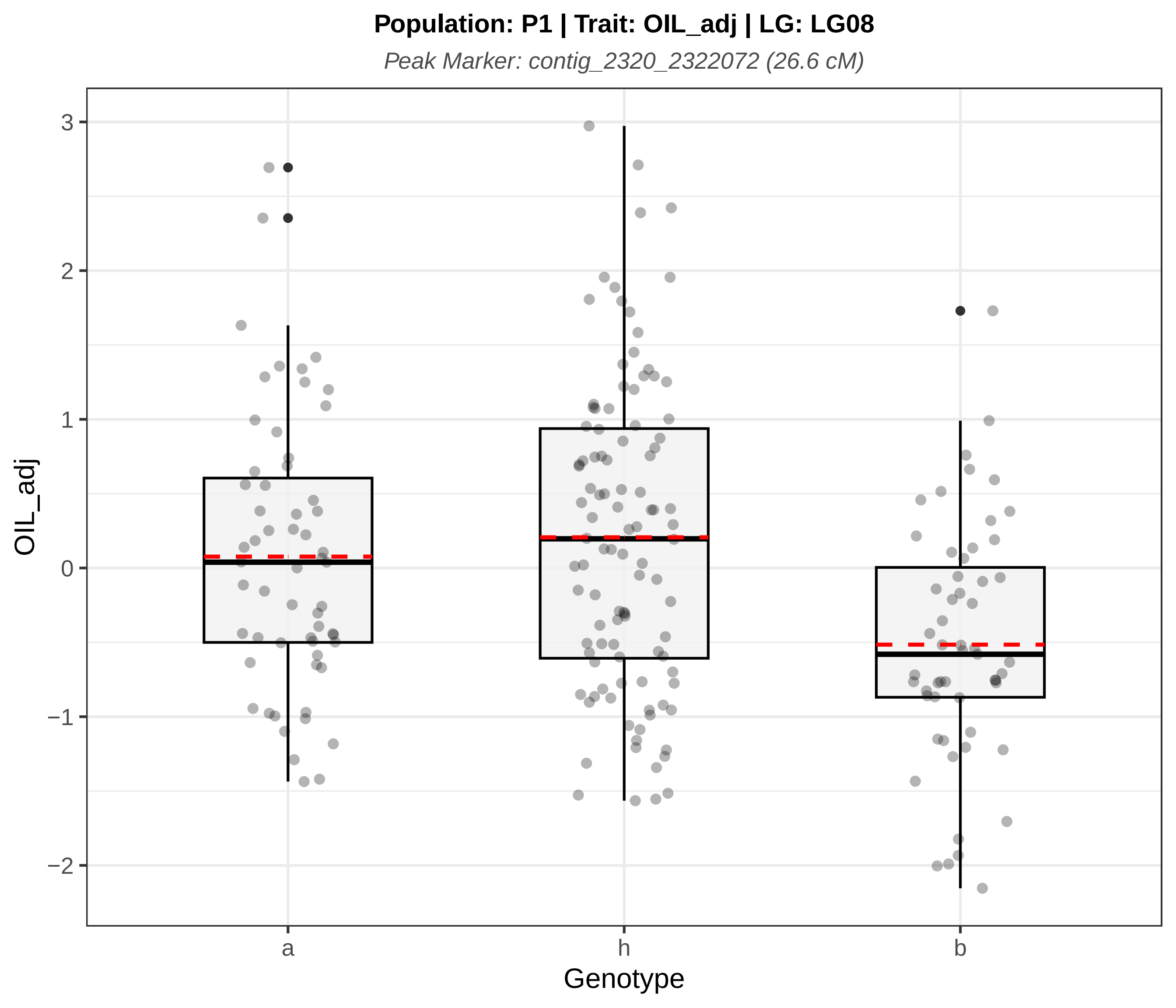
Population: P1 | Trait: OIL_adj | LG: LG08
Peak Marker: contig_2320_2322072 (26.6 cM)
3
2
1
0
−1
−2
OIL_adj
a
h
b
Genotype

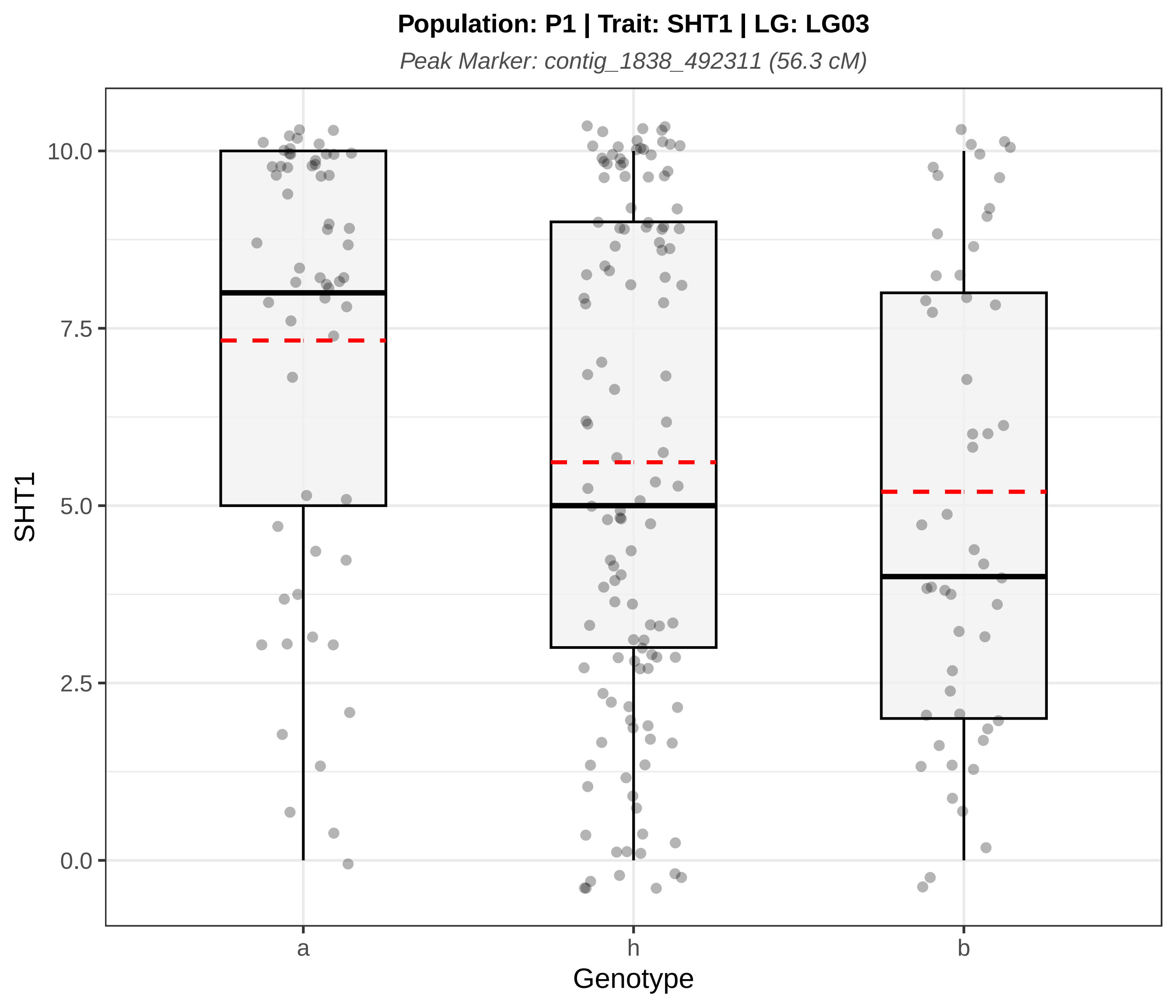

Population: P1 | Trait: SHT1 | LG: LG03
Peak Marker: contig_1838_492311 (56.3 cM)
SHT1
0.0
2.5
5.0
7.5
10.0
a
h
b
Genotype

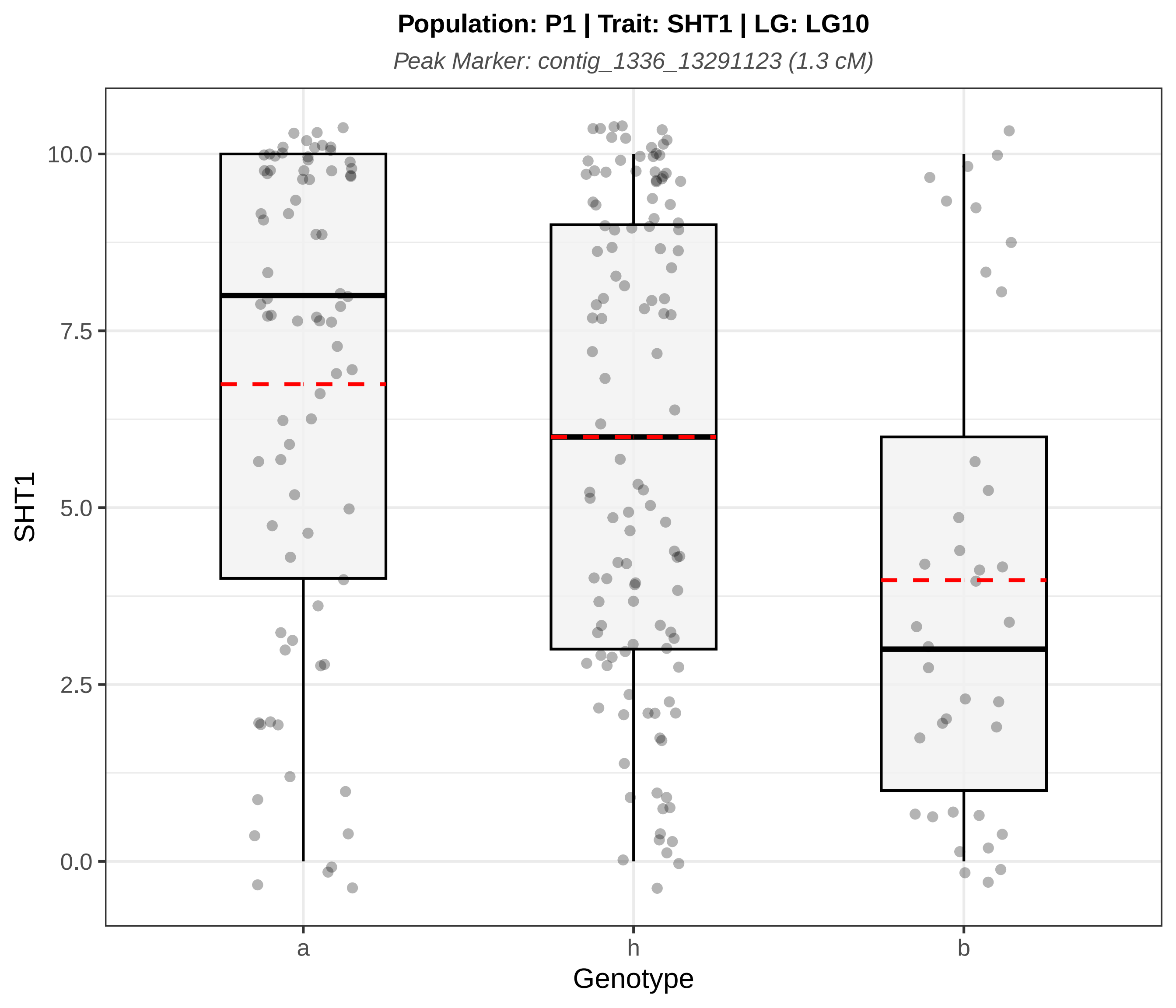
Population: P1 | Trait: SHT1 | LG: LG10
Peak Marker: contig_1336_13291123 (1.3 cM)
SHT1
10.0
7.5
5.0
2.5
0.0
a
h
b
Genotype

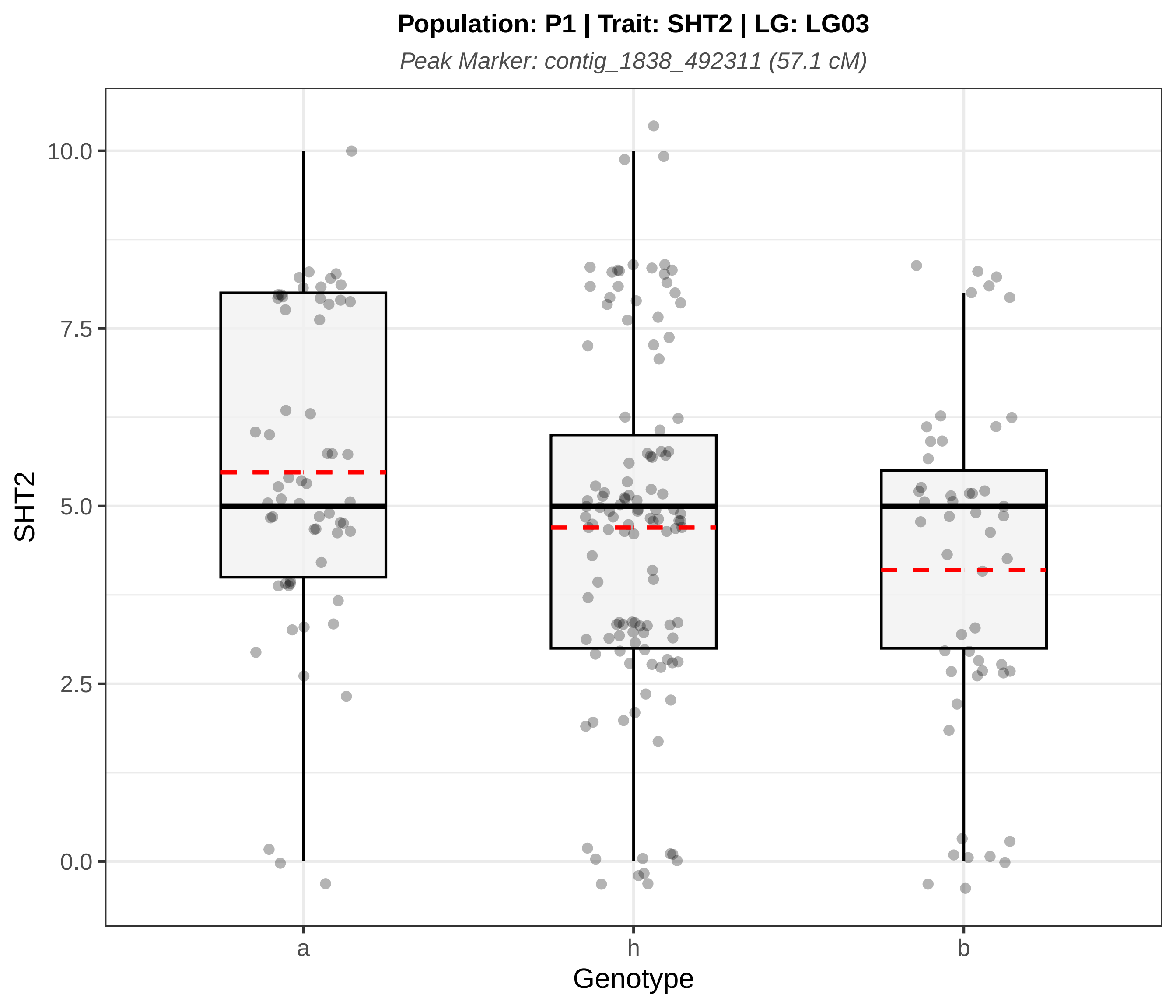

Population: P1 | Trait: SHT2 | LG: LG03
Peak Marker: contig_1838_492311 (57.1 cM)
SHT2
10.0
7.5
5.0
2.5
0.0
a
h
b
Genotype

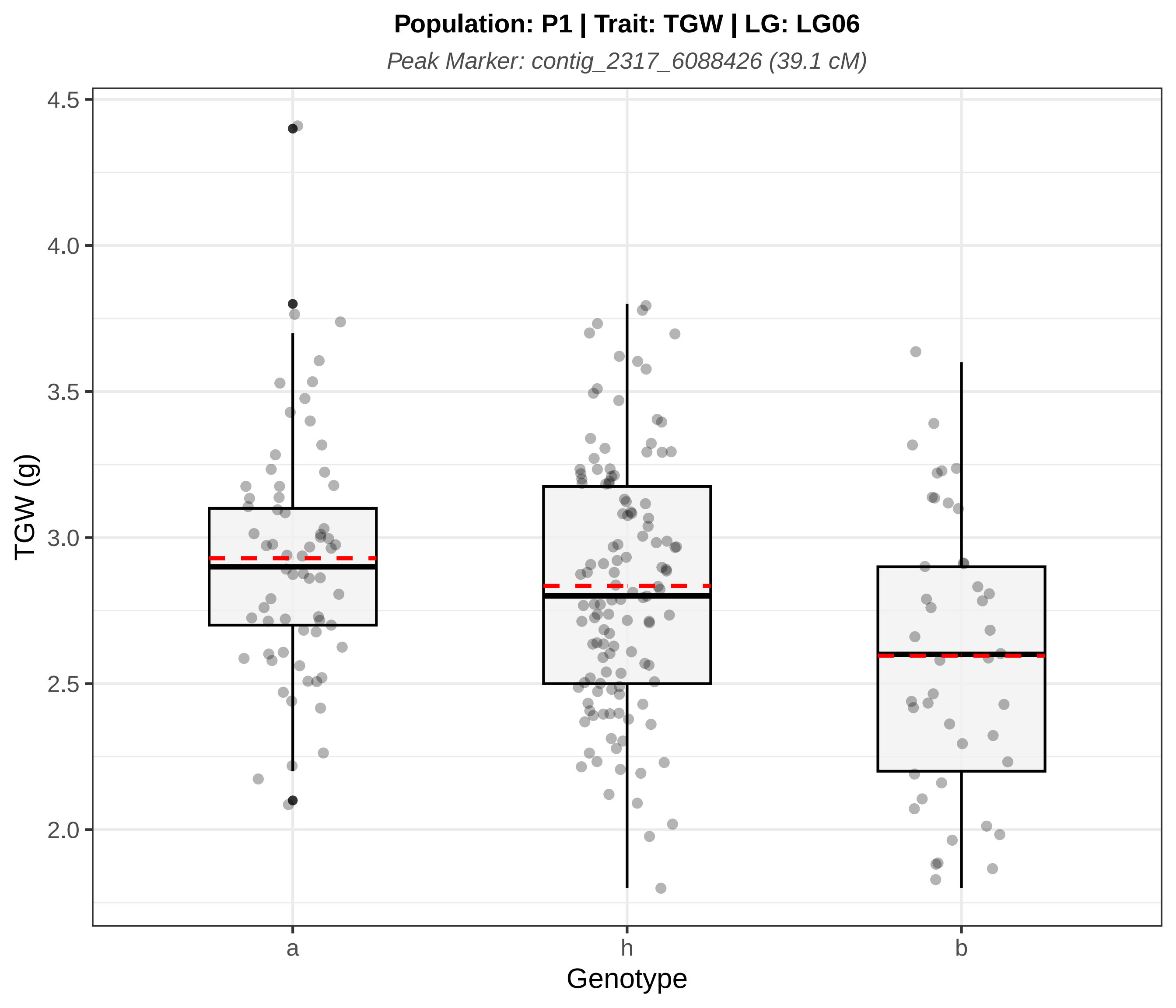

Population: P1 | Trait: TGW | LG: LG06
Peak Marker: contig_2317_6088426 (39.1 cM)
TGW (g)
4.5
4.0
3.5
3.0
2.5
2.0
a
h
b
Genotype

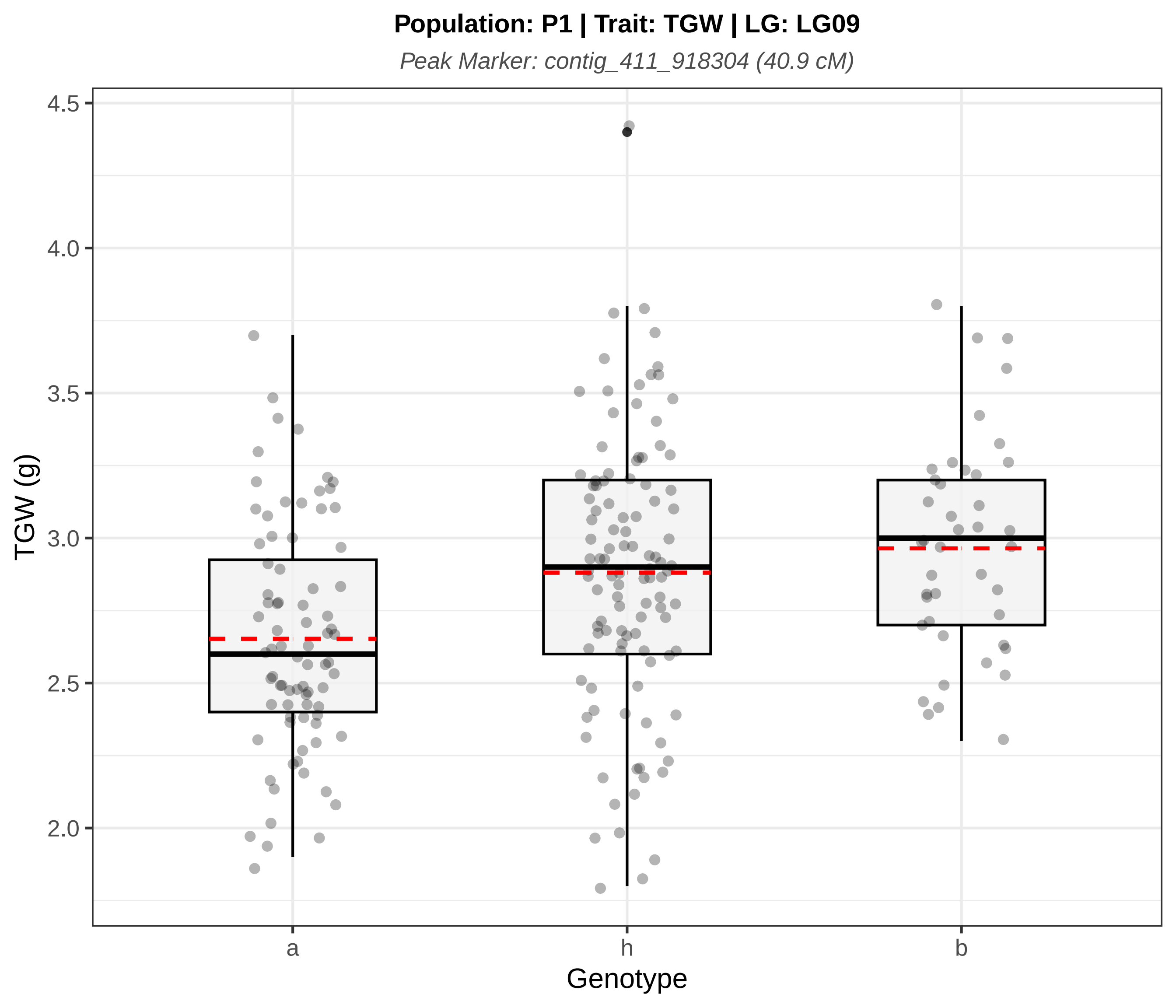
Population: P1 | Trait: TGW | LG: LG09
Peak Marker: contig_411_918304 (40.9 cM)
TGW (g)
4.5
4.0
3.5
3.0
2.5
2.0
a
h
b
Genotype

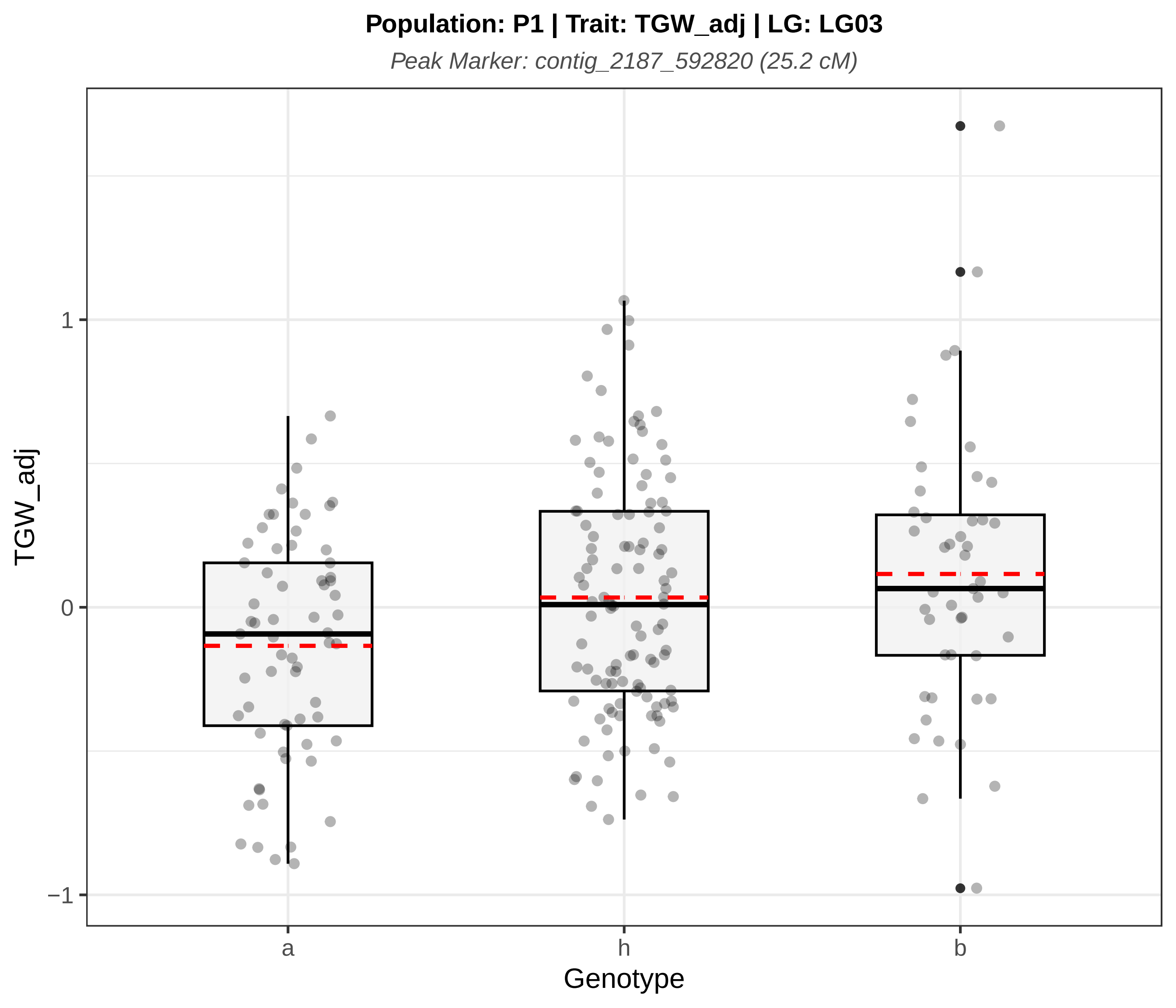
Population: P1 | Trait: TGW_adj | LG: LG03
Peak Marker: contig_2187_592820 (25.2 cM)
TGW_adj
1
0
−1
a
h
b
Genotype

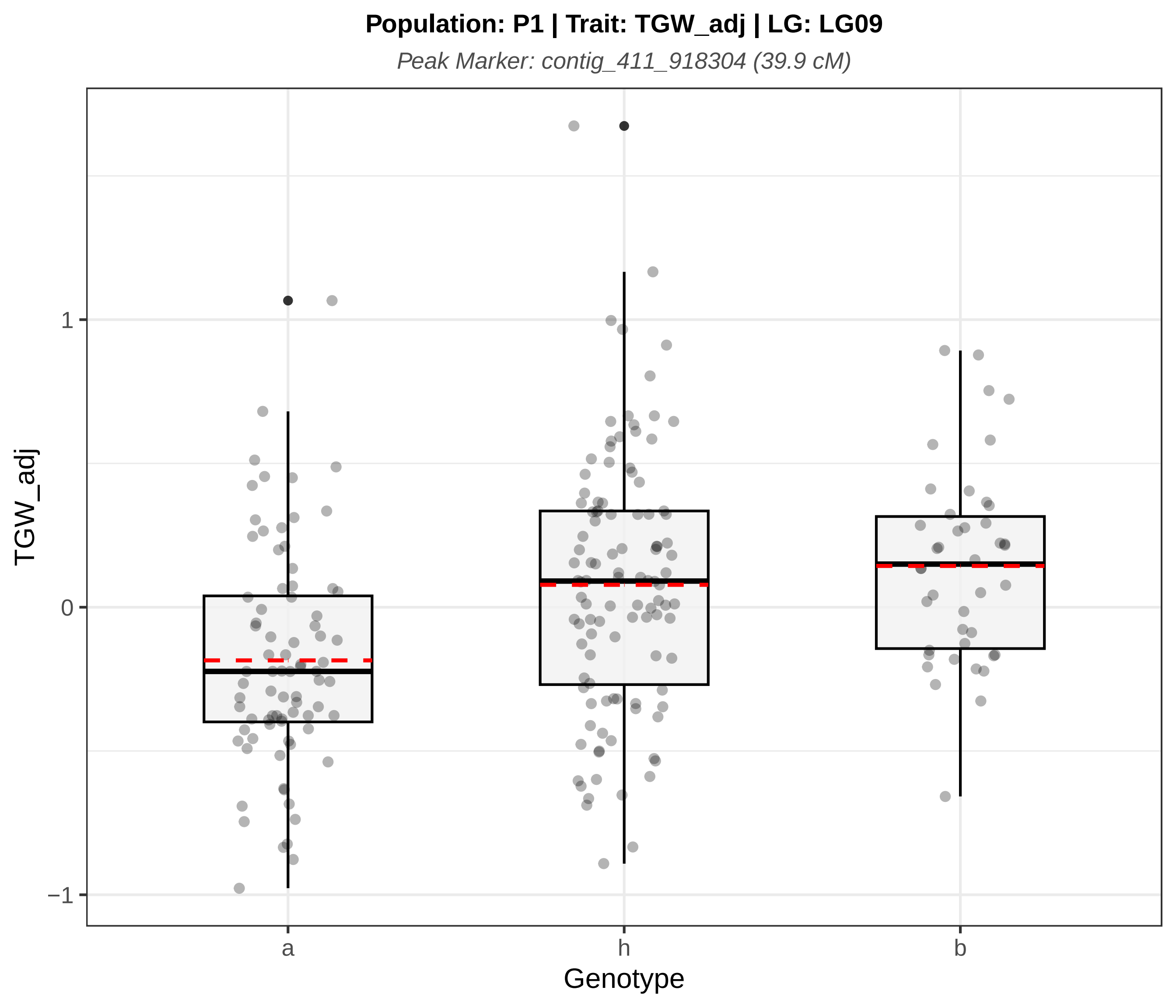

Population: P1 | Trait: TGW_adj | LG: LG09
Peak Marker: contig_411_918304 (39.9 cM)
TGW_adj
1
0
−1
a
h
b
Genotype
