## Supplementary material for "QTL analysis of essential oil, seed shattering, thousand-grain weight and stalk attachment rate in *Carum carvi*": Figure S5

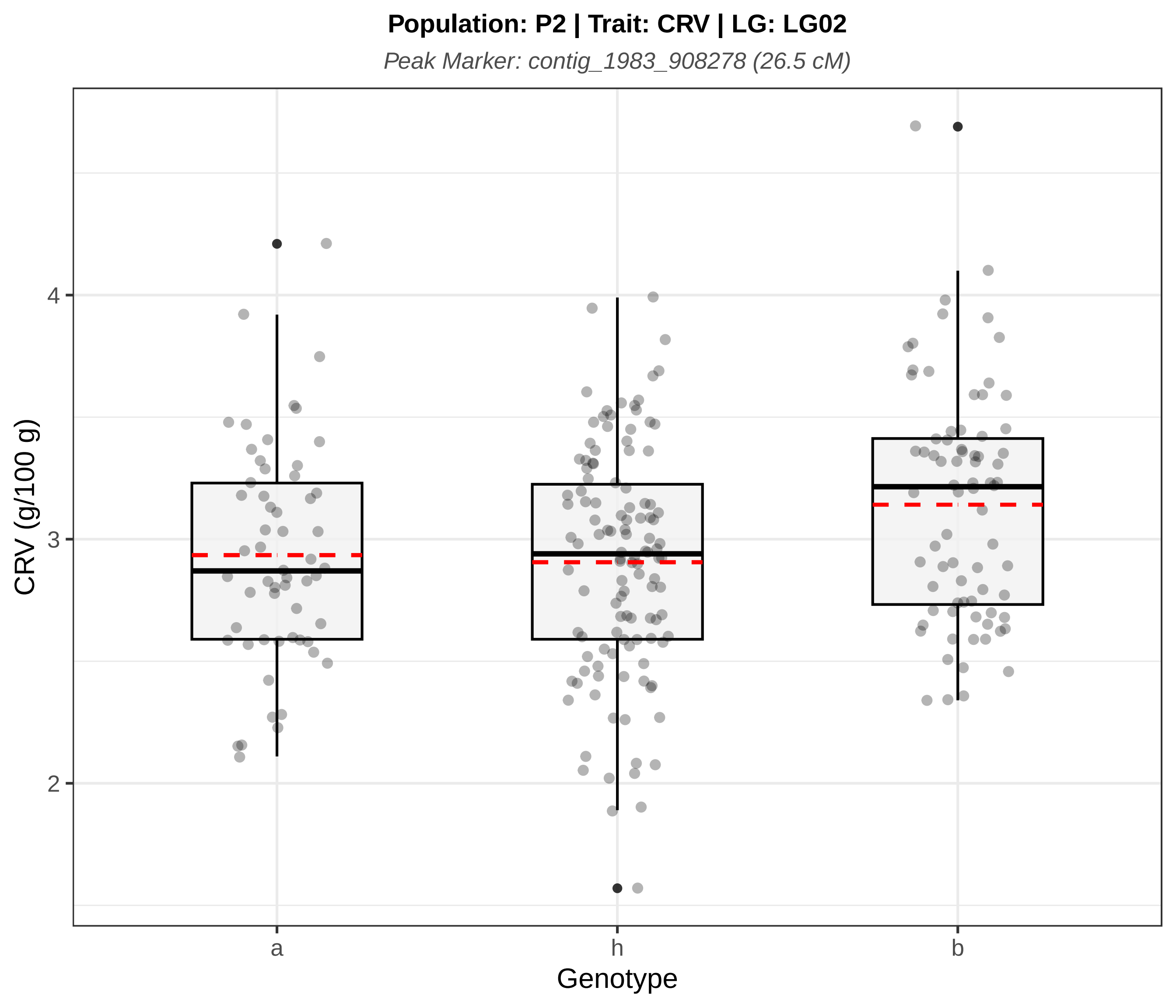
Population: P2 | Trait: CRV | LG: LG02
Peak Marker: contig_1983_908278 (26.5 cM)
CRV (g/100 g)
4
3
2
a
h
b
Genotype

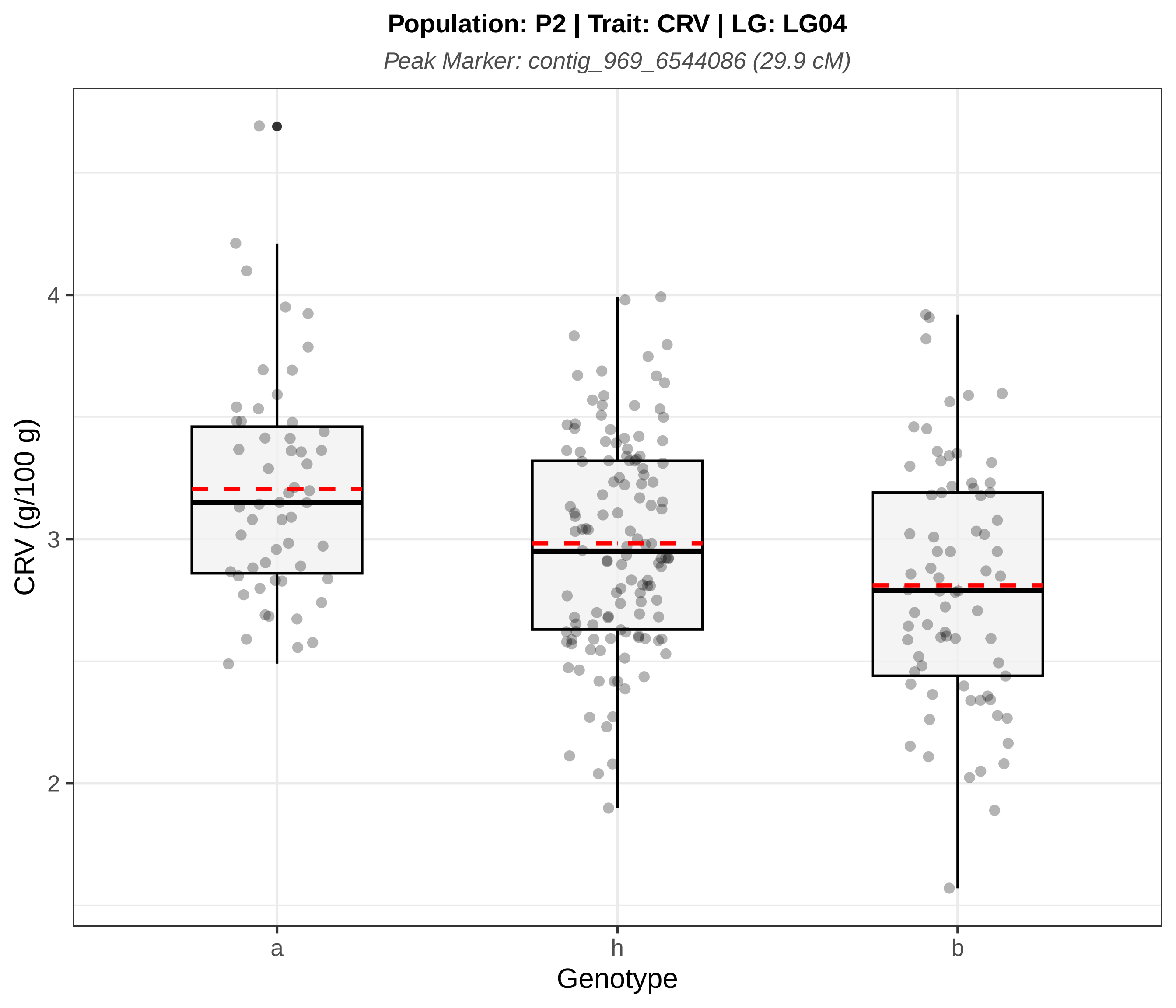
Population: P2 | Trait: CRV | LG: LG04
Peak Marker: contig_969_6544086 (29.9 cM)
CRV (g/100 g)
4
3
2
a
h
b
Genotype

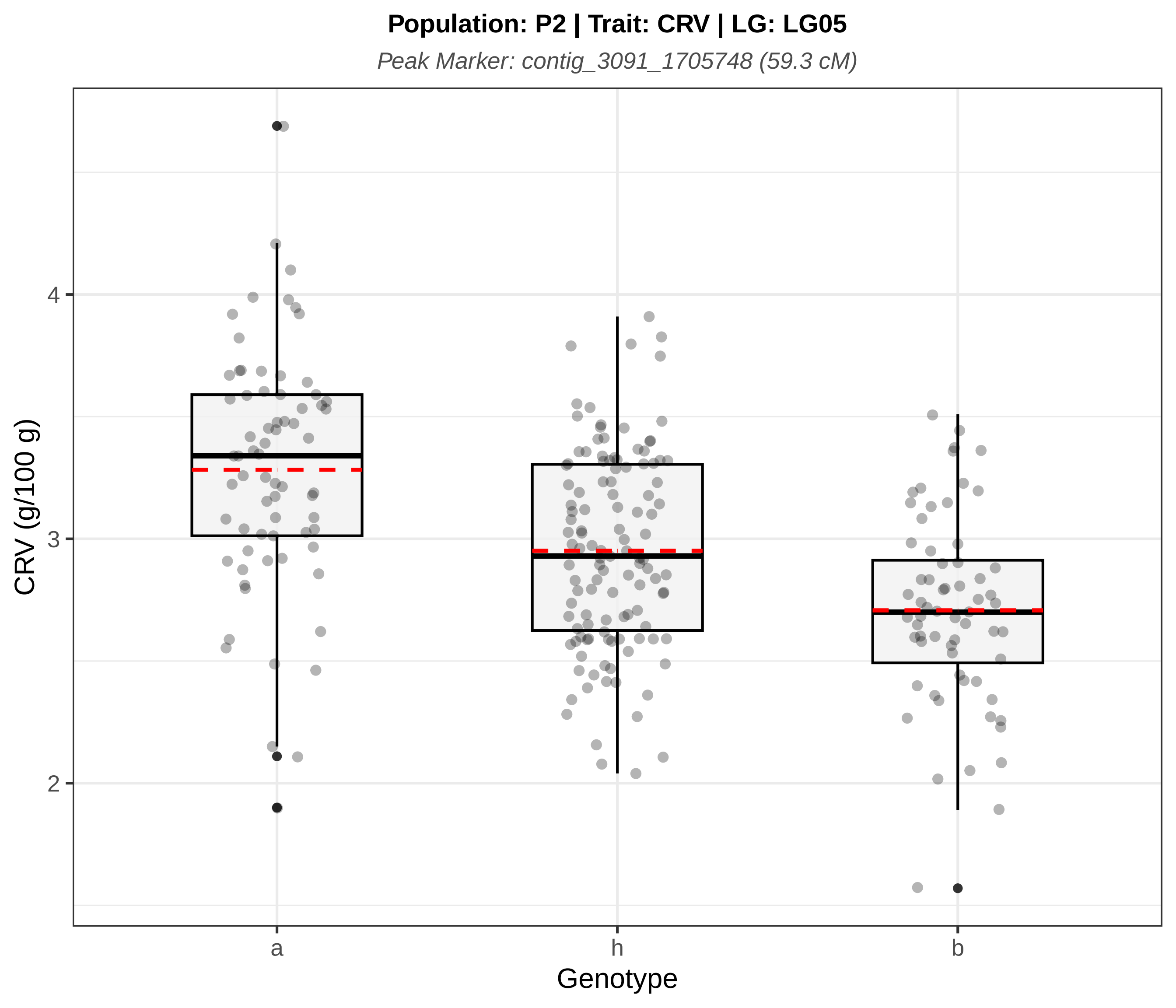

Population: P2 | Trait: CRV | LG: LG05
Peak Marker: contig_3091_1705748 (59.3 cM)
CRV (g/100 g)
4
3
2
a
h
b
Genotype

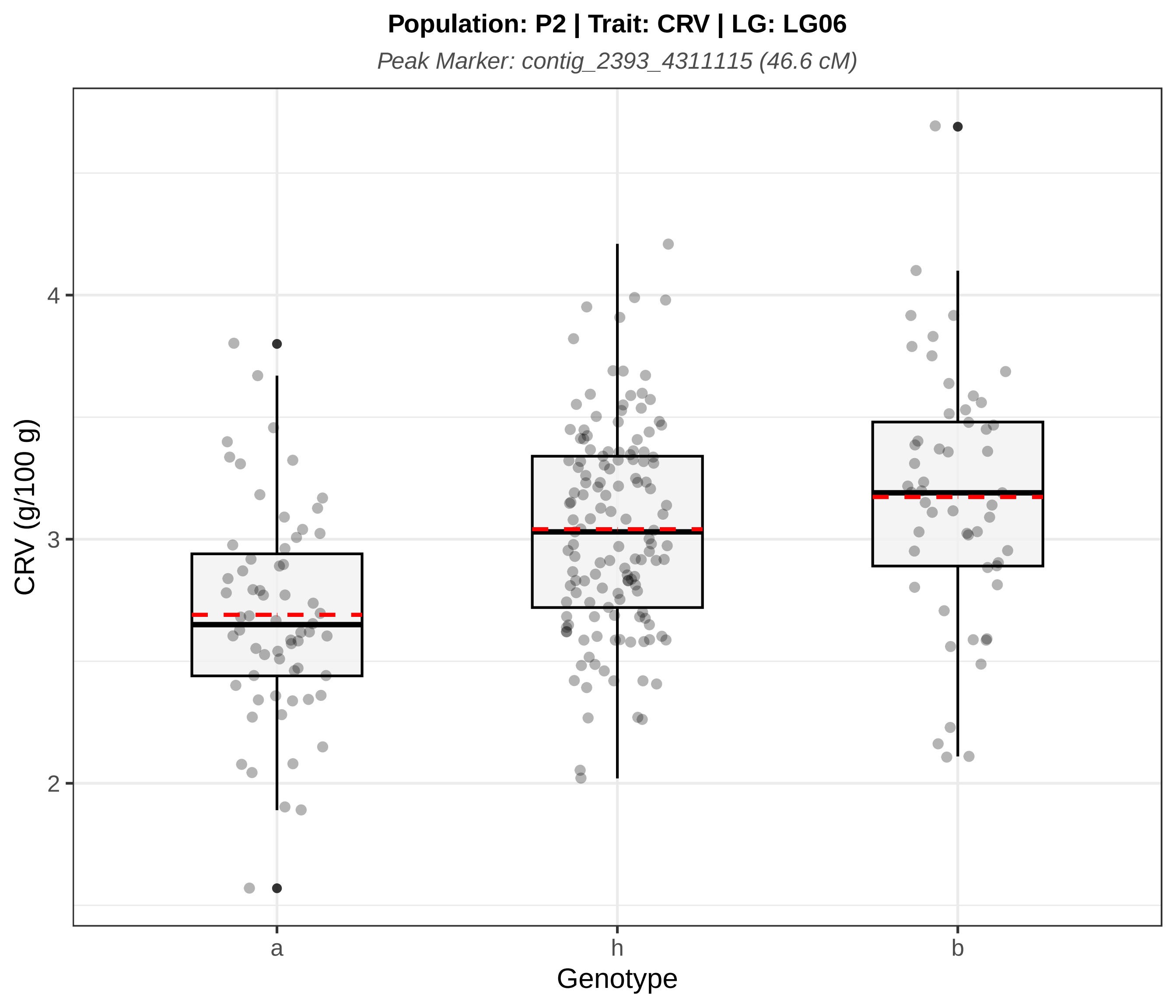

Population: P2 | Trait: CRV | LG: LG06
Peak Marker: contig_2393_4311115 (46.6 cM)
CRV (g/100 g)
4
3
2
a
h
b
Genotype

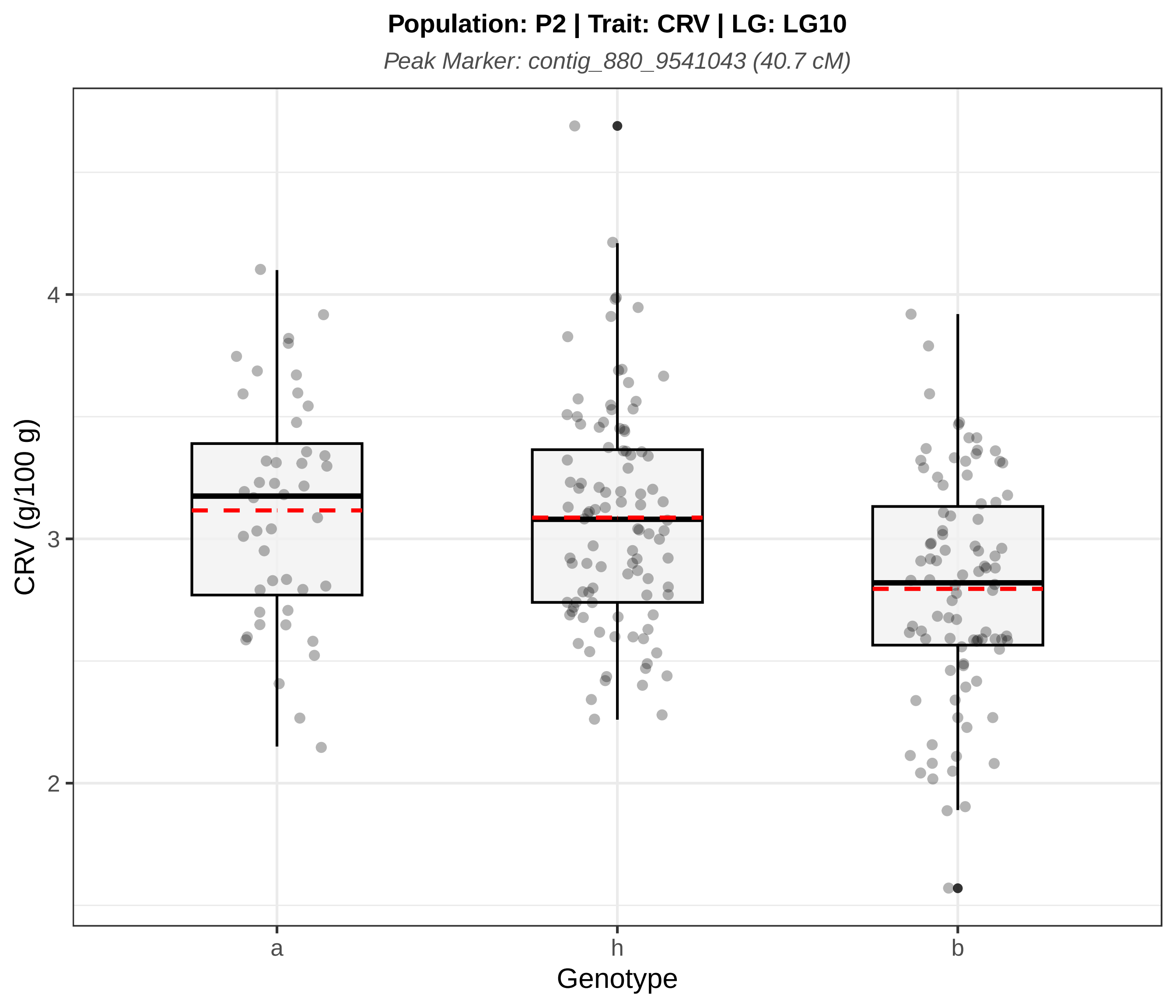
Population: P2 | Trait: CRV | LG: LG10
Peak Marker: contig_880_9541043 (40.7 cM)
CRV (g/100 g)
4
3
2
a
h
b
Genotype

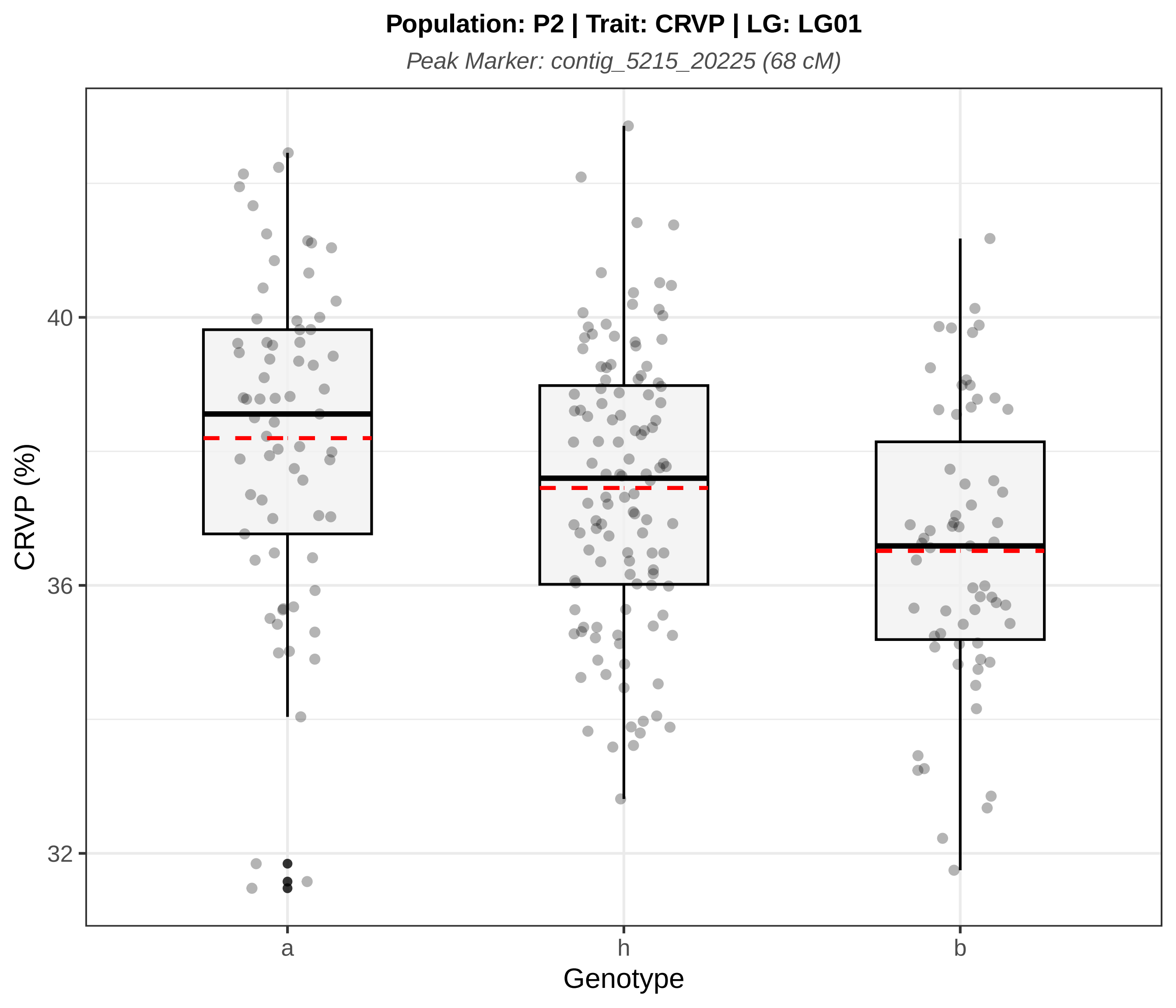
Population: P2 | Trait: CRVP | LG: LG01
Peak Marker: contig_5215_20225 (68 cM)
CRVP (%)
40
36
32
a
h
b
Genotype

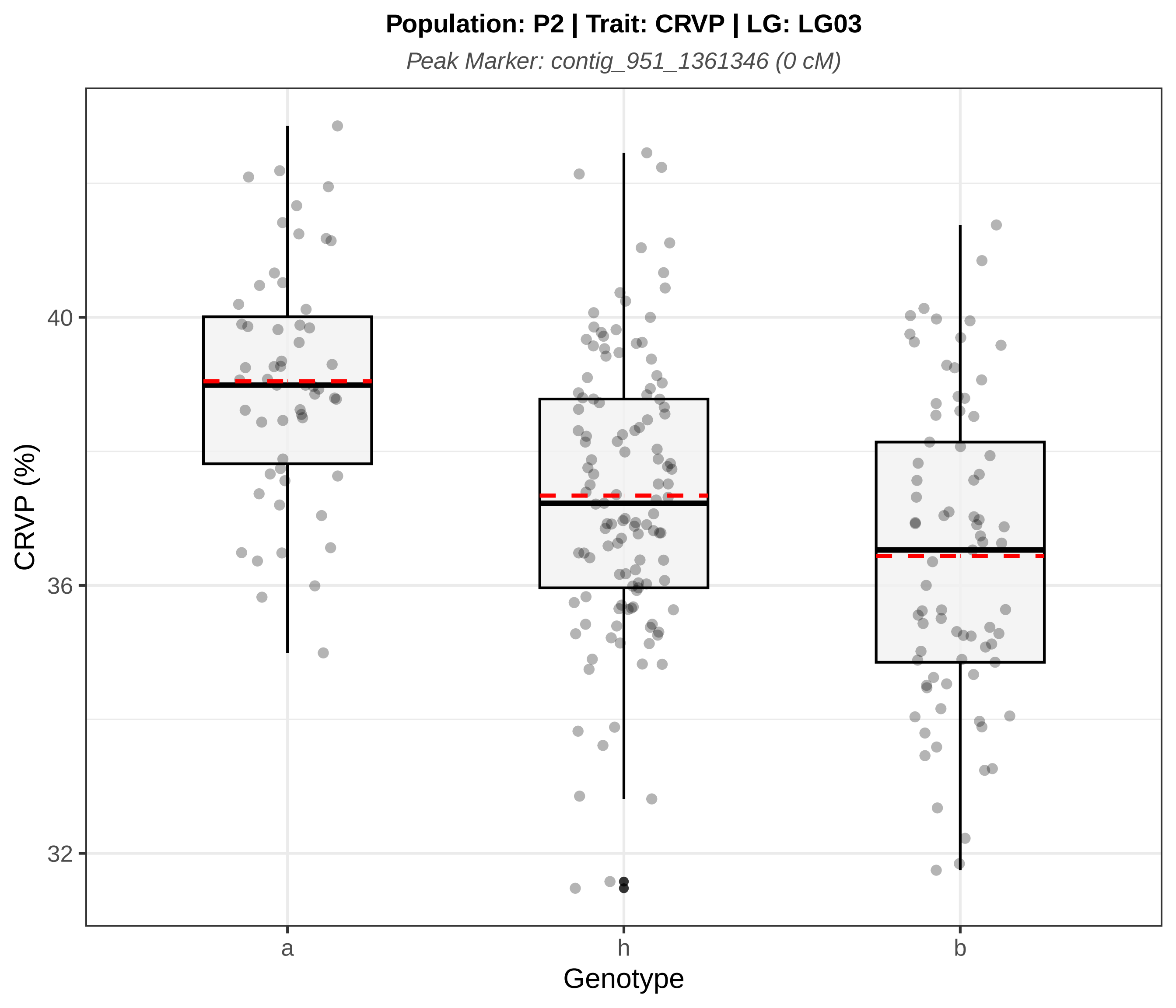
Population: P2 | Trait: CRVP | LG: LG03
Peak Marker: contig_951_1361346 (0 cM)
CRVP (%)
40
36
32
a
h
b
Genotype

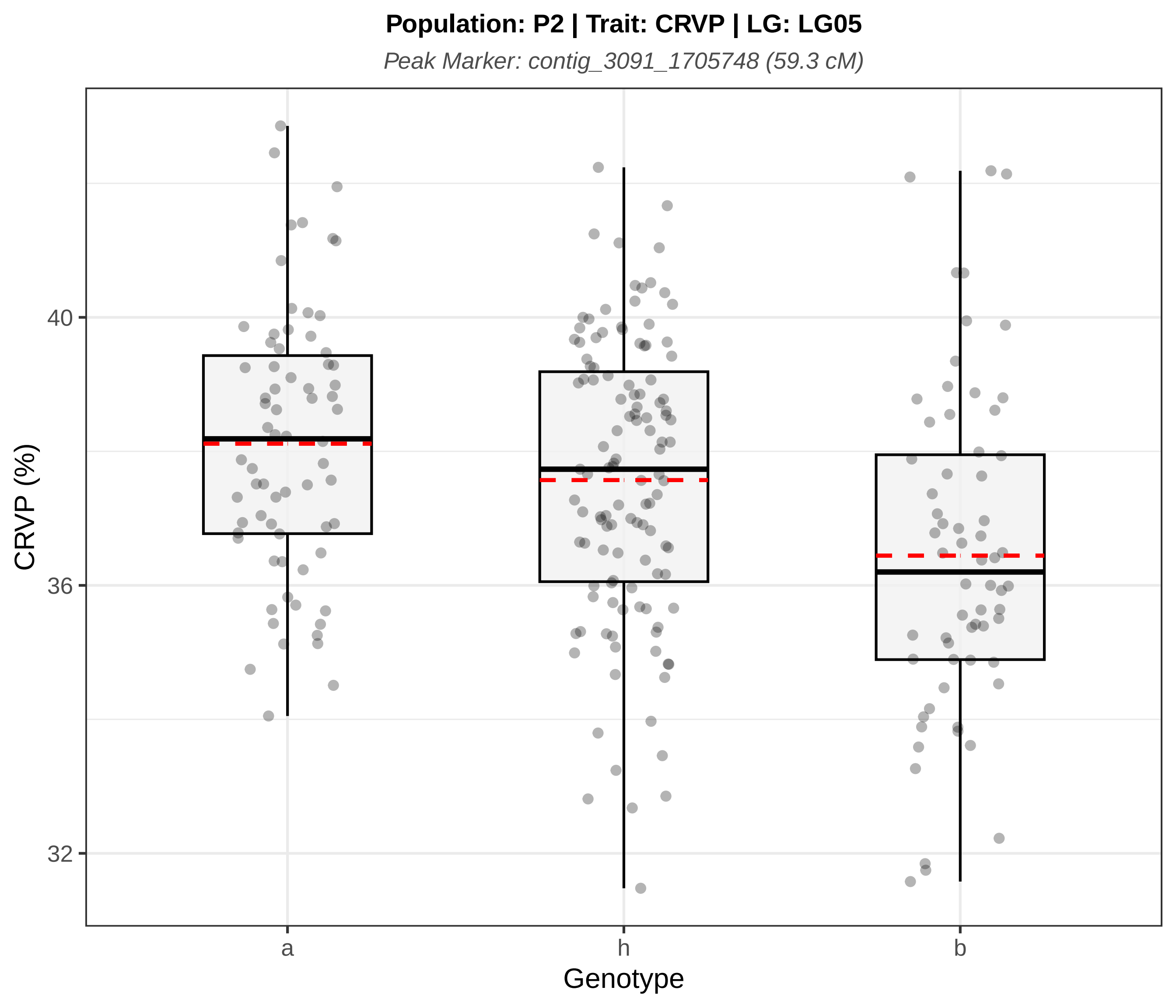

Population: P2 | Trait: CRVP | LG: LG05
Peak Marker: contig_3091_1705748 (59.3 cM)
CRVP (%)
40
36
32
a
h
b
Genotype

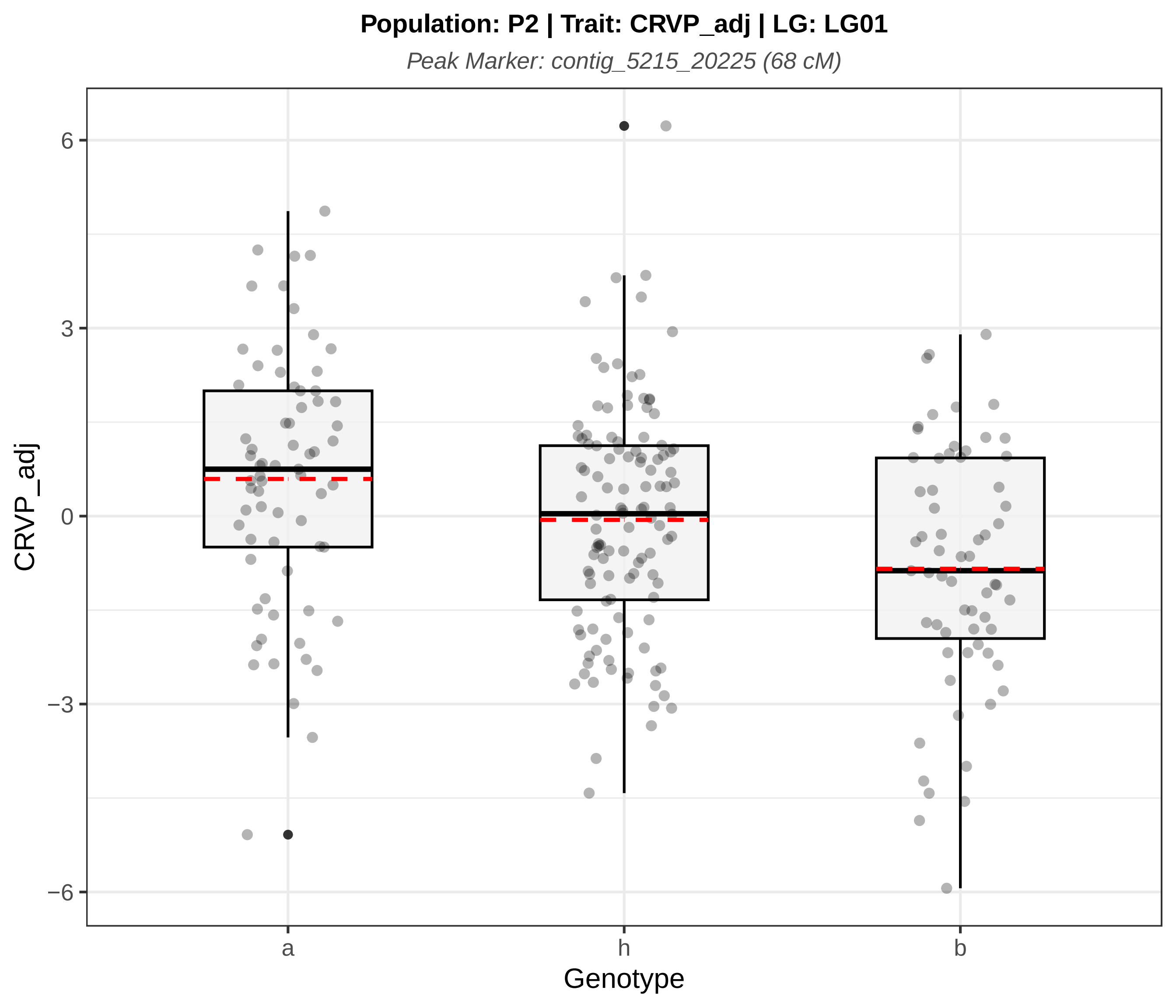
Population: P2 | Trait: CRVP_adj | LG: LG01
Peak Marker: contig_5215_20225 (68 cM)
CRVP_adj
6
3
0
−3
−6
a
h
b
Genotype

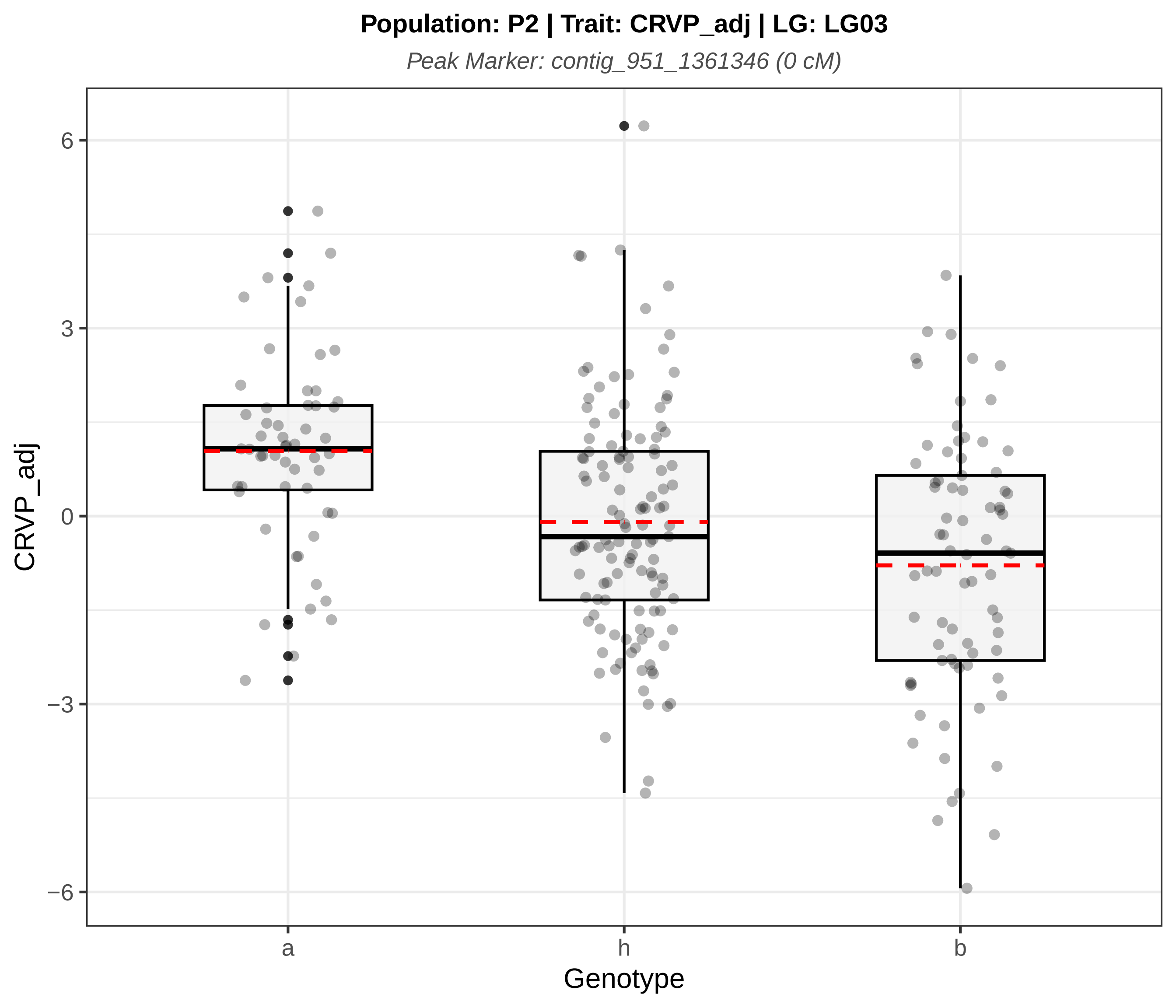

Population: P2 | Trait: CRVP_adj | LG: LG03
Peak Marker: contig_951_1361346 (0 cM)
6
3
0
−3
−6
CRVP_adj
a
h
b
Genotype

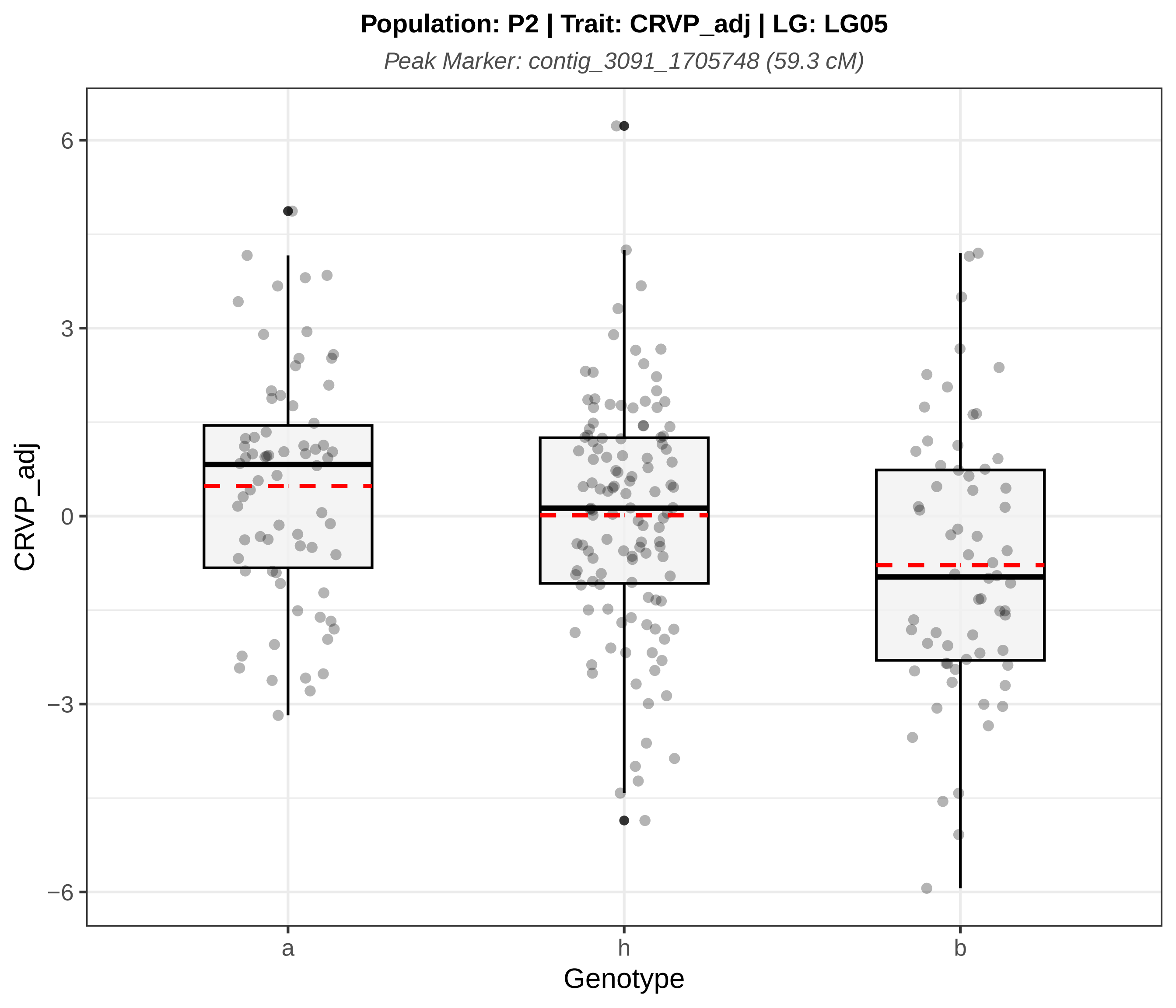
Population: P2 | Trait: CRVP_adj | LG: LG05
Peak Marker: contig_3091_1705748 (59.3 cM)
CRVP_adj
6
3
0
−3
−6
a
h
b
Genotype

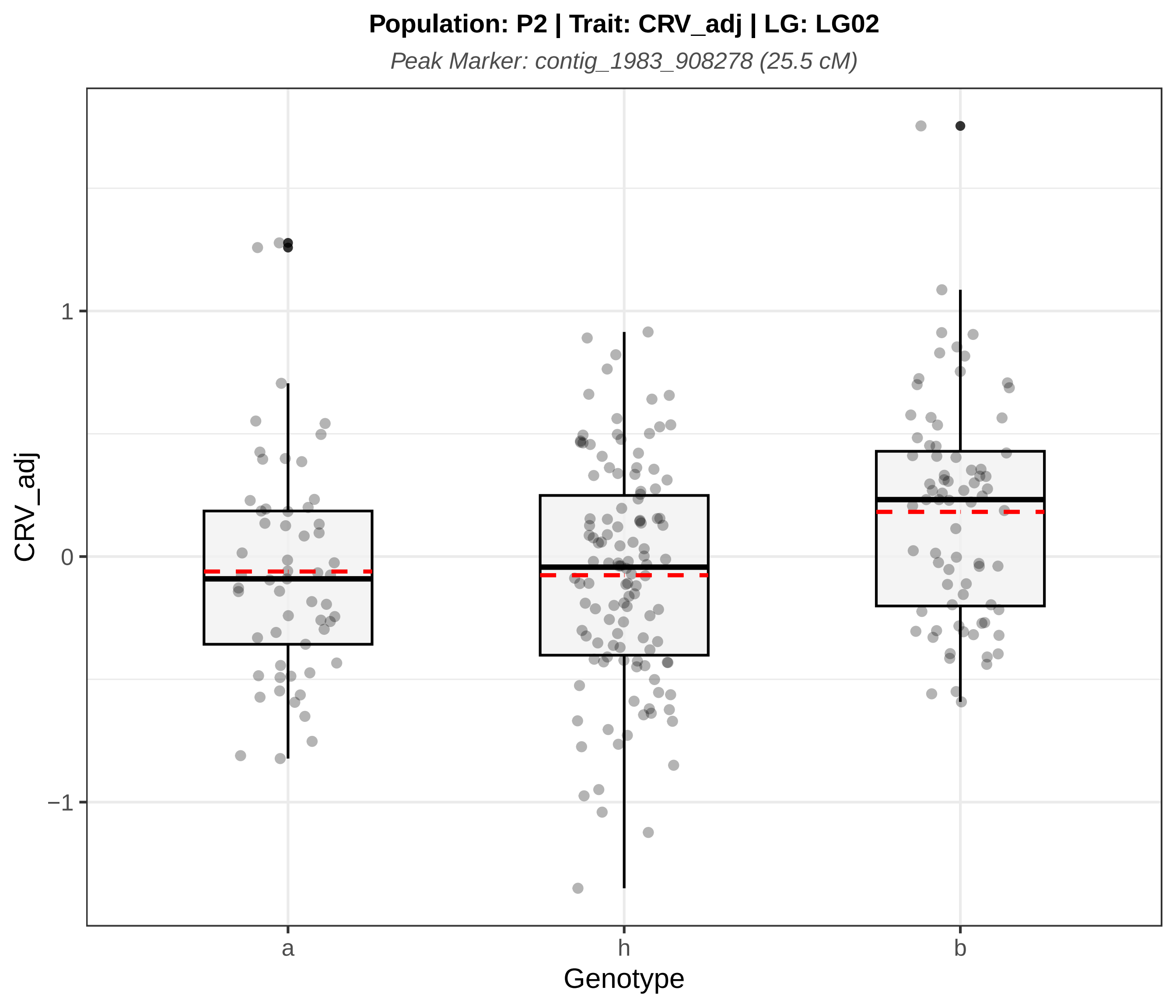

Population: P2 | Trait: CRV_adj | LG: LG02
Peak Marker: contig_1983_908278 (25.5 cM)
CRV_adj
1
0
−1
a
h
b
Genotype

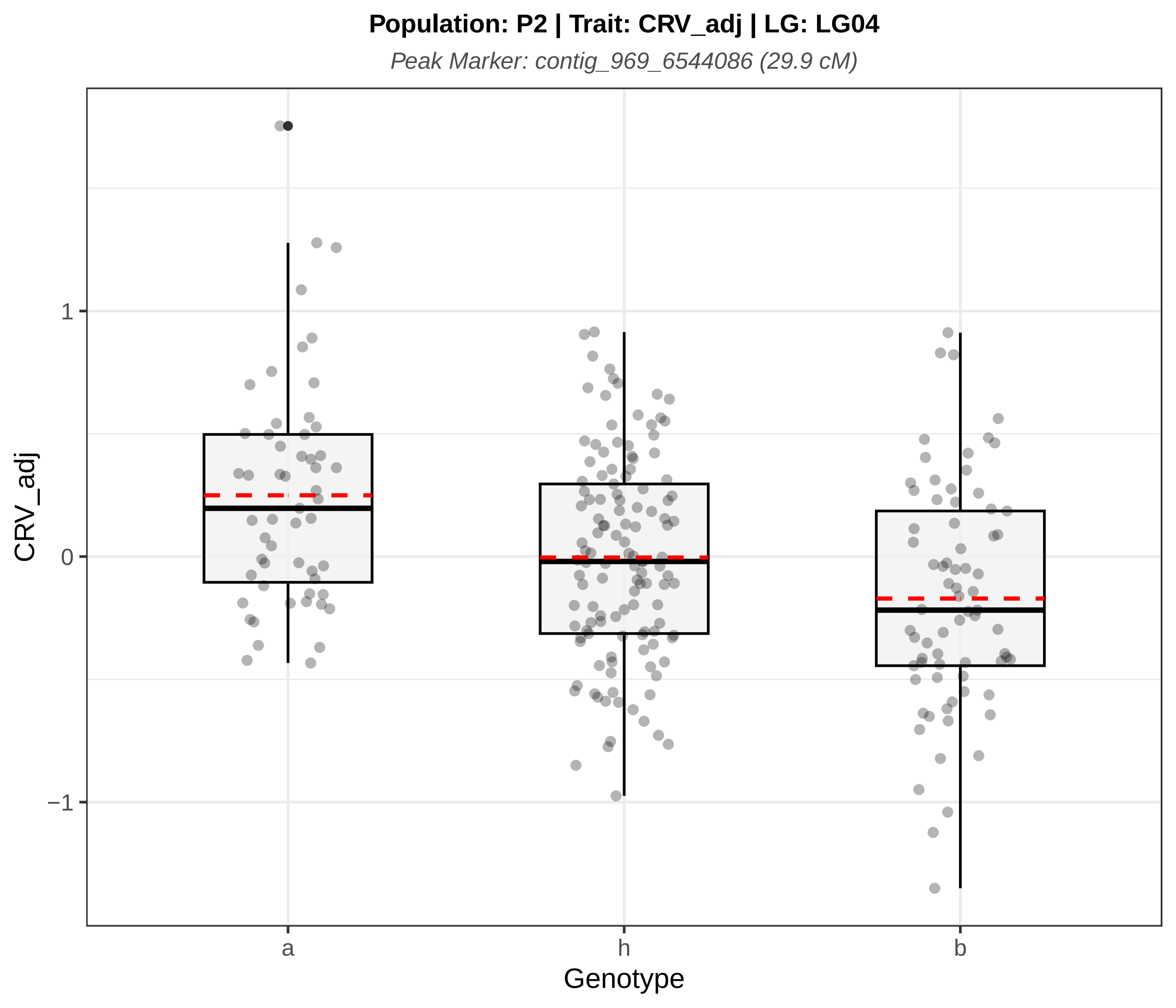
Population: P2 | Trait: CRV_adj | LG: LG04
Peak Marker: contig_969_6544086 (29.9 cM)
CRV_adj
1
0
−1
a
h
b
Genotype

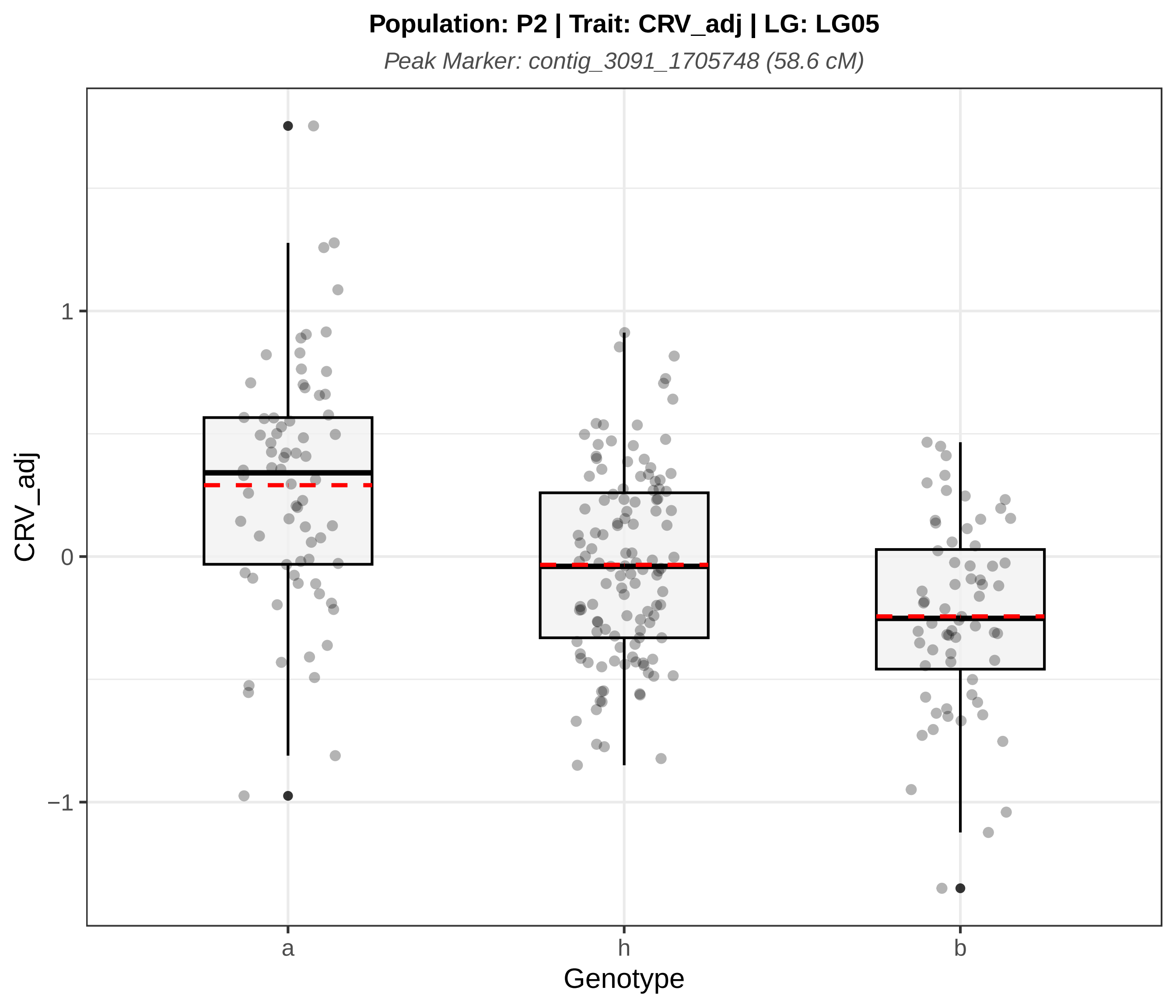
Population: P2 | Trait: CRV_adj | LG: LG05
Peak Marker: contig_3091_1705748 (58.6 cM)
CRV_adj
1
0
−1
a
h
b
Genotype

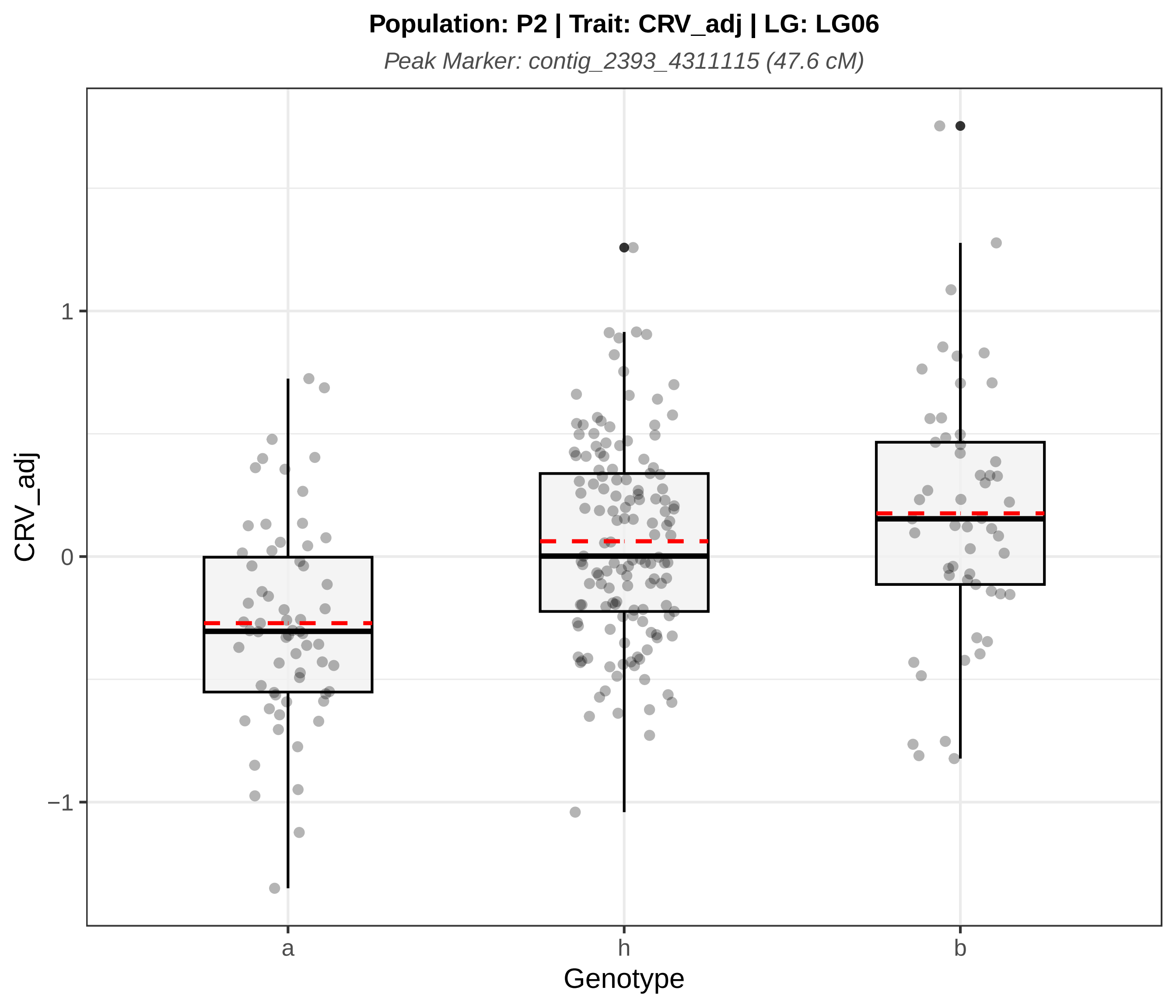

Population: P2 | Trait: CRV_adj | LG: LG06
Peak Marker: contig_2393_4311115 (47.6 cM)
CRV_adj
1
0
−1
a
h
b
Genotype

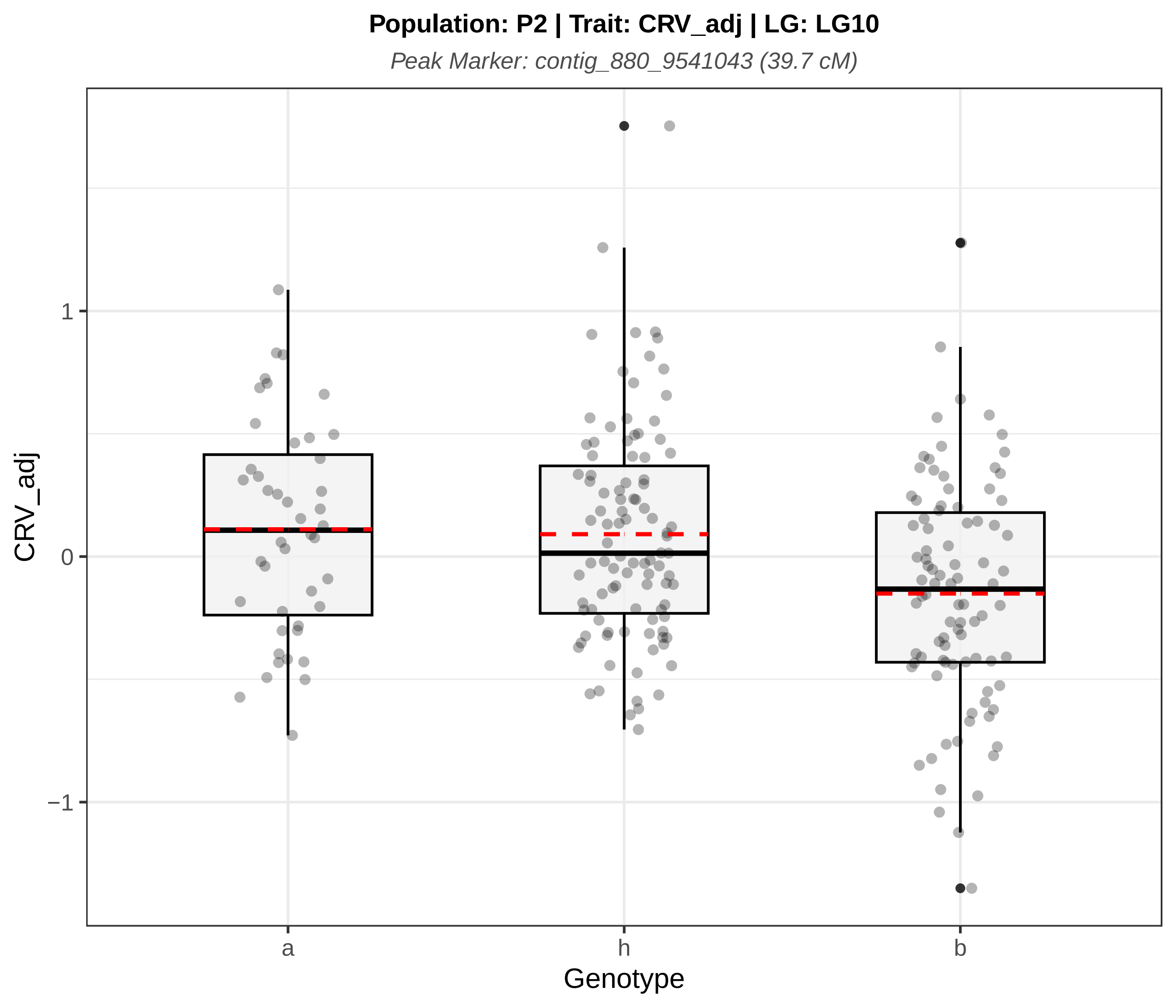

Population: P2 | Trait: CRV_adj | LG: LG10
Peak Marker: contig_880_9541043 (39.7 cM)
CRV_adj
1
0
−1
a
h
b
Genotype

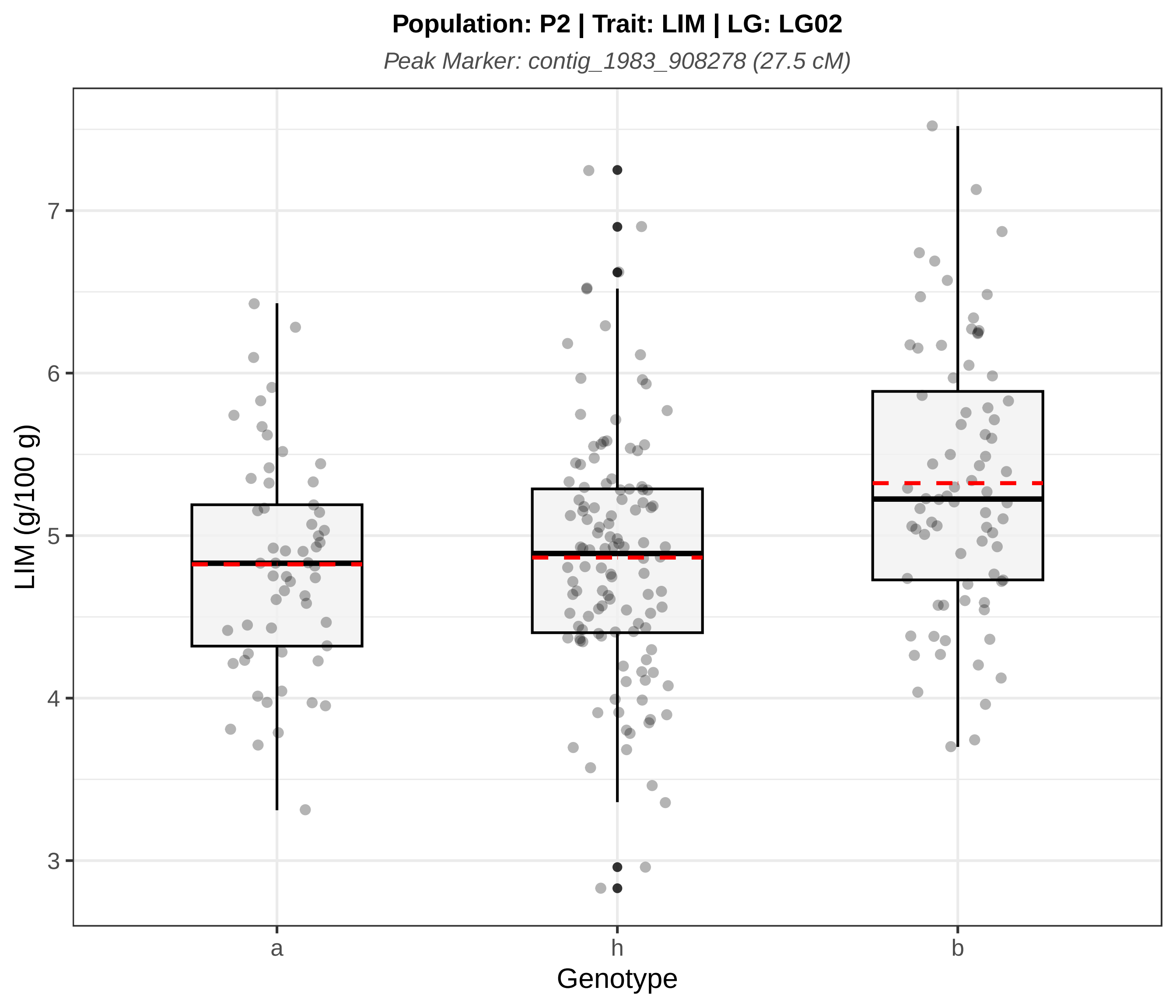
Population: P2 | Trait: LIM | LG: LG02
Peak Marker: contig_1983_908278 (27.5 cM)
LIM (g/100 g)
7
6
5
4
3
a
h
b
Genotype

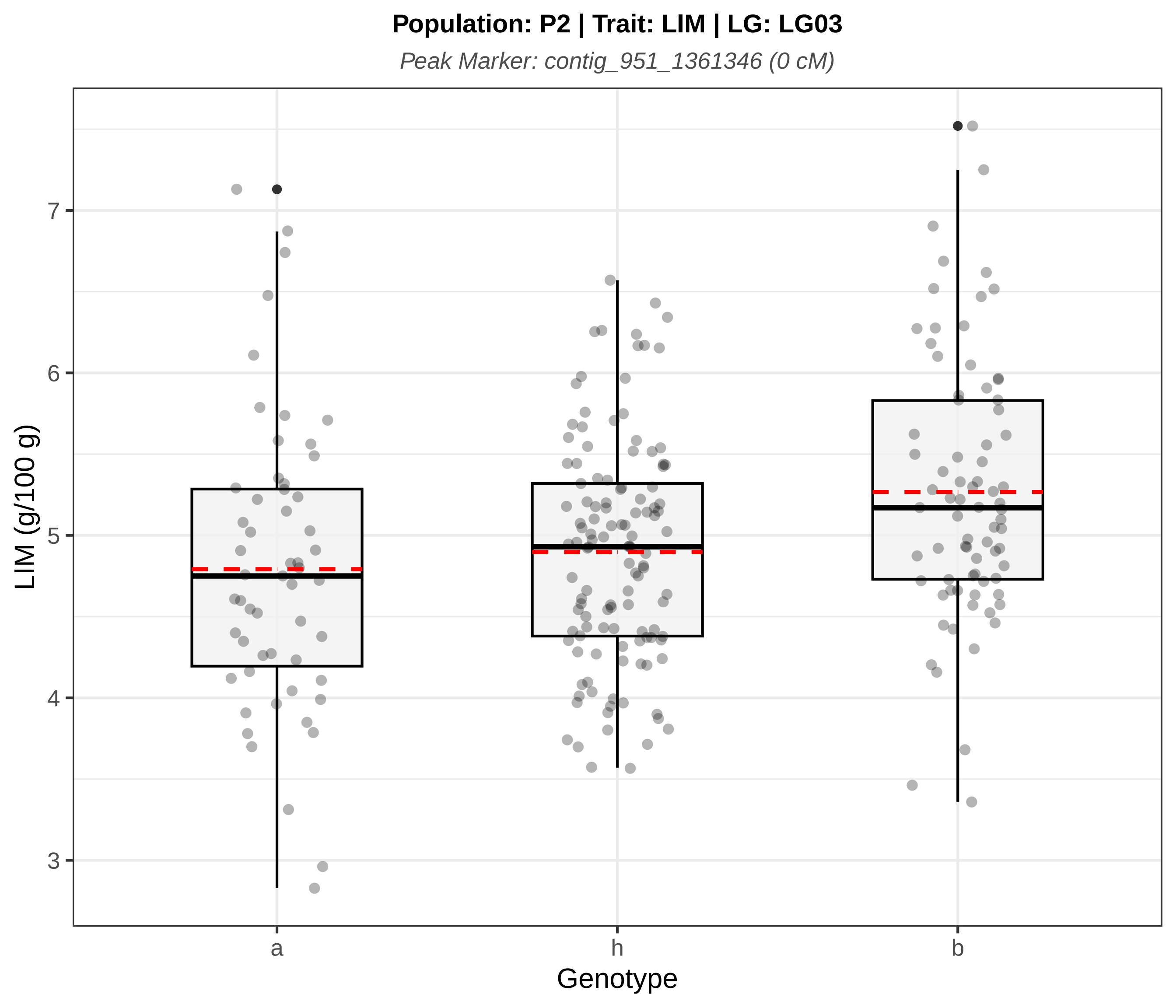
Population: P2 | Trait: LIM | LG: LG03
Peak Marker: contig_951_1361346 (0 cM)
LIM (g/100 g)
3
4
5
6
7
a
h
b
Genotype

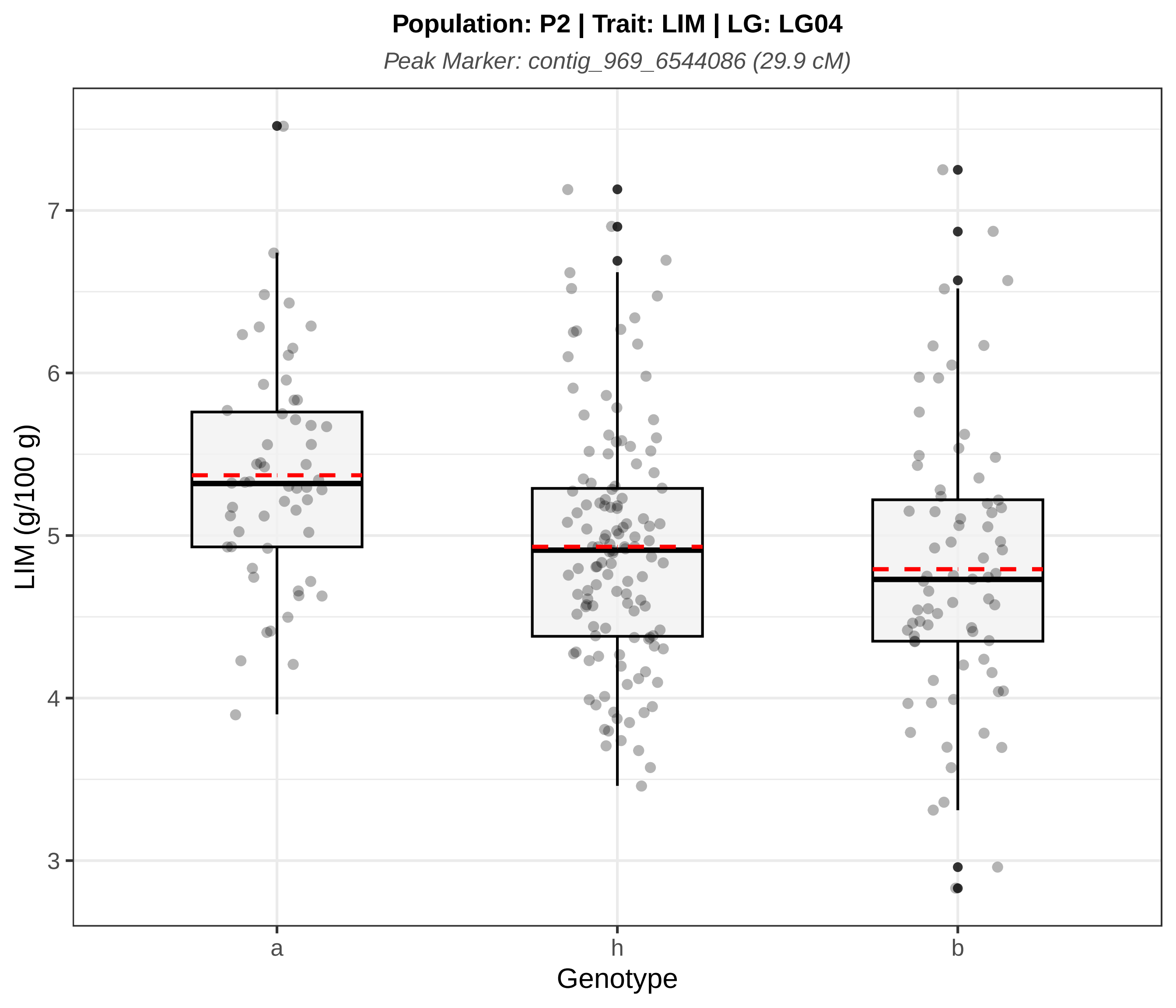
Population: P2 | Trait: LIM | LG: LG04
Peak Marker: contig_969_6544086 (29.9 cM)
LIM (g/100 g)
7
6
5
4
3
a
h
b
Genotype

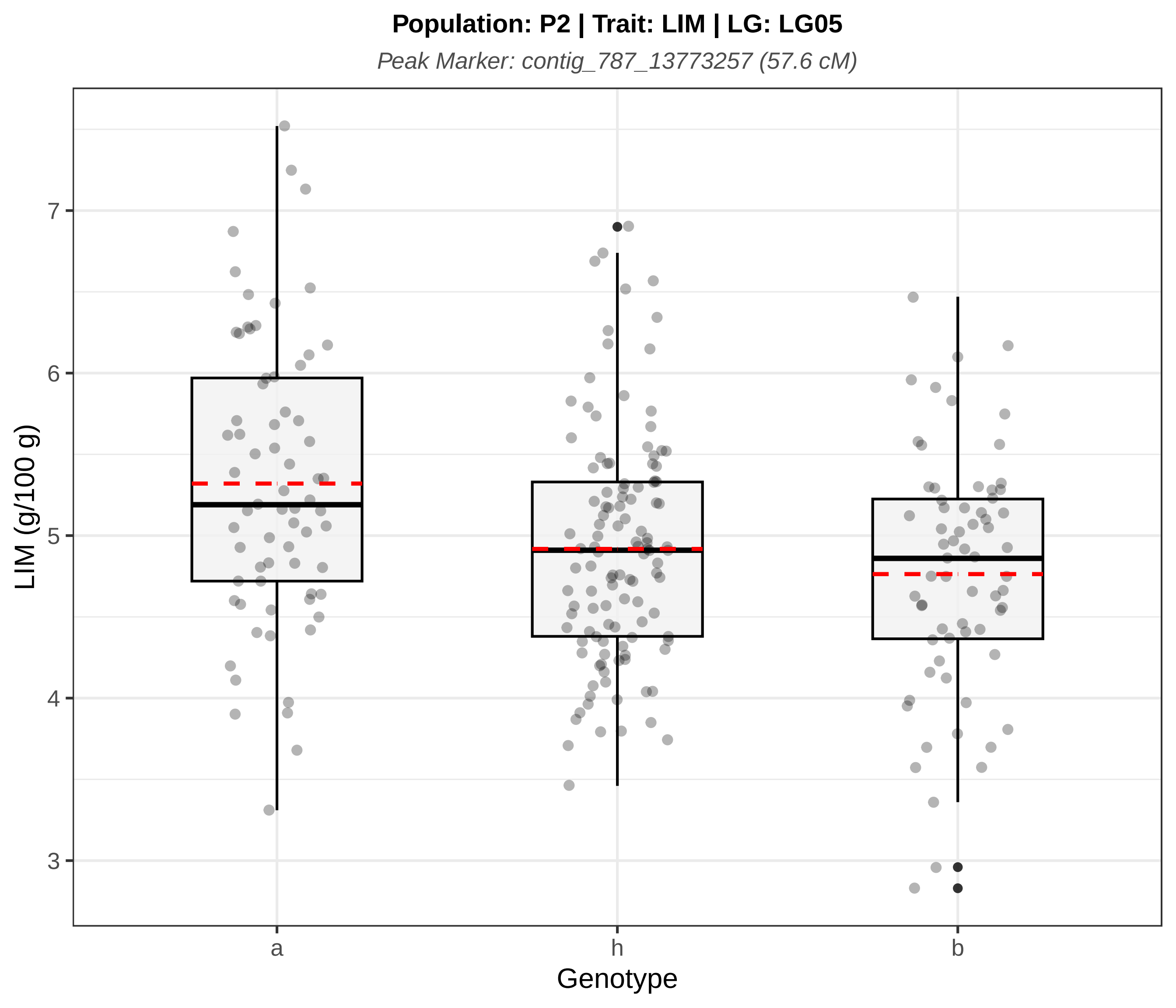
Population: P2 | Trait: LIM | LG: LG05
Peak Marker: contig_787_13773257 (57.6 cM)
LIM (g/100 g)
7
6
5
4
3
a
h
b
Genotype

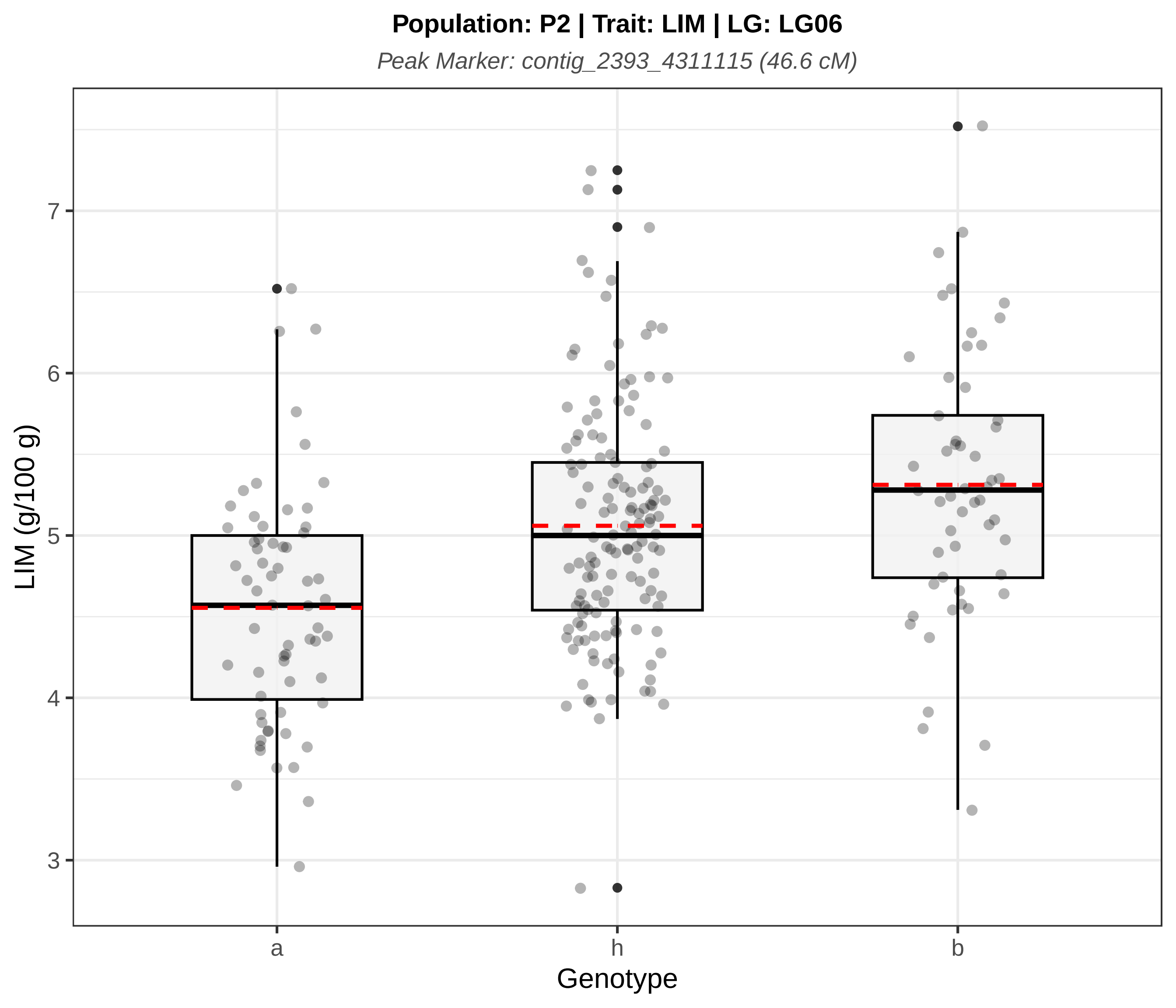

Population: P2 | Trait: LIM | LG: LG06
Peak Marker: contig_2393_4311115 (46.6 cM)
LIM (g/100 g)
7
6
5
4
3
a
h
b
Genotype

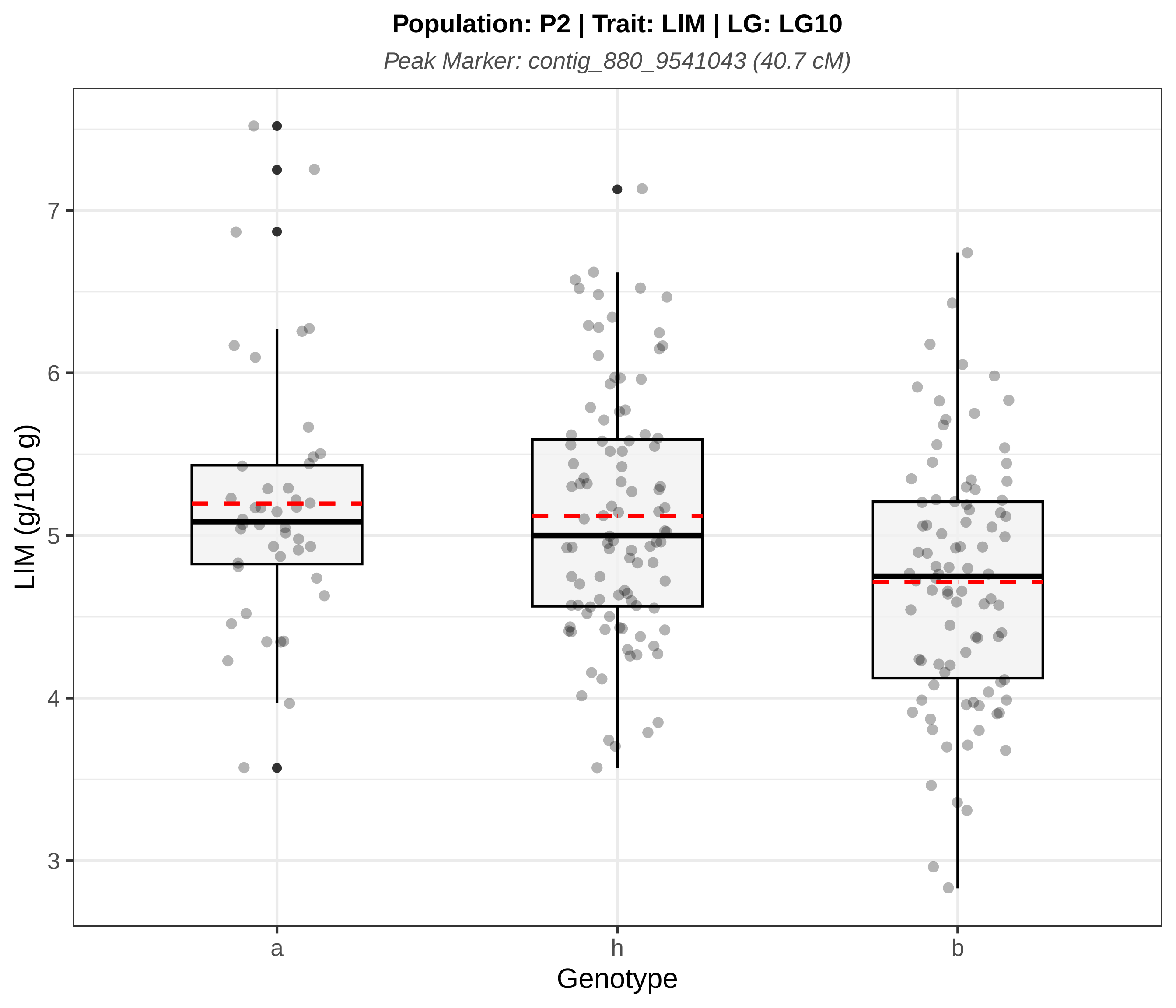
Population: P2 | Trait: LIM | LG: LG10
Peak Marker: contig_880_9541043 (40.7 cM)
LIM (g/100 g)
7
6
5
4
3
a
h
b
Genotype

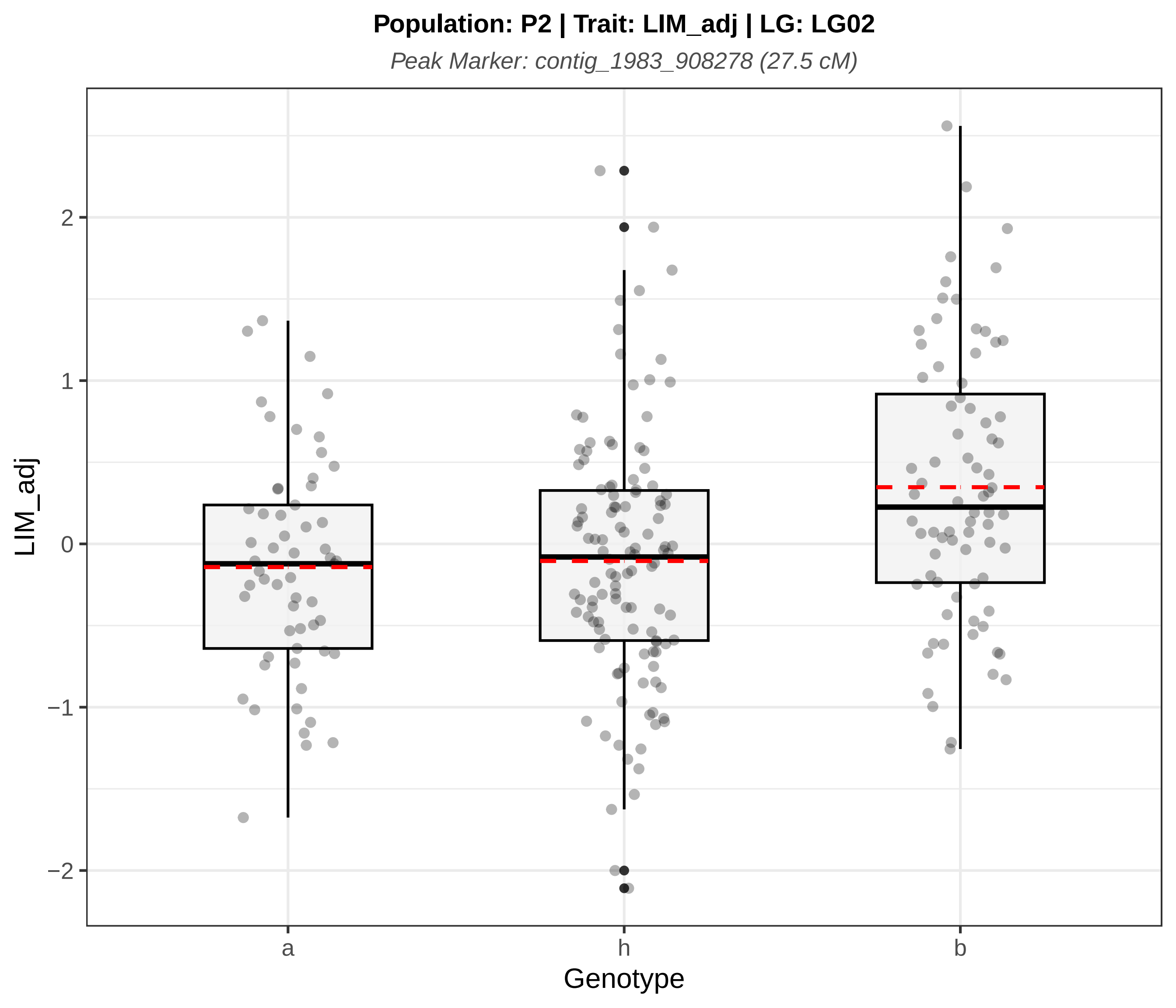

Population: P2 | Trait: LIM_adj | LG: LG02
Peak Marker: contig_1983_908278 (27.5 cM)
LIM_adj
2
1
0
−1
−2
a
h
b
Genotype

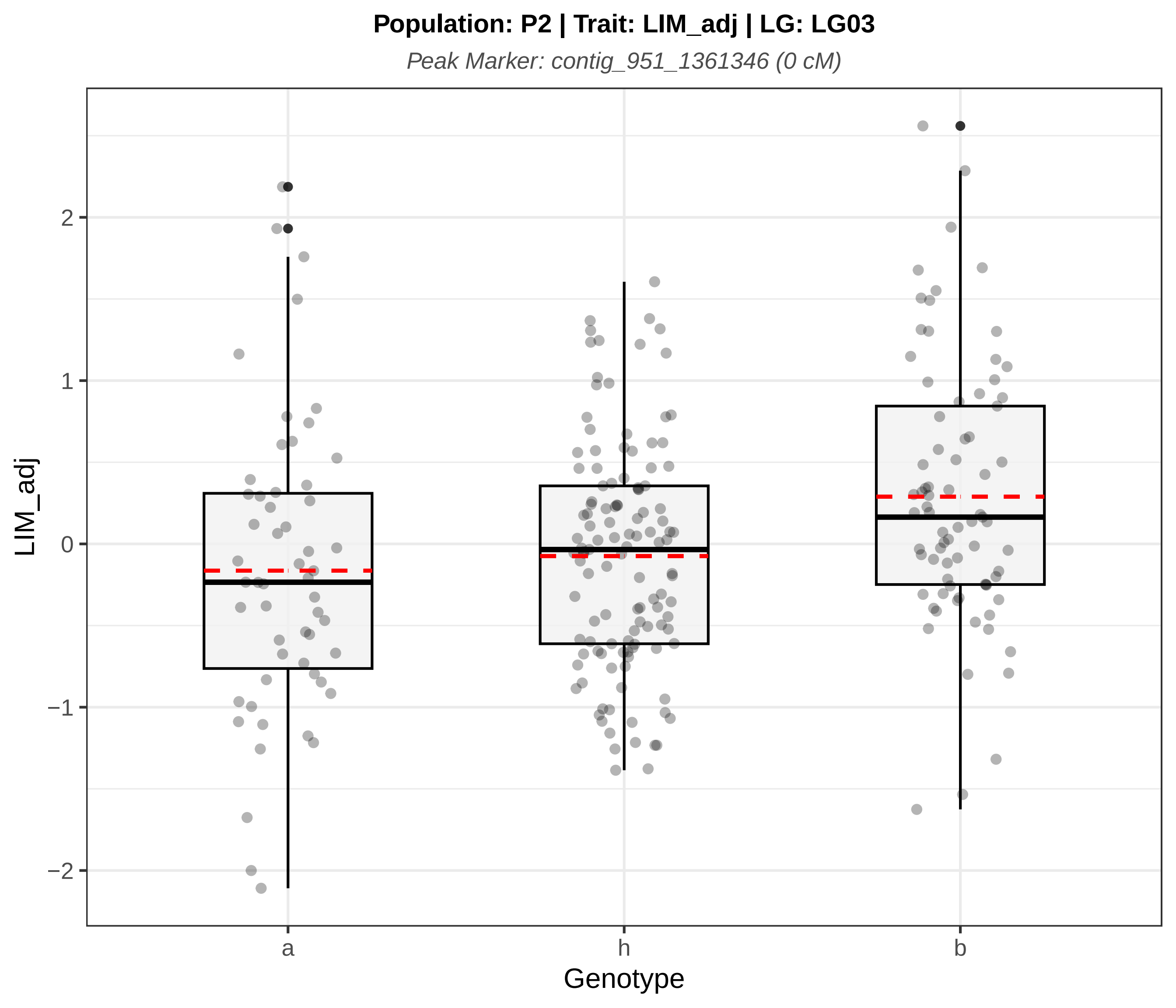
Population: P2 | Trait: LIM_adj | LG: LG03
Peak Marker: contig_951_1361346 (0 cM)
LIM_adj
Genotype
a
h
b
2
1
0
−1
−2

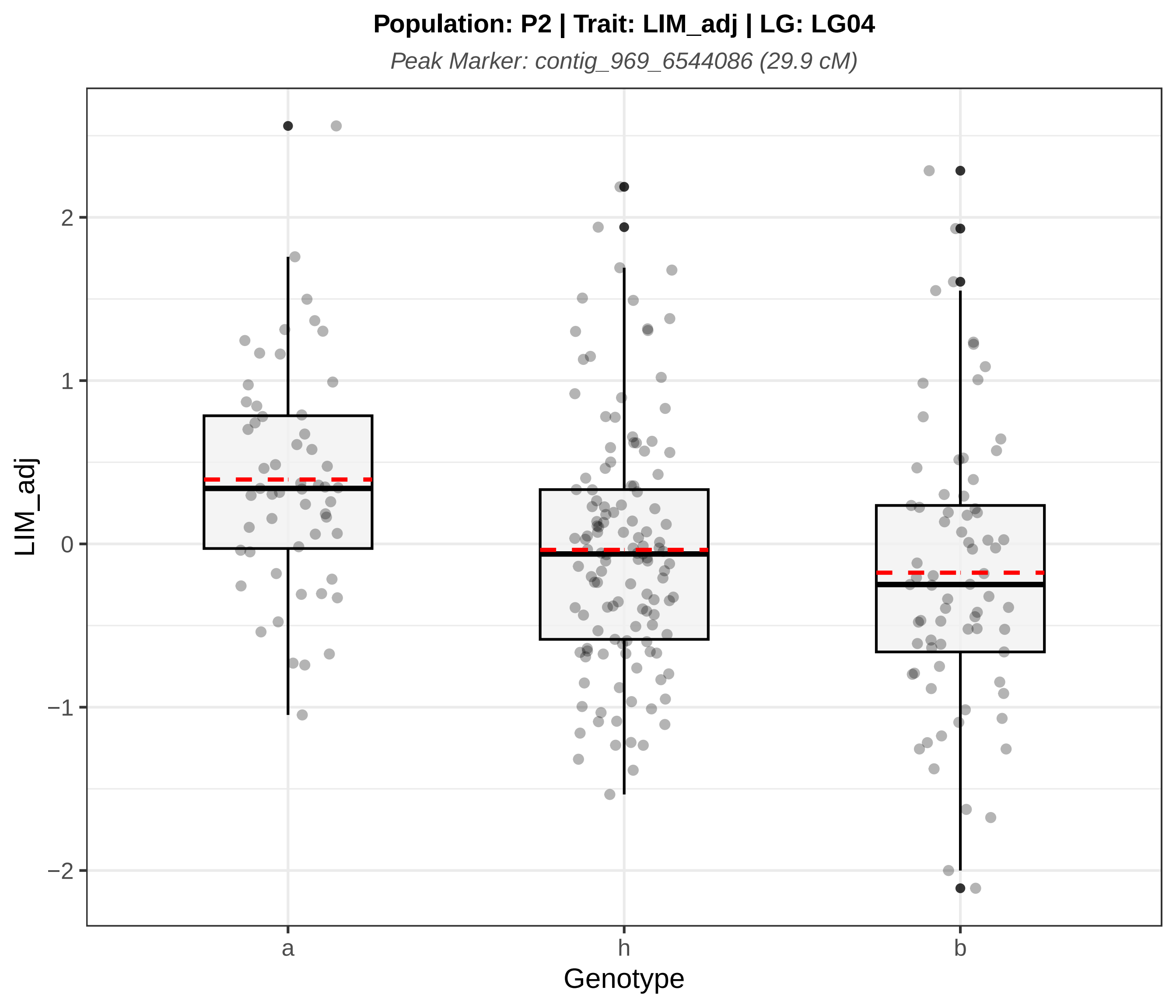

Population: P2 | Trait: LIM_adj | LG: LG04
Peak Marker: contig_969_6544086 (29.9 cM)
LIM_adj
2
1
0
−1
−2
a
h
b
Genotype

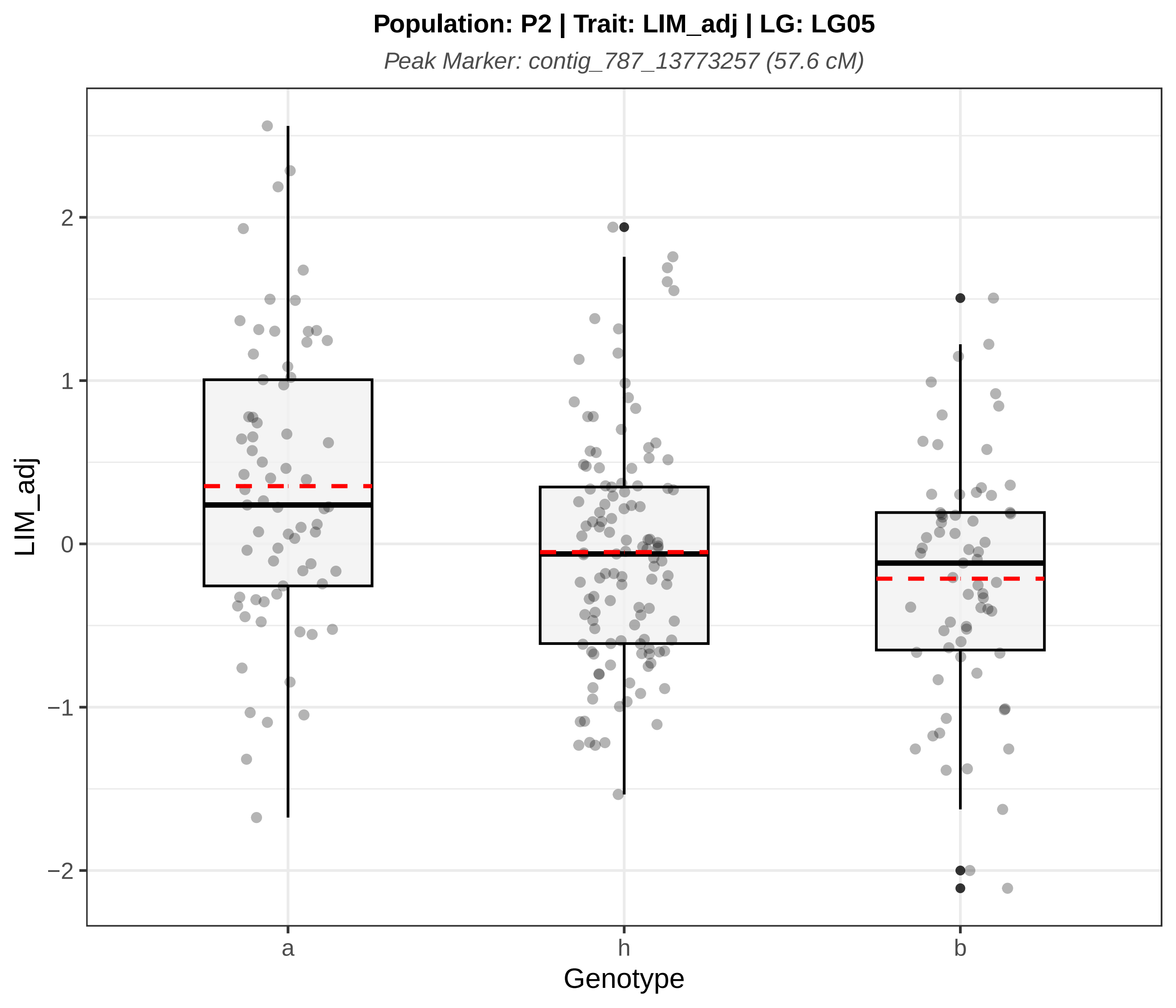

Population: P2 | Trait: LIM_adj | LG: LG05
Peak Marker: contig_787_13773257 (57.6 cM)
LIM_adj
2
1
0
−1
−2
a
h
b
Genotype

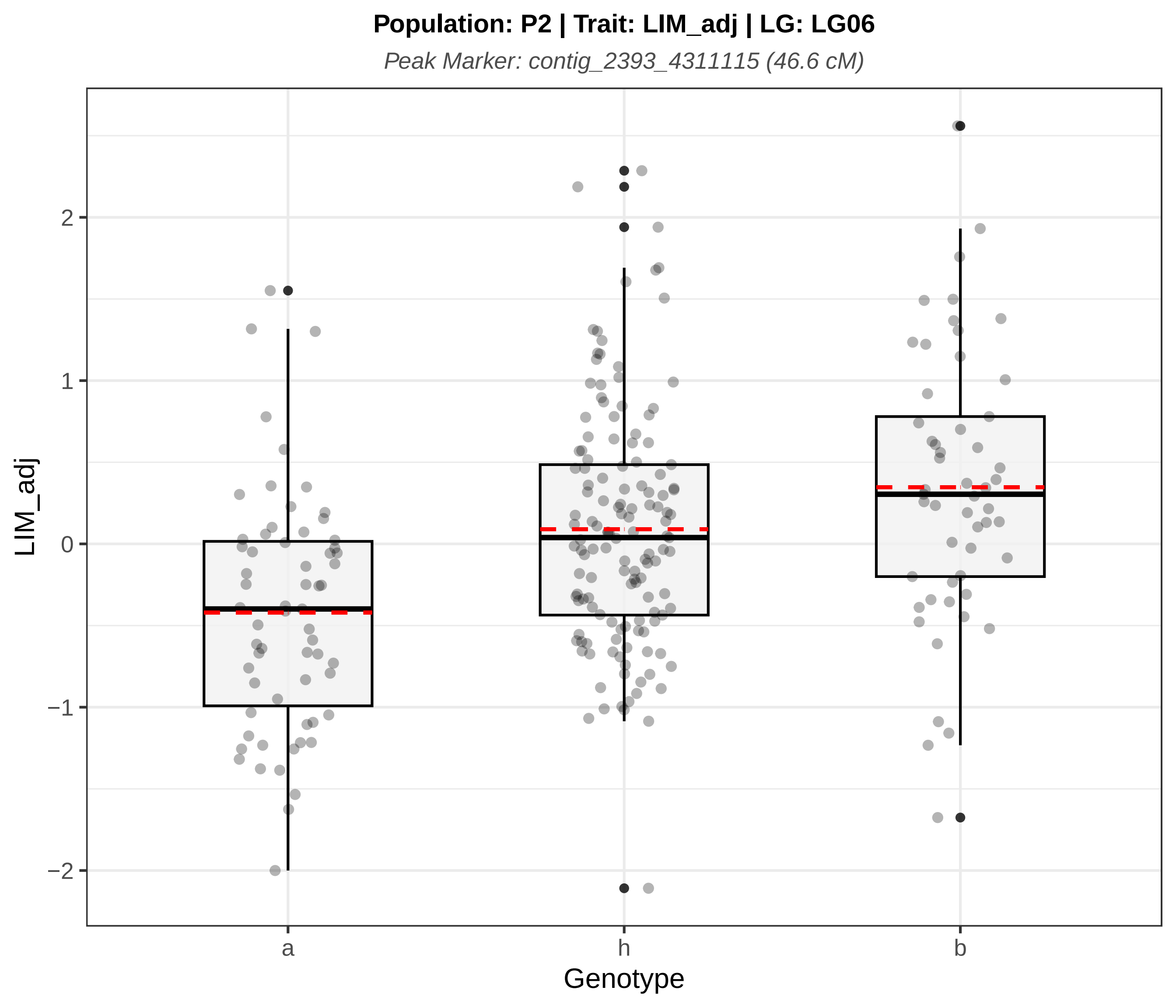
Population: P2 | Trait: LIM_adj | LG: LG06
Peak Marker: contig_2393_4311115 (46.6 cM)
LIM_adj
2
1
0
−1
−2
a
h
b
Genotype

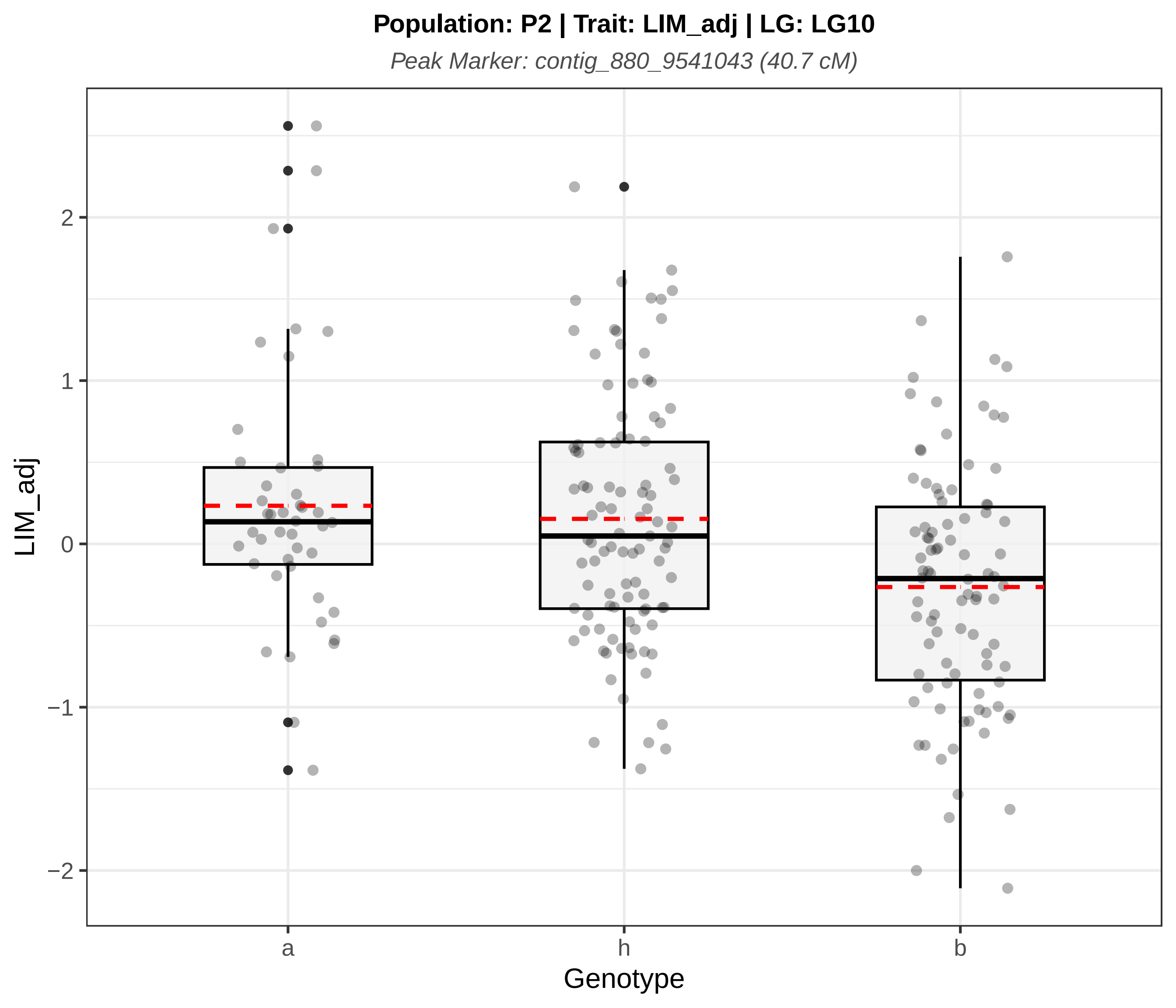
Population: P2 | Trait: LIM_adj | LG: LG10
Peak Marker: contig_880_9541043 (40.7 cM)
LIM_adj
2
1
0
−1
−2
a
h
b
Genotype

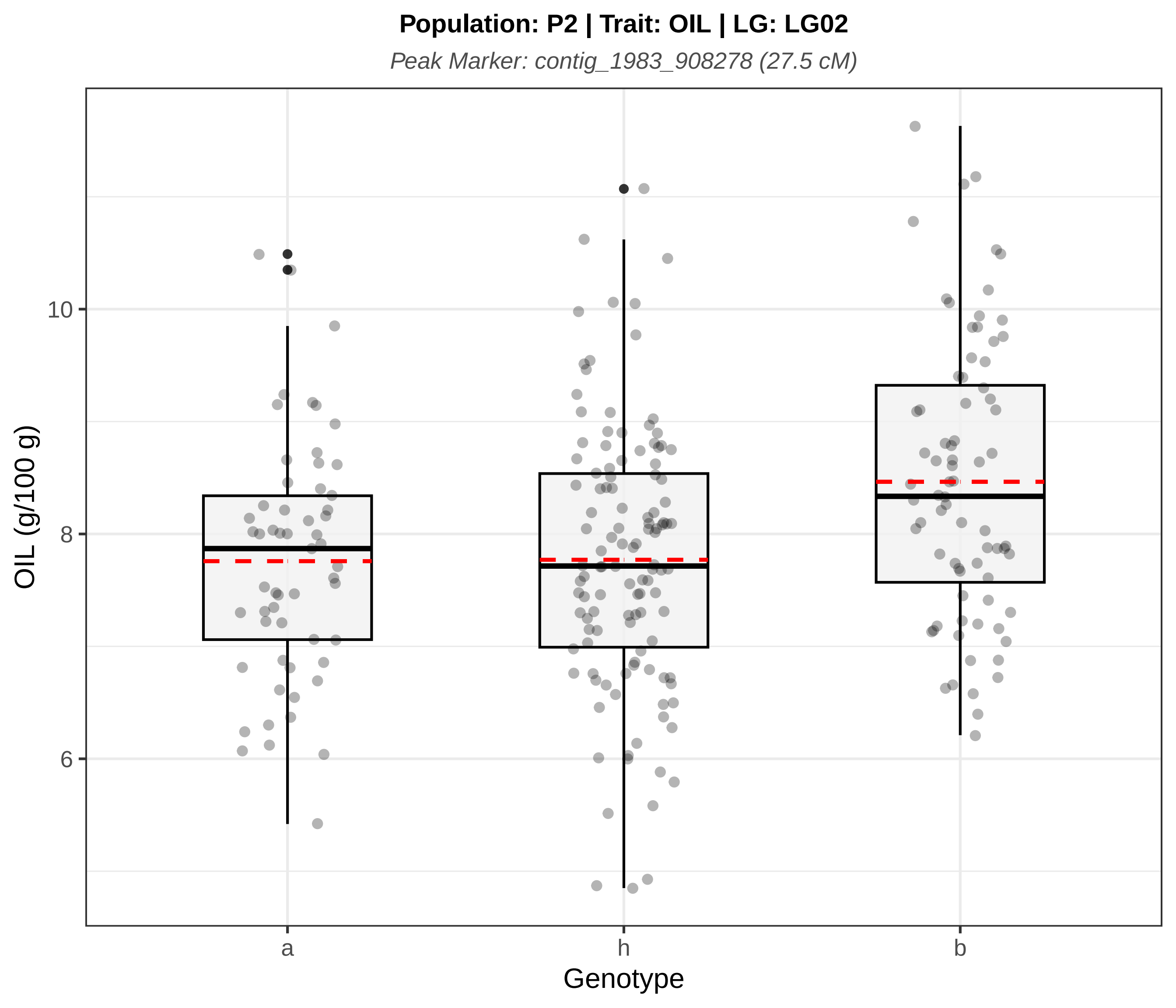
Population: P2 | Trait: OIL | LG: LG02
Peak Marker: contig_1983_908278 (27.5 cM)
OIL (g/100 g)
10
8
6
a
h
b
Genotype

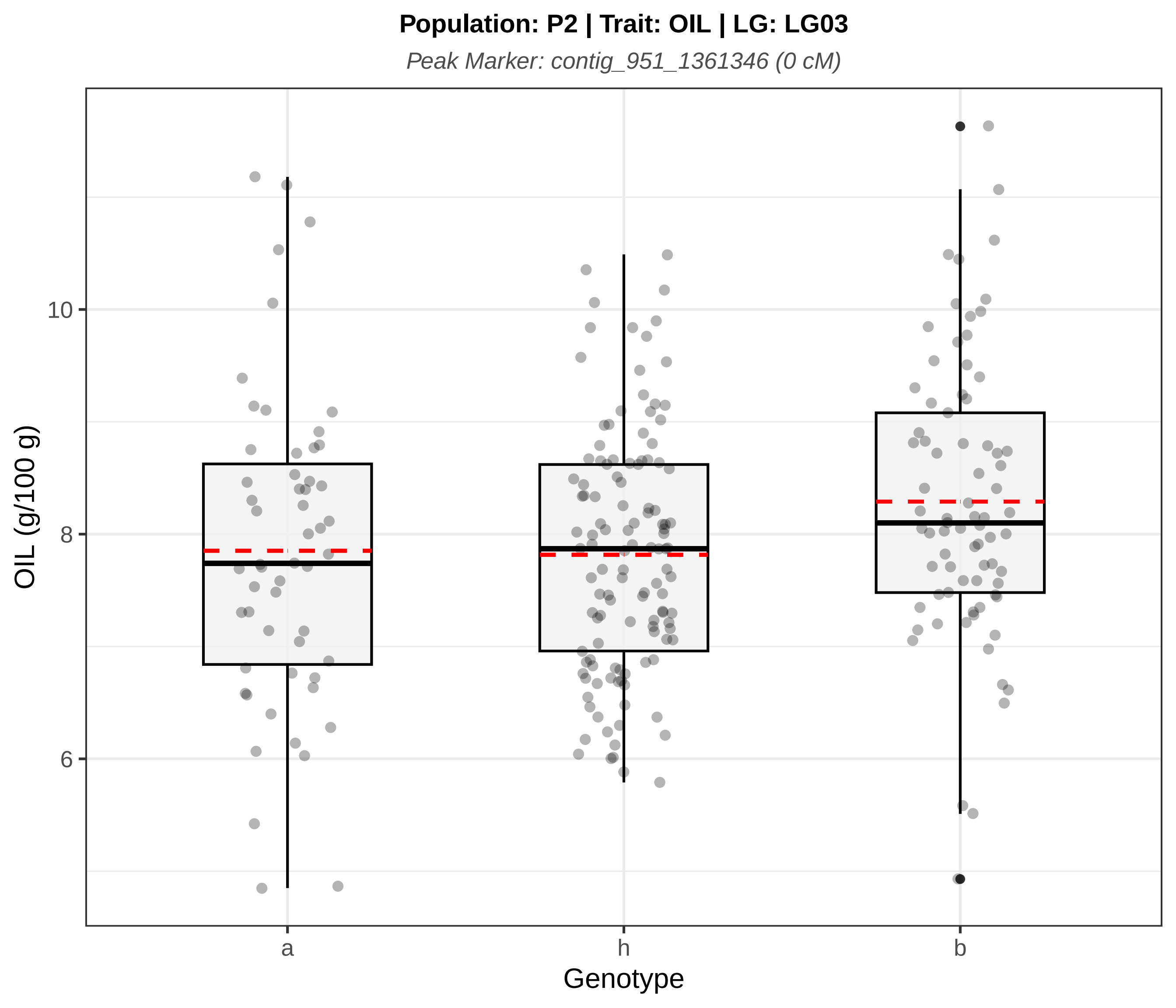
Population: P2 | Trait: OIL | LG: LG03
Peak Marker: contig_951_1361346 (0 cM)
OIL (g/100 g)
10
8
6
a
h
b
Genotype

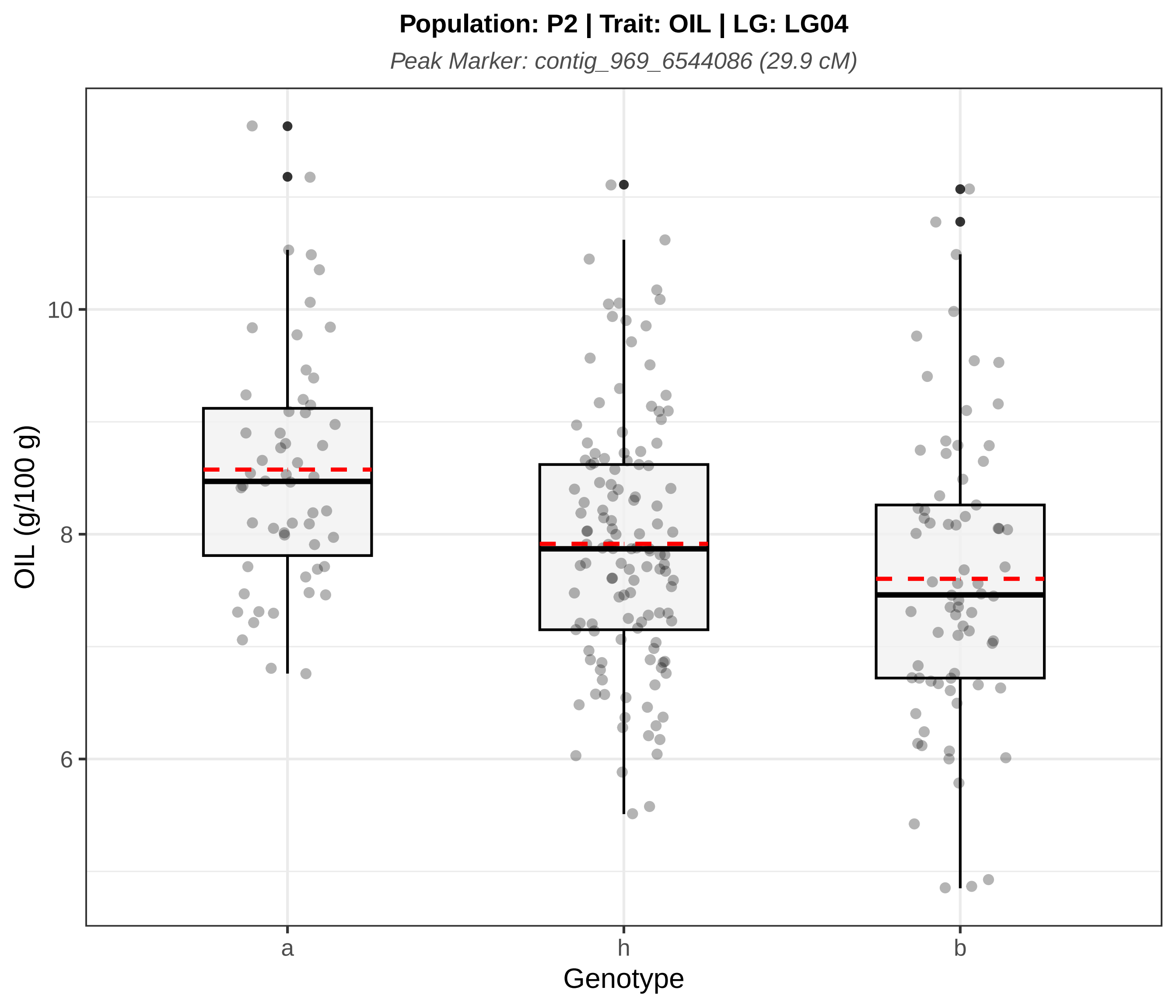
Population: P2 | Trait: OIL | LG: LG04
Peak Marker: contig_969_6544086 (29.9 cM)
OIL (g/100 g)
10
8
6
a
h
b
Genotype

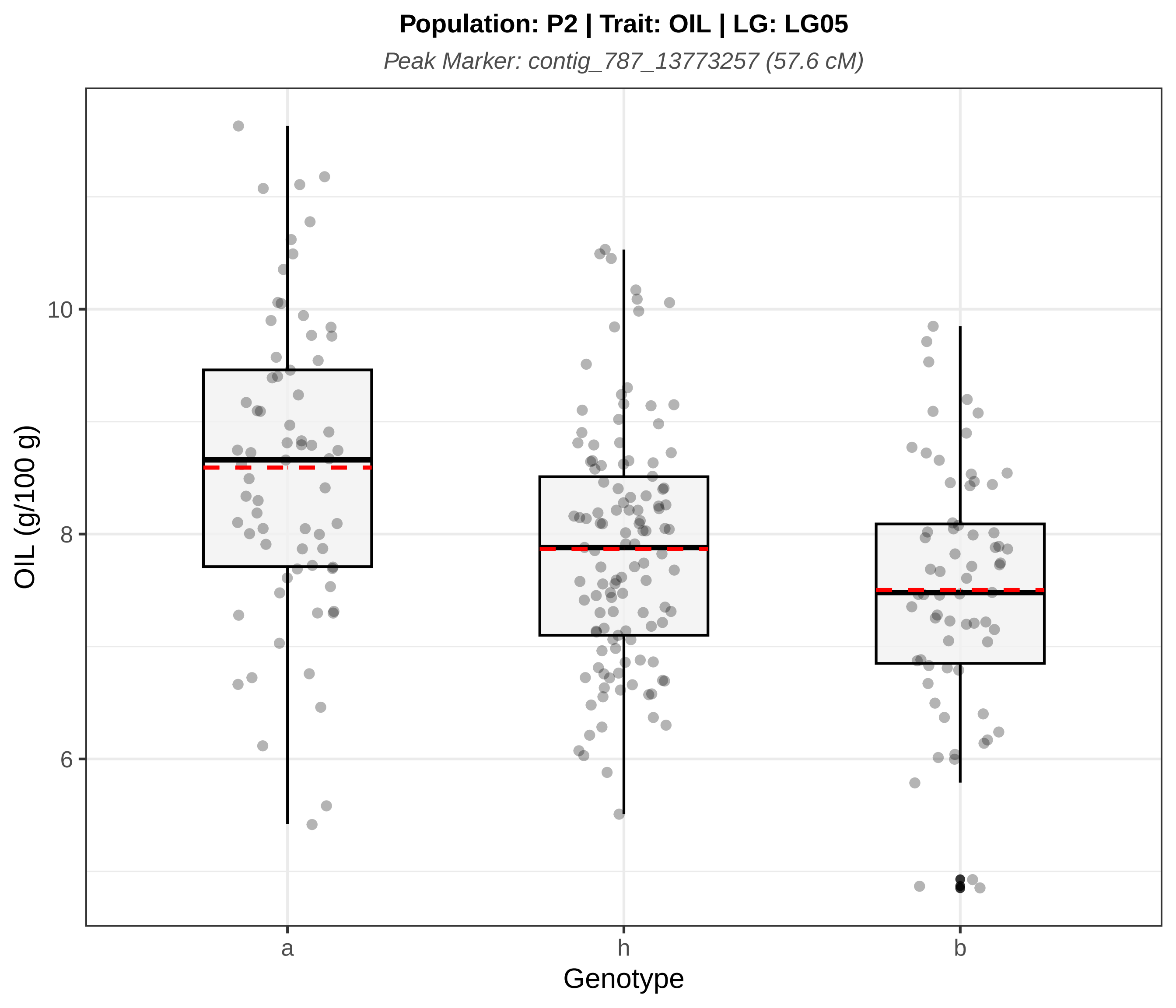
Population: P2 | Trait: OIL | LG: LG05
Peak Marker: contig_787_13773257 (57.6 cM)
OIL (g/100 g)
10
8
6
a
h
b
Genotype

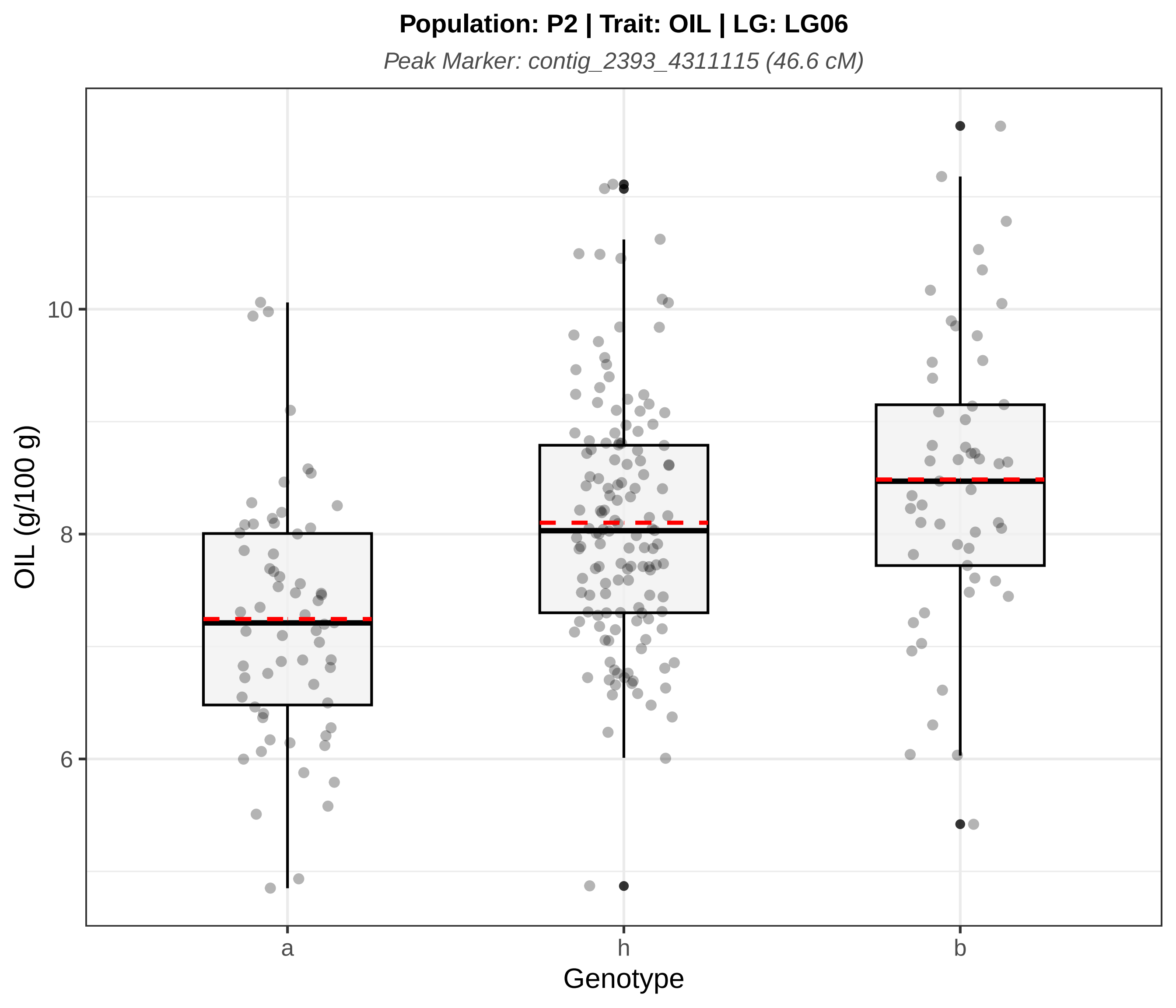

Population: P2 | Trait: OIL | LG: LG06
Peak Marker: contig_2393_4311115 (46.6 cM)
OIL (g/100 g)
10
8
6
a
h
b
Genotype

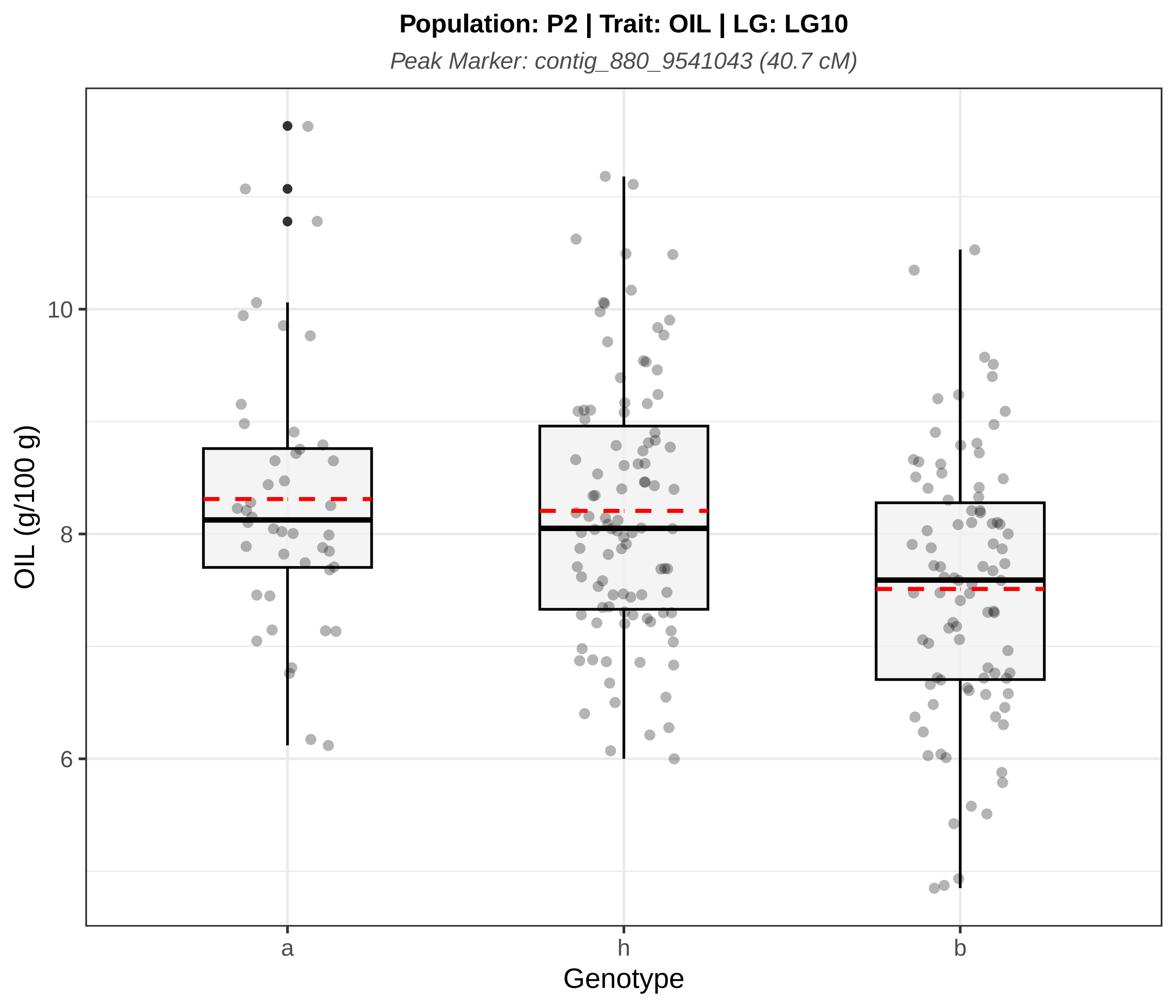

Population: P2 | Trait: OIL | LG: LG10
Peak Marker: contig_880_9541043 (40.7 cM)
OIL (g/100 g)
10
8
6
a
h
b
Genotype

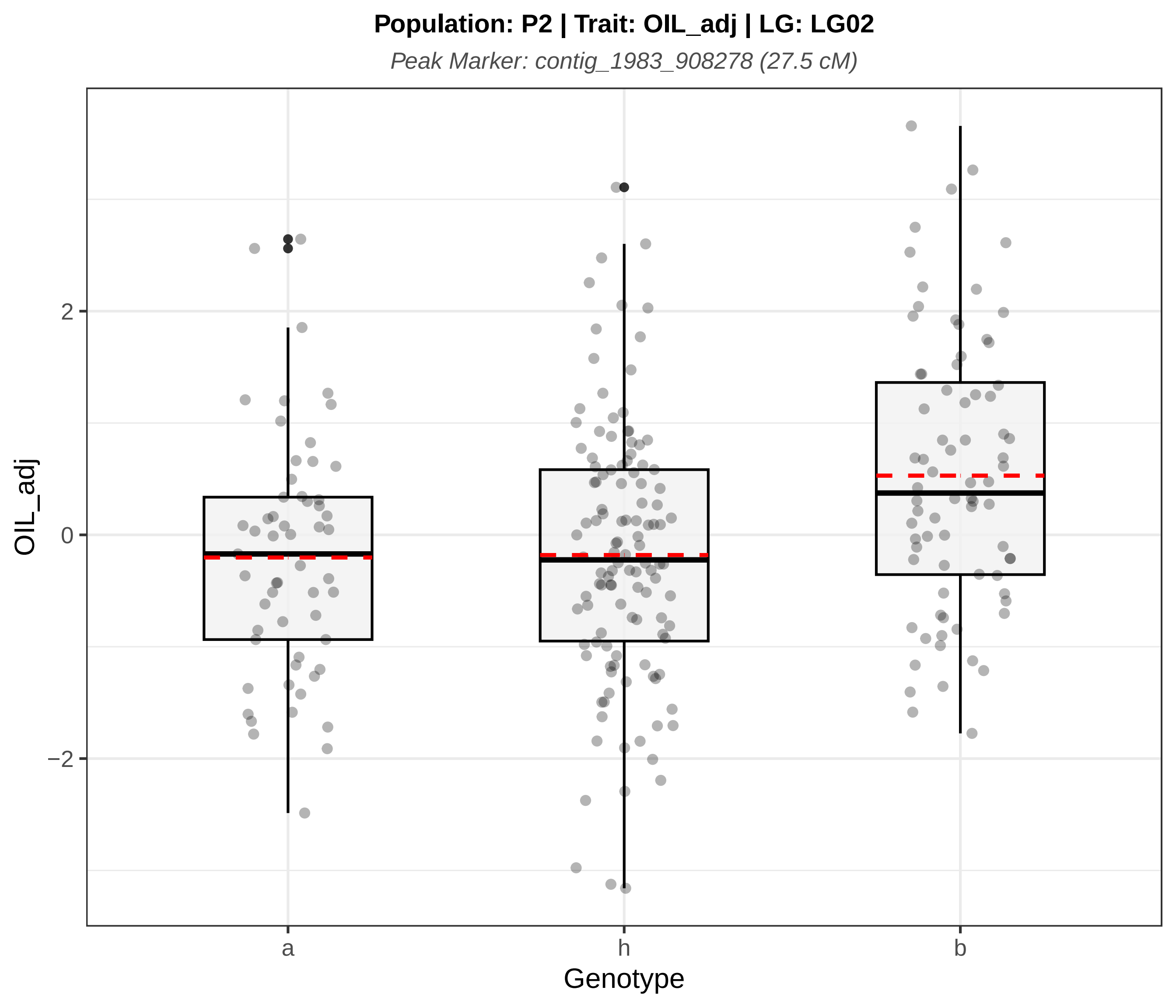
Population: P2 | Trait: OIL_adj | LG: LG02
Peak Marker: contig_1983_908278 (27.5 cM)
OIL_adj
2
0
−2
a
h
b
Genotype

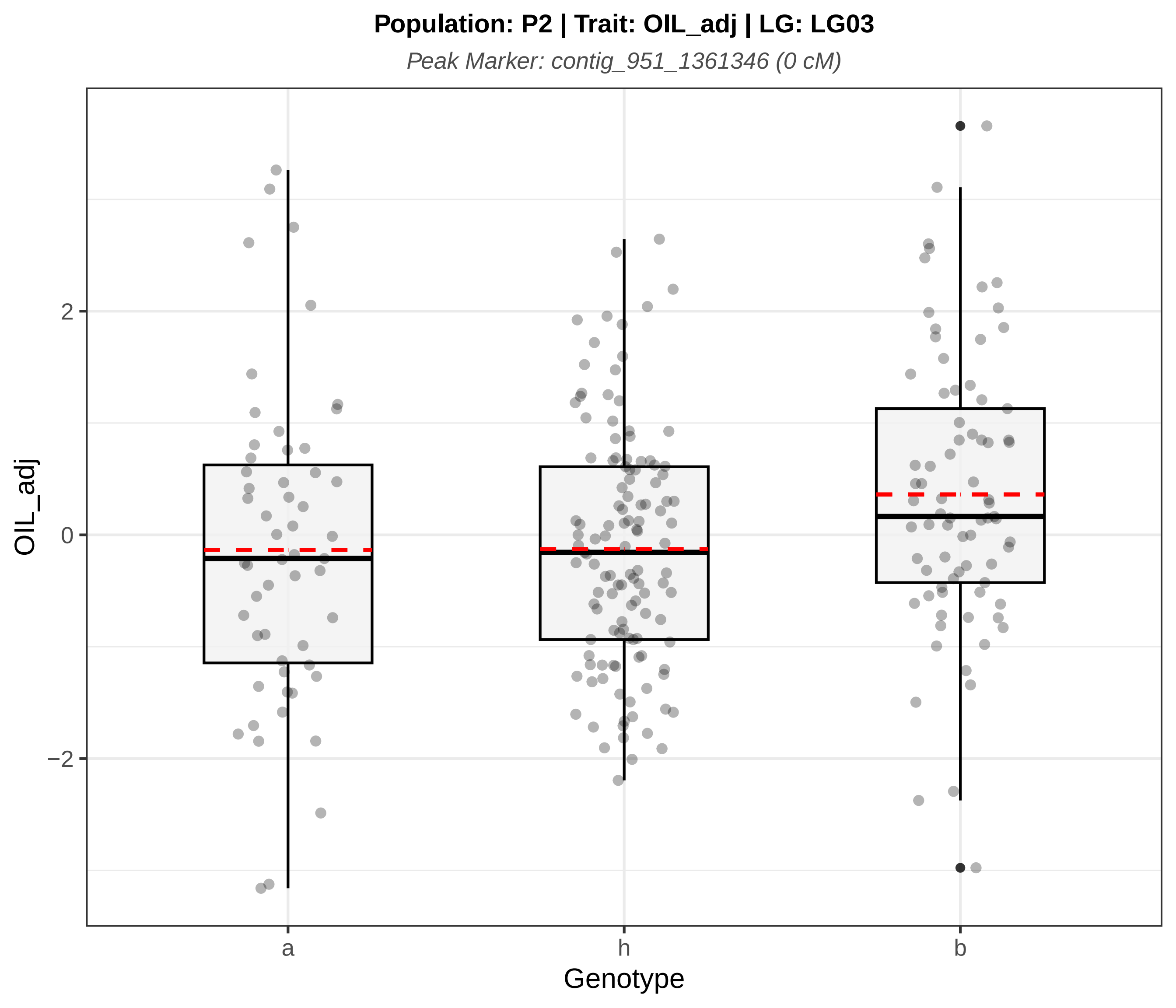

Population: P2 | Trait: OIL_adj | LG: LG03
Peak Marker: contig_951_1361346 (0 cM)
OIL_adj
2
0
−2
a
h
b
Genotype

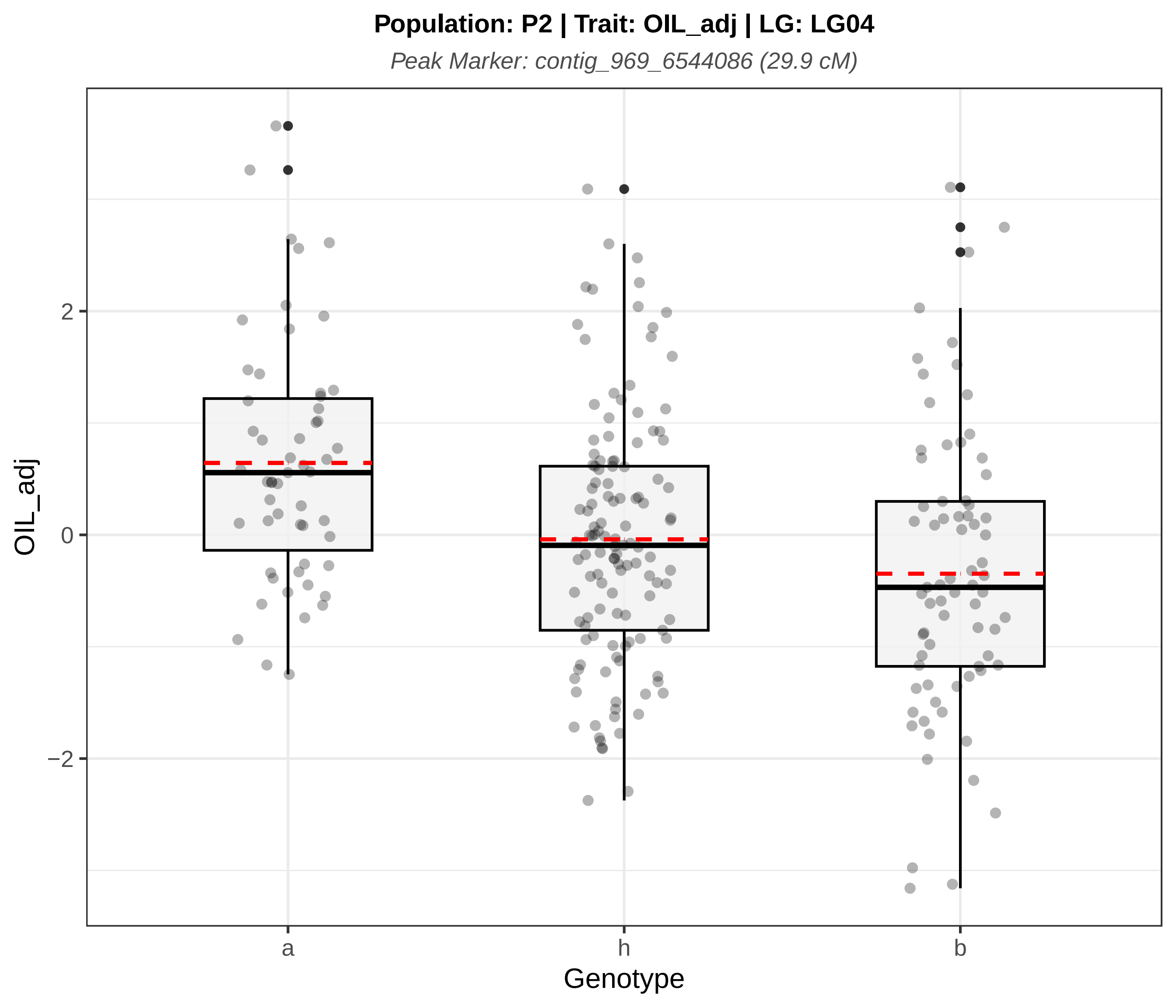
Population: P2 | Trait: OIL_adj | LG: LG04
Peak Marker: contig_969_6544086 (29.9 cM)
OIL_adj
2
0
−2
a
h
b
Genotype

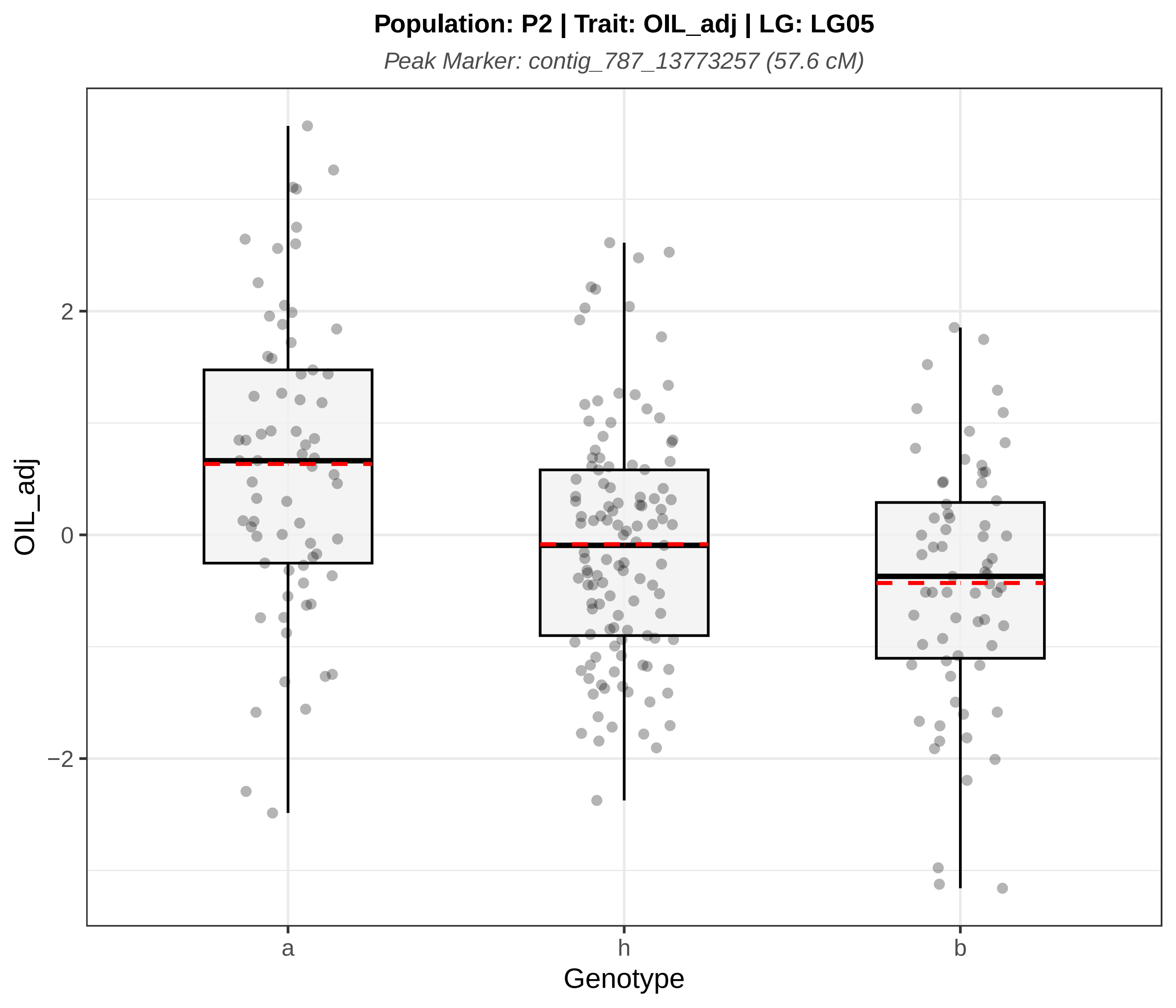
Population: P2 | Trait: OIL_adj | LG: LG05
Peak Marker: contig_787_13773257 (57.6 cM)
OIL_adj
2
0
−2
a
h
b
Genotype

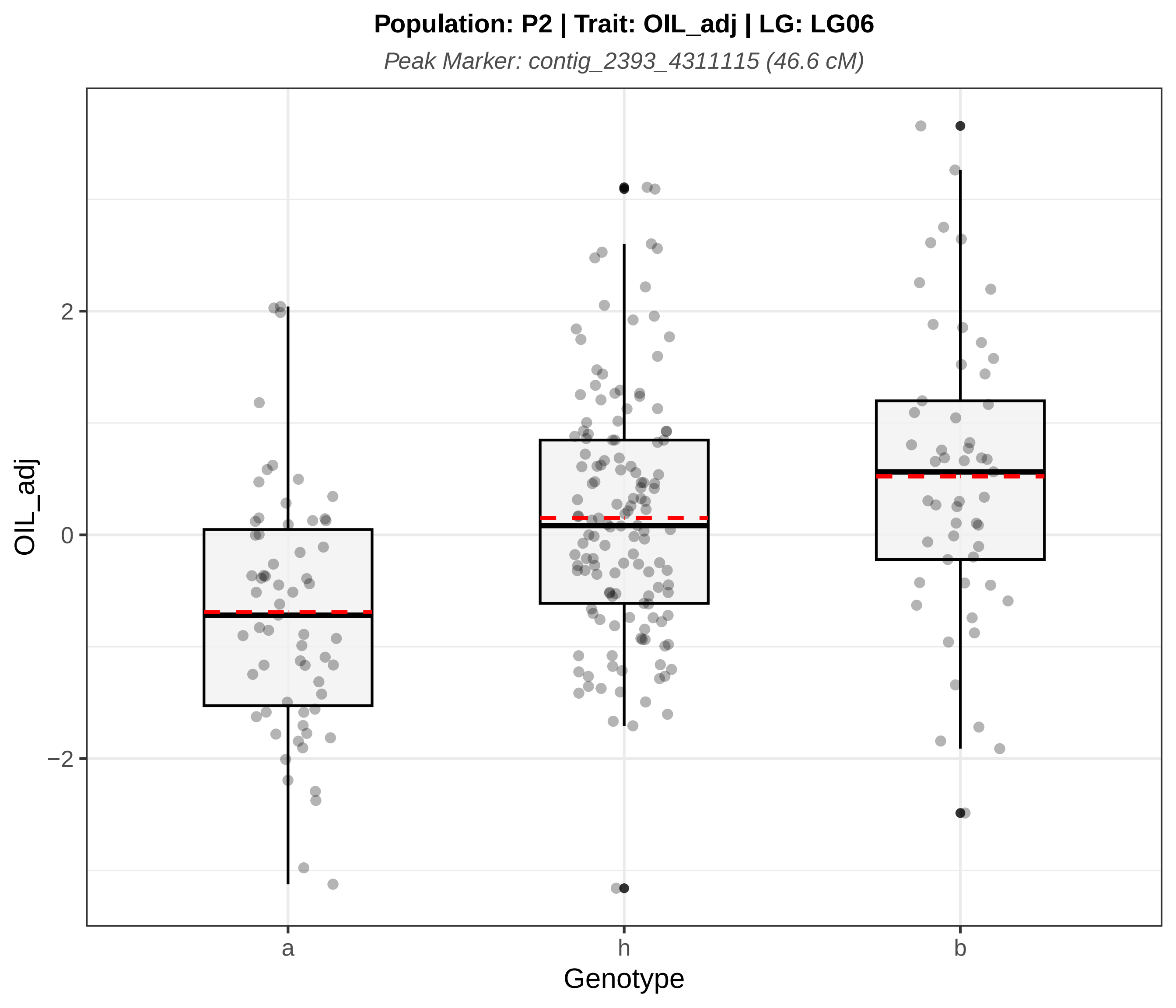
Population: P2 | Trait: OIL_adj | LG: LG06
Peak Marker: contig_2393_4311115 (46.6 cM)
OIL_adj
2
0
−2
a
h
b
Genotype

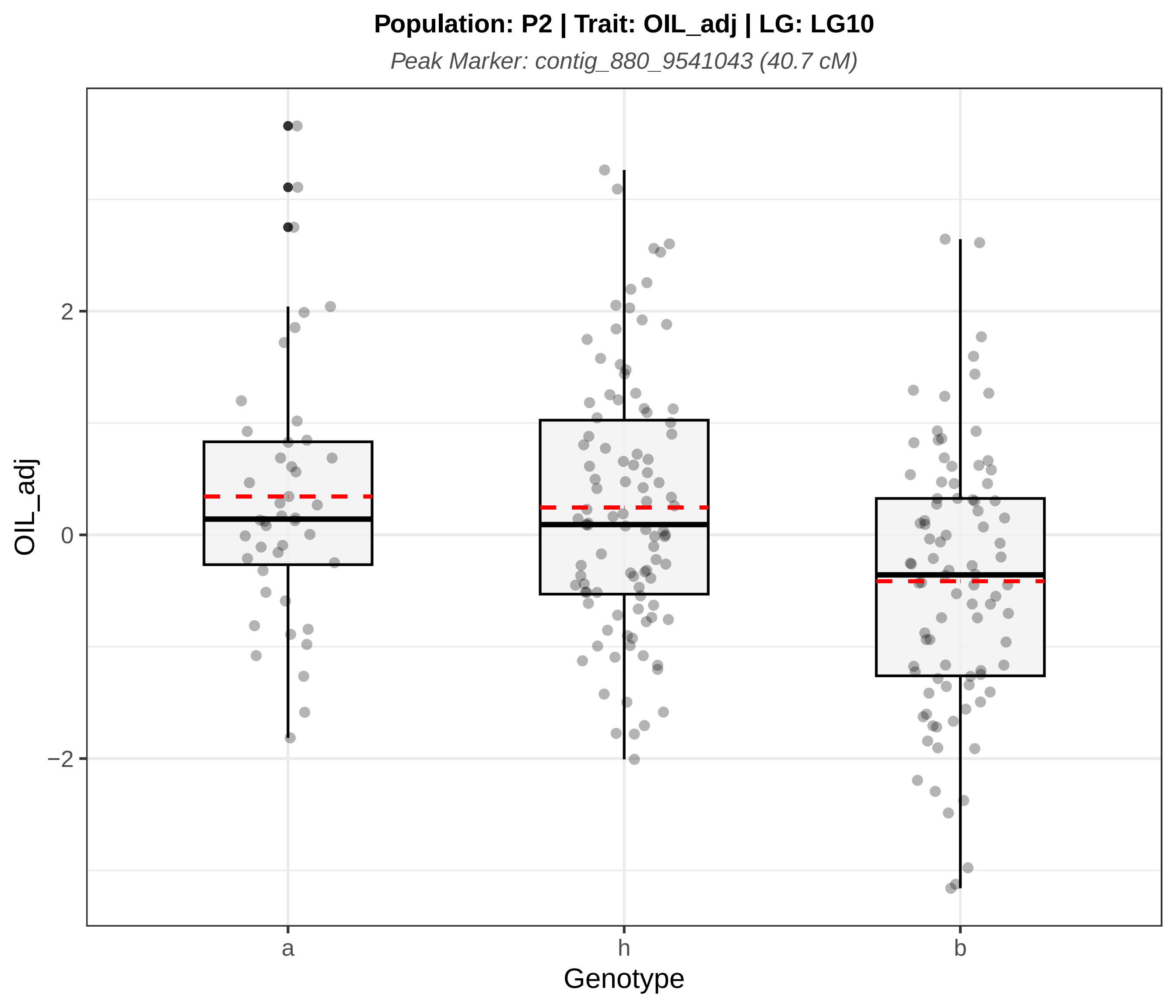
Population: P2 | Trait: OIL_adj | LG: LG10
Peak Marker: contig_880_9541043 (40.7 cM)
OIL_adj
2
0
−2
a
h
b
Genotype

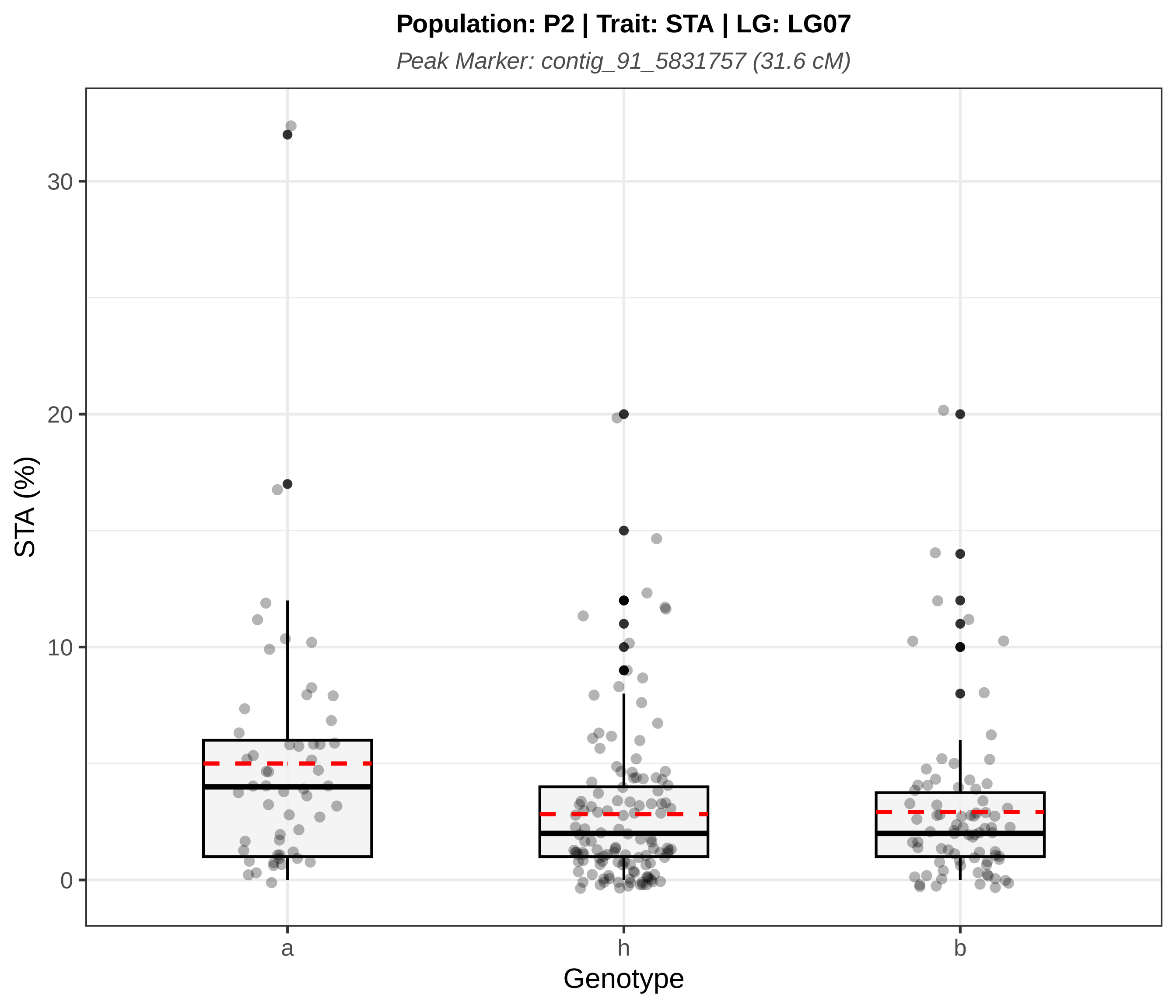
Population: P2 | Trait: STA | LG: LG07
Peak Marker: contig_91_5831757 (31.6 cM)
STA (%)
0
10
20
30
a
h
b
Genotype

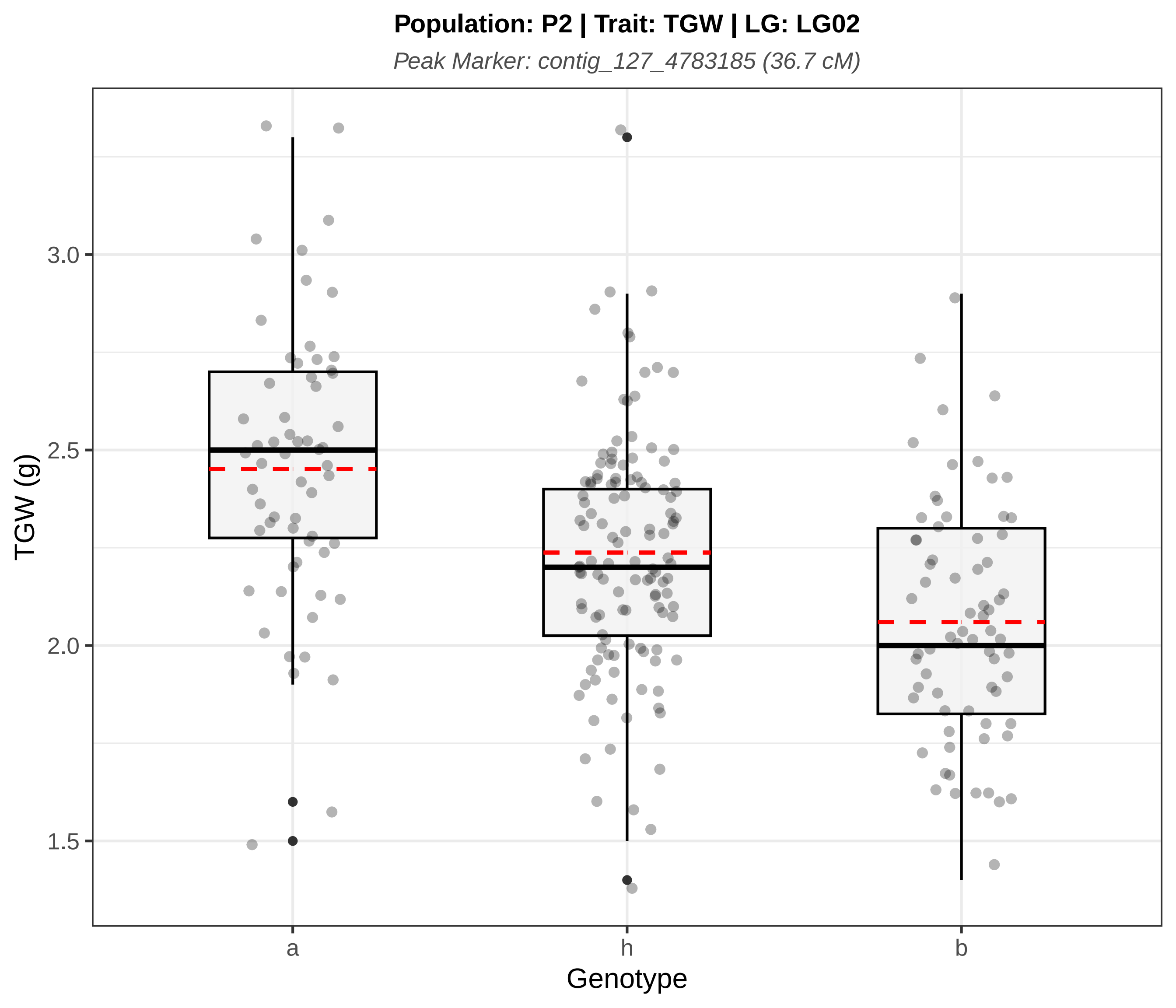
Population: P2 | Trait: TGW | LG: LG02
Peak Marker: contig_127_4783185 (36.7 cM)
TGW (g)
3.0
2.5
2.0
1.5
a
h
b
Genotype

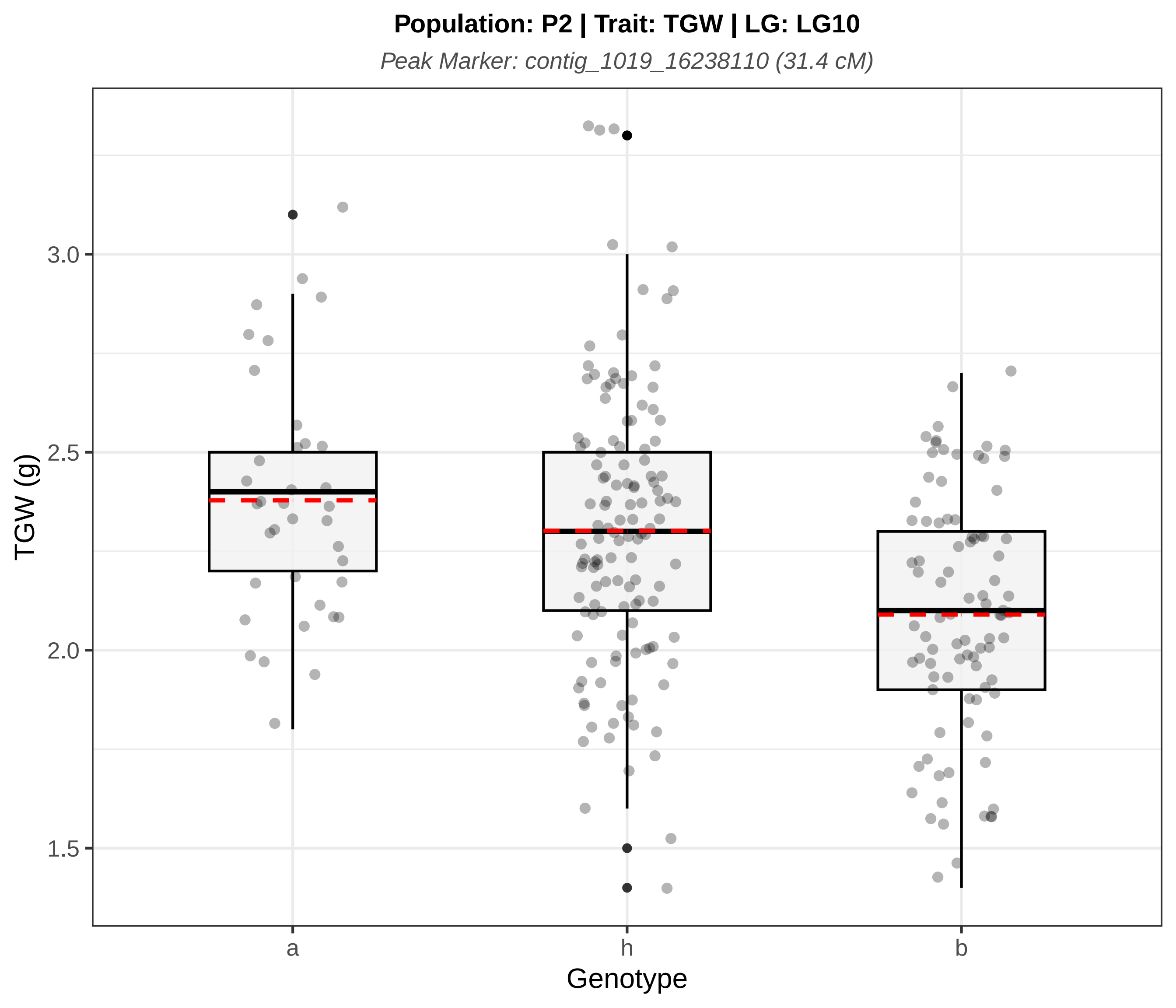

Population: P2 | Trait: TGW | LG: LG10
Peak Marker: contig_1019_16238110 (31.4 cM)
TGW (g)
3.0
2.5
2.0
1.5
a
h
b
Genotype

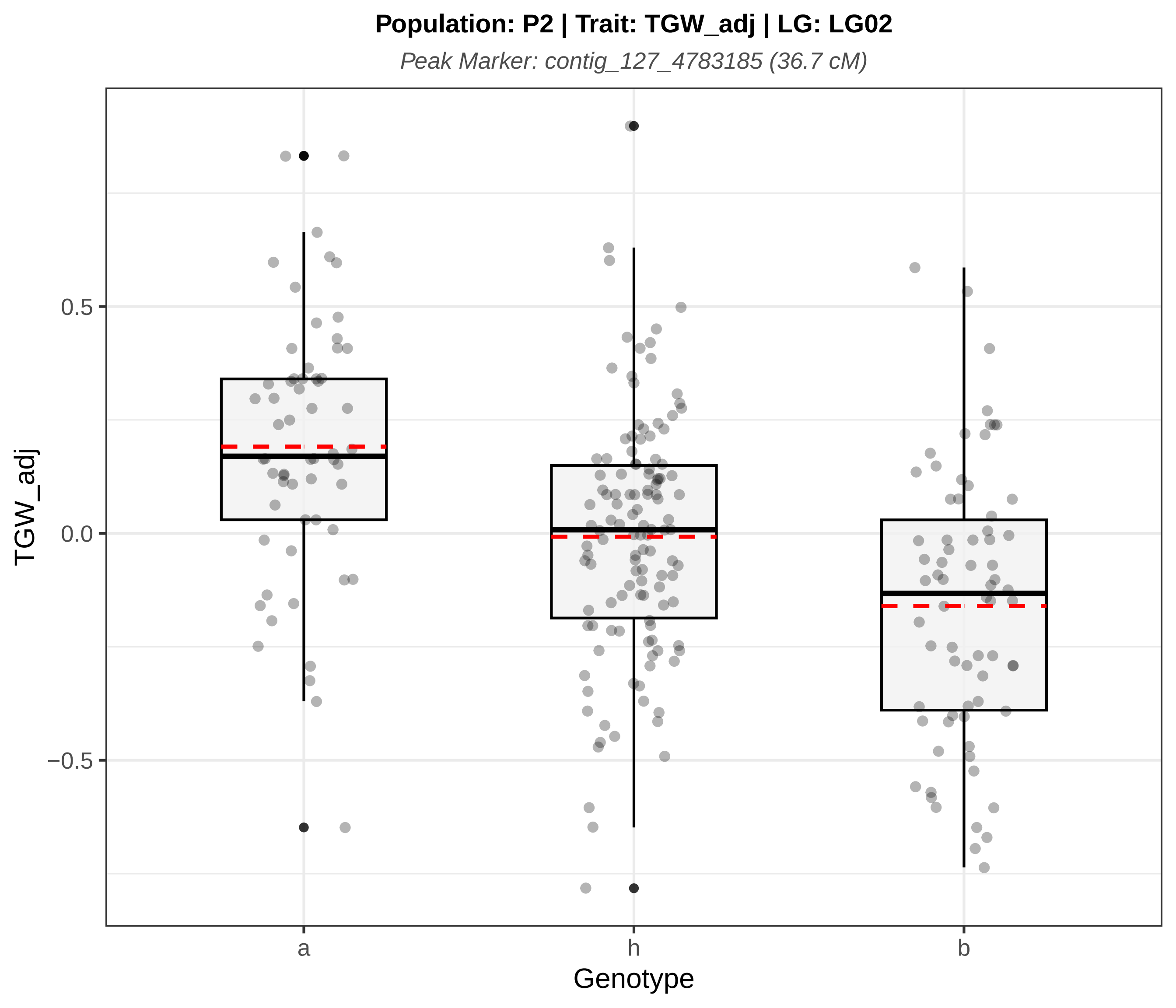
Population: P2 | Trait: TGW_adj | LG: LG02
Peak Marker: contig_127_4783185 (36.7 cM)
TGW_adj
0.5
0.0
−0.5
a
h
b
Genotype
