## Supplementary figures and images for "QTL analysis of essential oil, seed shattering, thousand-grain weight and stalk attachment rate in *Carum carvi*"

### Figure S1

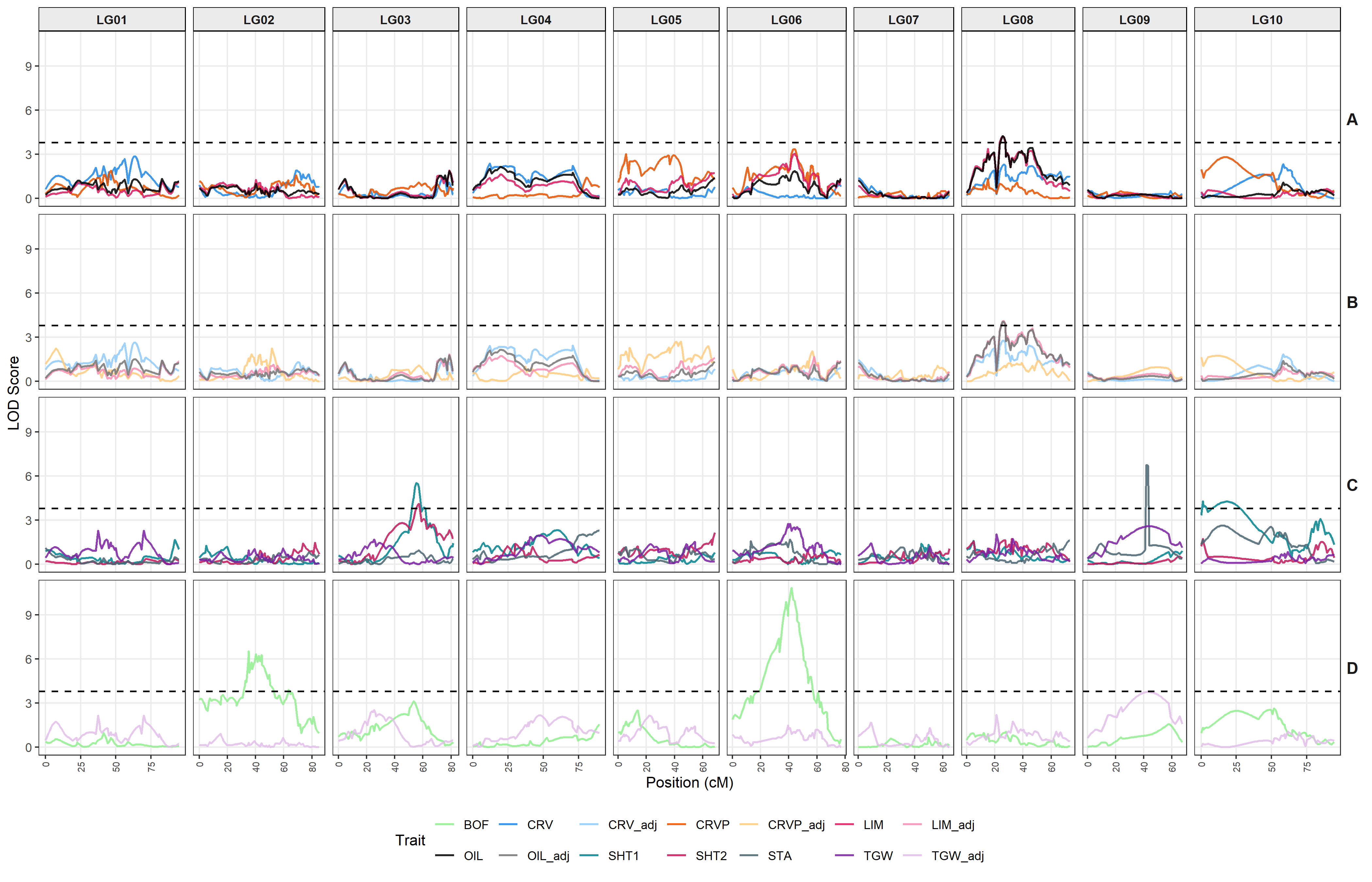

### Figure S2

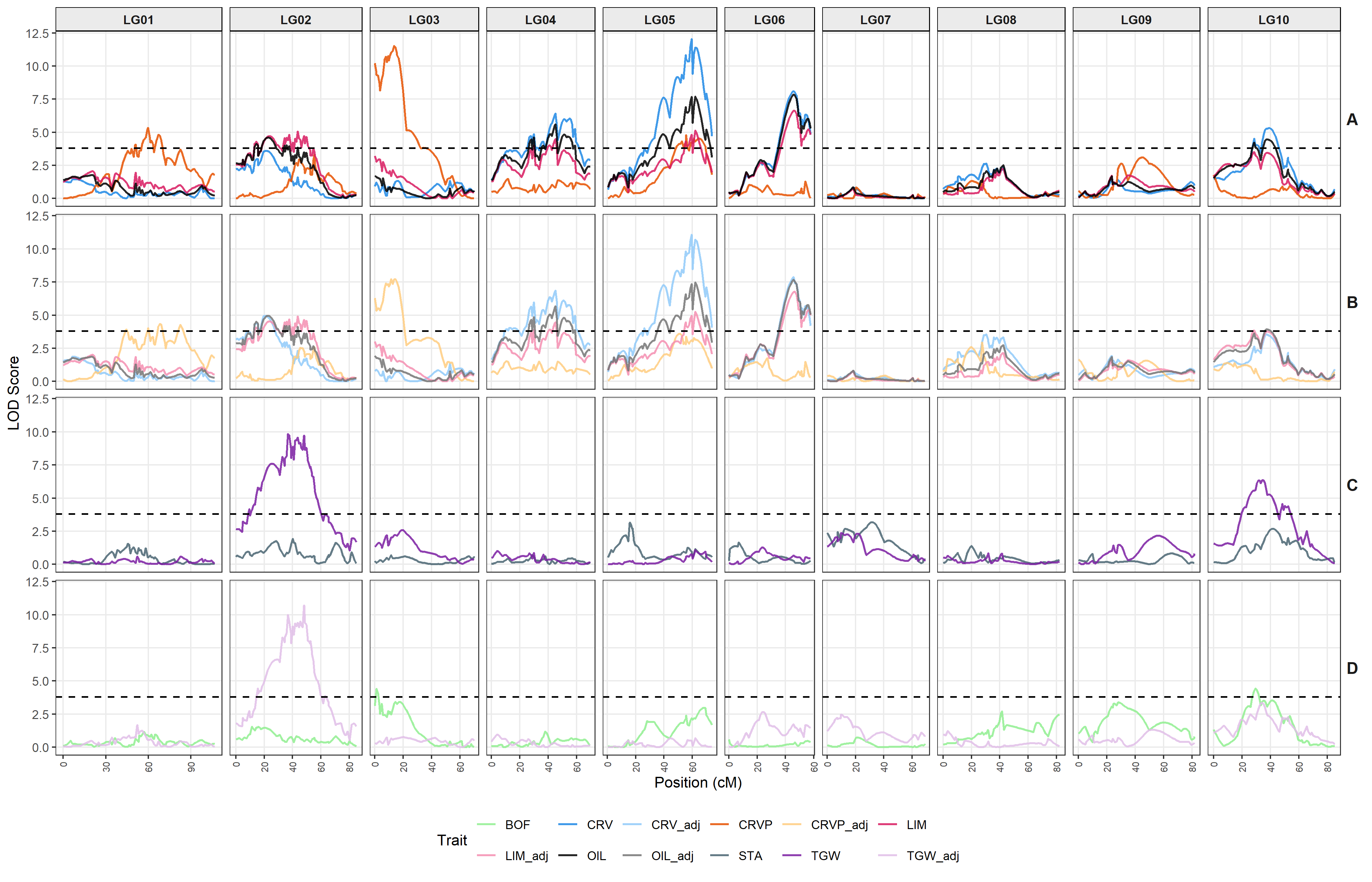

### Figure S3

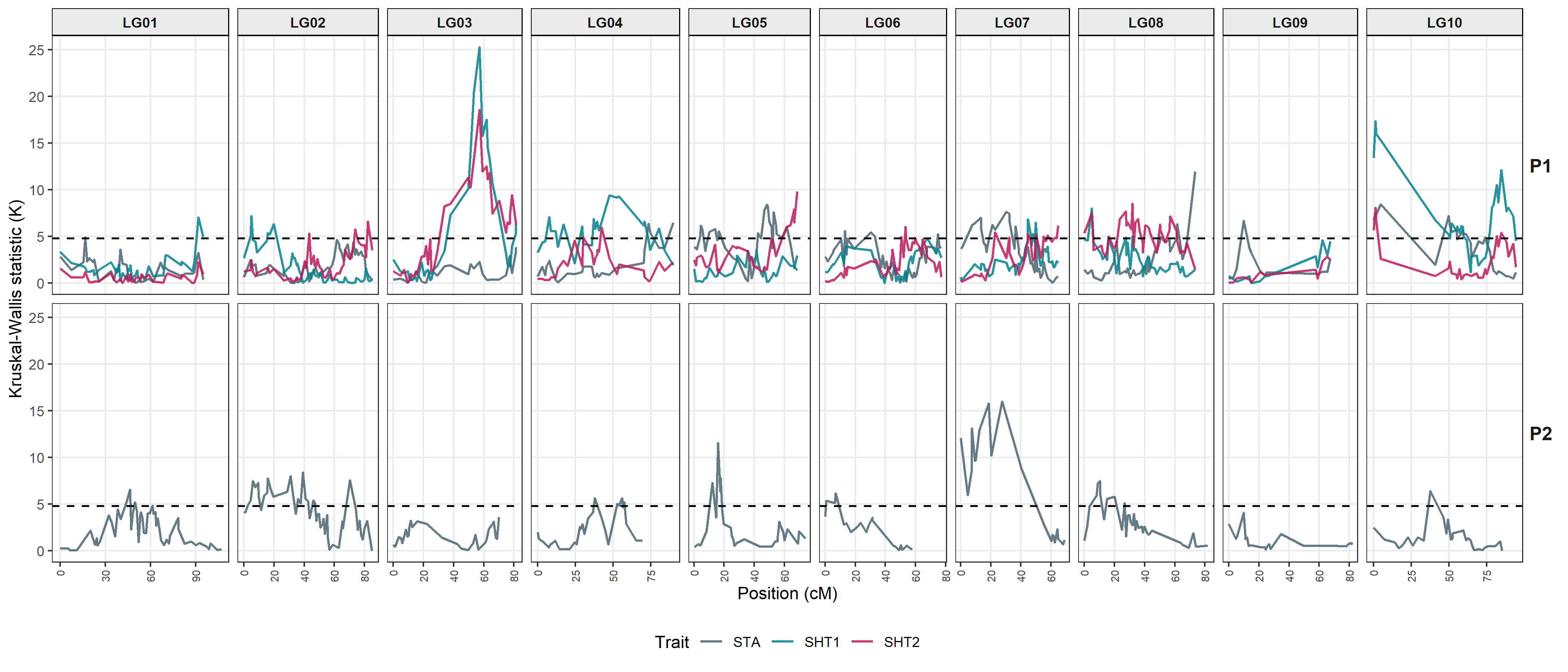
